# Identification of an epigenetically and phenotypically distinct peritumoral glioblastoma cell population linked to inferior patient outcome

**DOI:** 10.64898/2026.01.16.699862

**Authors:** Inês Neves, Xi Lu, Veera Jokinen, Francesco Latini, Nagaprathyusha Maturi, Anders Sundström, Yonglong Dang, Irem Uppman, Tobias Bergström, Pengwei Xing, Mats Ryttlefors, Xingqi Chen, Fredrik J. Swartling, Lene Uhrbom

## Abstract

Glioblastoma (GB) is an aggressive and therapy-resistant primary brain tumor with dismal prognosis. Lethality is in most patients caused by a local peritumoral (Edge) relapse near the resection cavity. This region is insufficiently studied and experimental models are scarce. We have analyzed matched tissue samples and cell cultures from bulk tumors and Edge regions of 11 GB patients. Genomic profiling displayed similar genetic alterations and subclonal distributions of matched bulk and Edge regions. Functional, phenotypic and combined single nucleus (sn) RNA-seq and ATAC-seq analyses showed distinct differences between bulk and Edge GB cells with similar shifts across patients. Sn multiomics uncovered a subset of Edge cells defined by unique chromatin accessibility signatures and elevated mesenchymal and immune gene expression. Integrated computational and functional investigations converged on microglia-derived Oncostatin-M as a driver of the Edge-unique phenotype, and high expression of Edge-related signatures correlated with inferior patient outcome in independent GB cohorts.

## INTRODUCTION

GB is the most frequent and aggressive primary brain cancer and remains incurable. The overall median survival is 6-10 months^1^ in registry databases and 14-21 months in patients treated with standard of care^2,3^. The standard of care has for two decades included maximal-safe surgical resection of contrast-enhanced tissue followed by radiation therapy with concomitant and adjuvant temozolomide^4^. Intra-operative tumor visualization using 5-aminolevulinic acid (5-ALA) fluorescence has increased progression-free survival^5^, by extending the tumor border up to 10 mm beyond the T1 contrast-enhanced region^6^. Despite treatment, tumor relapse is inevitable, with recurrent tumors occurring near the resection cavity in over 90% of patients^7,8,9^. This emphasizes the importance of targeting the therapy-resistant tumor cells residing in the local infiltrative margin of the primary bulk tumor.

GB has been extensively investigated and shows remarkable inter-patient and intra-tumor heterogeneity^9–11^ with dynamic cancer cells that transition between different transcriptome-defined states^12,13^. Cells from primary tumors display a range of non-genetically determined molecular phenotypes^14,15^, and an analysis of paired primary and recurrent bulk tumor samples proposed that GB progression occurs as a consequence of state dynamics rather than through mutation-driven clonal evolution^16^. Importantly, these studies have been performed on tissues and cells of the primary and recurrent bulk tumor, while comparisons between infiltrative primary peritumoral regions with matched recurrences are lacking.

Furthermore, despite strong support of relapse-causing cells residing close to the primary bulk tumor^7,8^ there are few investigations where these have been analyzed in relation to their matched bulk. In those, T1 contrast-enhancement was used to define the tumor border and peritumoral samples were collected from a T2 hyperintense region near the bulk tumor^17–21^. Matched samples were profiled with single cell^17,20,21^ or bulk^18,19^ RNA-seq, and overall spatial differences regarding presence and composition of cell types and cell states of both GB and stromal cells were reported, with for example a more invasion-related phenotype at the migrating front^18^, and expression of gene signatures related to myeloid cells, mesenchymal subtype, wound healing response and glycolytic metabolism in peripheral GB samples^22^.

To extend the understanding and establish validated models of local peritumoral (Edge) GB cells we have collected matched samples from the bulk tumor and an Edge region beyond the 5-ALA fluorescent border of 11 patients. Tumor samples and matched sustainable GB cell cultures (GCCs) were comprehensively characterized through functional, phenotypic, genomic and combined sn multiomic analyses. Genomic profiling showed high similarities between matched bulk and Edge samples whereas functional and sn multiomic analyses identified distinct differences between bulk and Edge with similar shifts across patients. Chromatin accessibility-based subcluster analysis identified a subset of Edge cells that were defined by Edge-unique gene signatures. Integrated computational and functional investigations converged on a microglia-derived factor, Oncostatin-M (OSM), as a driver of these Edge-unique features. Furthermore, both Edge-unique and OSM-activation signatures could predict patient outcome in independent GB cohorts.

## RESULTS

### Local peritumoral GB Edge cells can be maintained in culture and display distinct functional characteristics

Our investigation builds on samples from 11 GB patients undergoing 5-ALA fluorescence-guided surgery (Table S1, Figure S1). Multiple samples were collected from each patient that were used for explantation and range of analyses to uncover the nature of peritumoral GB cells (Fig. 1a). Tissue specimens were resected following a stringent protocol to avoid contamination between bulk and Edge samples (described in Methods). Two samples, Aspirate and Core, were taken from the bulk tumor defined by 5-ALA fluorescence. The Edge sample was collected from a non-fluorescent and non-eloquent (as determined intra-operatively) region at least 10 mm outside of the fluorescent margin. Samples were processed and explanted under stem cell conditions which resulted in sustainable and authenticated (Table S2) matched Aspirate, Core and Edge cultures from six patients. Presence and frequency of tumor cells were analyzed by immunostainings for SOX2, a GB cell marker with rare expression in normal adult brain^23^, in tissue samples (Fig. 1b, Extended Data Fig. 1a) and cell cultures (Fig.1c, Extended Data Fig. 1b).

**Figure 1.**
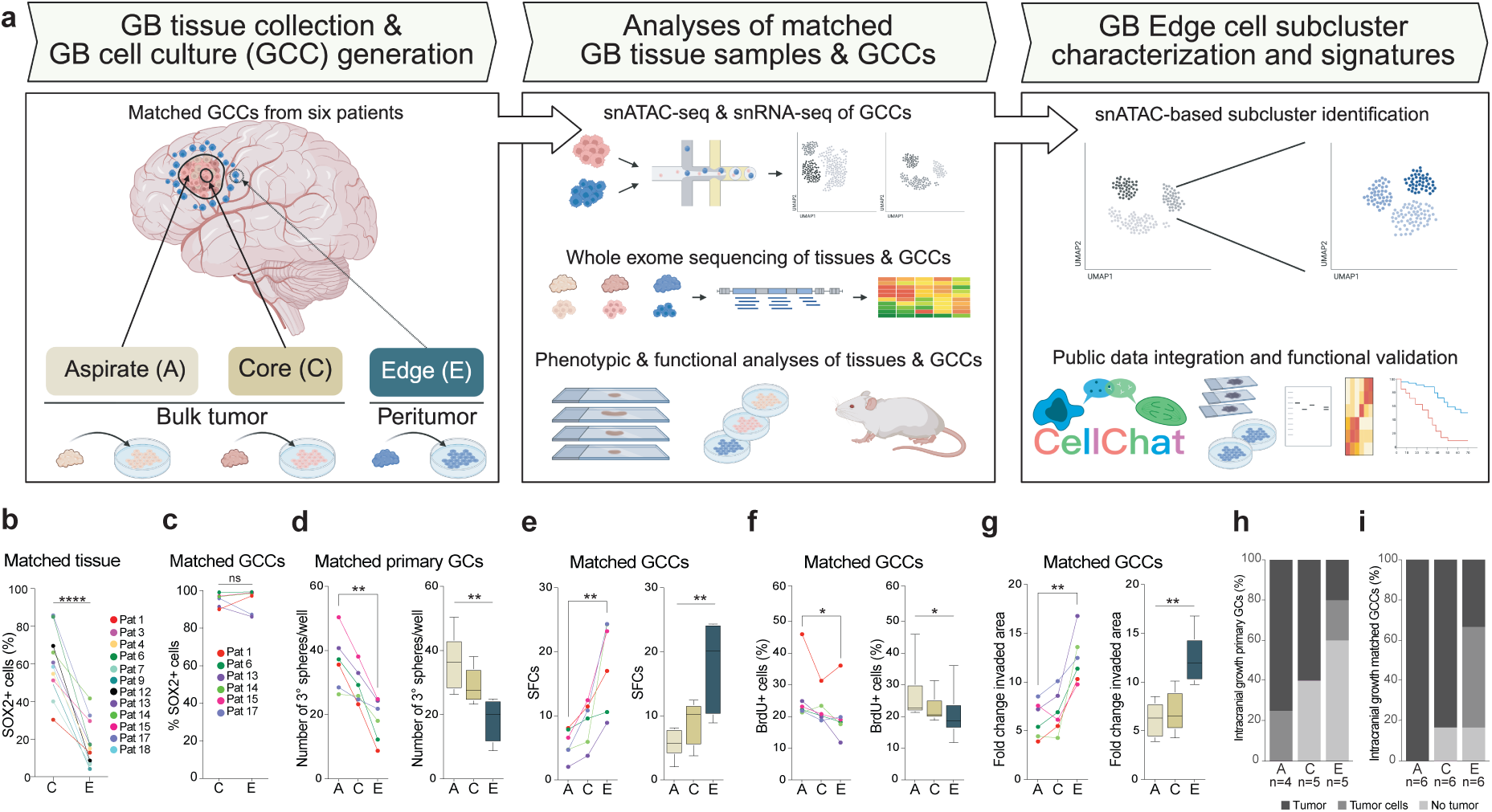
Peritumoral GB Edge cell cultures display distinct functional characteristics compared to their matched bulk tumor cultures. **(a)** Schematic overview of the study. **(b)** Proportion of SOX2^+^ cells relative to all DAPI+ cells in matched Core and Edge tissue samples of 11 patients. Paired t-test Core vs Edge, p<0.0001. **(c-g)** Analyses of matched patient-derived primary or sustainable GB cell cultures of six patients. **(c)** Proportion of SOX2+ cells relative to all DAPI+ cells in matched Core and Edge cell cultures of 6 patients. Paired t-test Core vs. Edge, p=0.1337. **(d-g)** Analyses of matched patient samples. Left, paired t-test Aspirate vs. Edge. Right, RM One-way ANOVA on merged data. Box plots show the distribution’s median, first, and third quartiles, and whiskers represent either the 1.5-times interquartile range or the most extreme value. **(d)** Number of tertiary spheres formed from freshly dissociated matched tissue samples, p=0.0045 (left), p=0.0033 (right). **(e)** Number of sphere forming cells (SFCs) from ELDA based on three biological replicates, p=0.0083 (left), p=0.0080 (right). **(f)** Proportion of BrdU+ cells based on three biological replicates, p=0.0190 (left), p=0.0408 (right). **(g)** Fold-change invasion based on three biological replicates, p=0.0033 (left), p=0.0016 (right). **(h-i)** Tumorigenicity scoring of intracranially injected patient-derived Aspirate, Core and Edge freshly dissociated samples (n=4, n=5, n=5, respectively) (**h**), and matched GCCs (n=6) (**i**). Capacity to form a tumor mass (tumor), latent diffuse tumor cells (tumor cells) or no sign of tumor or tumor cells (no tumor) was determined with an experimental endpoint of 1 year.

Functional investigations of matched bulk and Edge cell cultures focused on key GB cell characteristics. Self-renewal was determined in both acutely dissociated tumor samples using consecutive sphere assays (Fig. 1d) and in established cultures by extreme limiting dilution assays (ELDA) (Fig. 1e). This showed that Aspirate cultures consistently had a higher sphere-forming capacity than their matched Edge culture in both primary cells and established cultures. Proliferation was analyzed through bromodeoxyuridine (BrdU) incorporation and showed significant differences between matched Aspirate and Edge cultures (Fig. 1f). Invasion was examined with the collagen invasion assay on cell aggregates from early passage cultures and showed that Edge cultures had a significantly higher invasive capacity compared to their matched Aspirate cultures (Fig. 1g).

We determined orthotopic tumor-forming ability of primary cells and established cultures through intracranial injections of tumor cells in immune-deficient mice. Brains of injected mice were evaluated through hematoxylin and eosin (H&E) staining and immunostainings for STEM121 and SOX2. Each sample was categorized as Tumor (tumor mass in at least one brain), Tumor cells (no tumor mass but a cluster of tumor cells in at least one brain) or No tumor (negative for Tumor and Tumor cells). Injections of non-matched acutely dissociated tissue samples showed that all types of samples had potential to produce patient-derived orthotopic xenografts (PDOXs) (Fig. 1h, Extended Data Fig. 1c). The lower tumor frequency of Edge samples could be due to a lower fraction of tumor cells in these samples but could also reflect a lower tumorigenic potential of Edge cells. In support of the latter, established Edge cultures also had a lower tumorigenic capacity compared to Core and Aspirate cultures (Fig. 1i, Extended Data Fig. 1d, Table S3).

Taken together, these analyses show that GB cells were readily detected in tissue samples isolated from at least 10 mm beyond the fluorescent tumor border. These Edge tumor cells could be sustained in culture and retained key GB cell properties but were qualitatively different to their matched bulk tumor cultures.

### GB Edge cells exhibit evolutionary early subclones that persist at tumor relapse

Whole exome sequencing (WES) was performed on matched Aspirate, Core and Edge GB tissue samples and GCCs. For five patients, a corresponding blood sample was available and used as a reference (Fig. 2a-b), while one patient was analyzed in tumor-only mode (Extended Data Fig. 2a-b). An Oncoprint visualization of common GB somatic mutations^11^ showed that a mutation found in an Aspirate tissue or cell culture of a patient in nearly all cases were detected in its matched Edge tissue or cell culture (Fig. 2a, Extended Data Fig. 2a). This was also true for copy number alterations (CNAs) (Fig. 2b, Extended Data Fig. 2b). Given the clear and similar functional differences between matched bulk and Edge cell cultures across patients we searched for recurrent Edge-unique mutations and pathways in all somatic variants across patients but were unable to identify any in our cohort.

**Figure 2.**
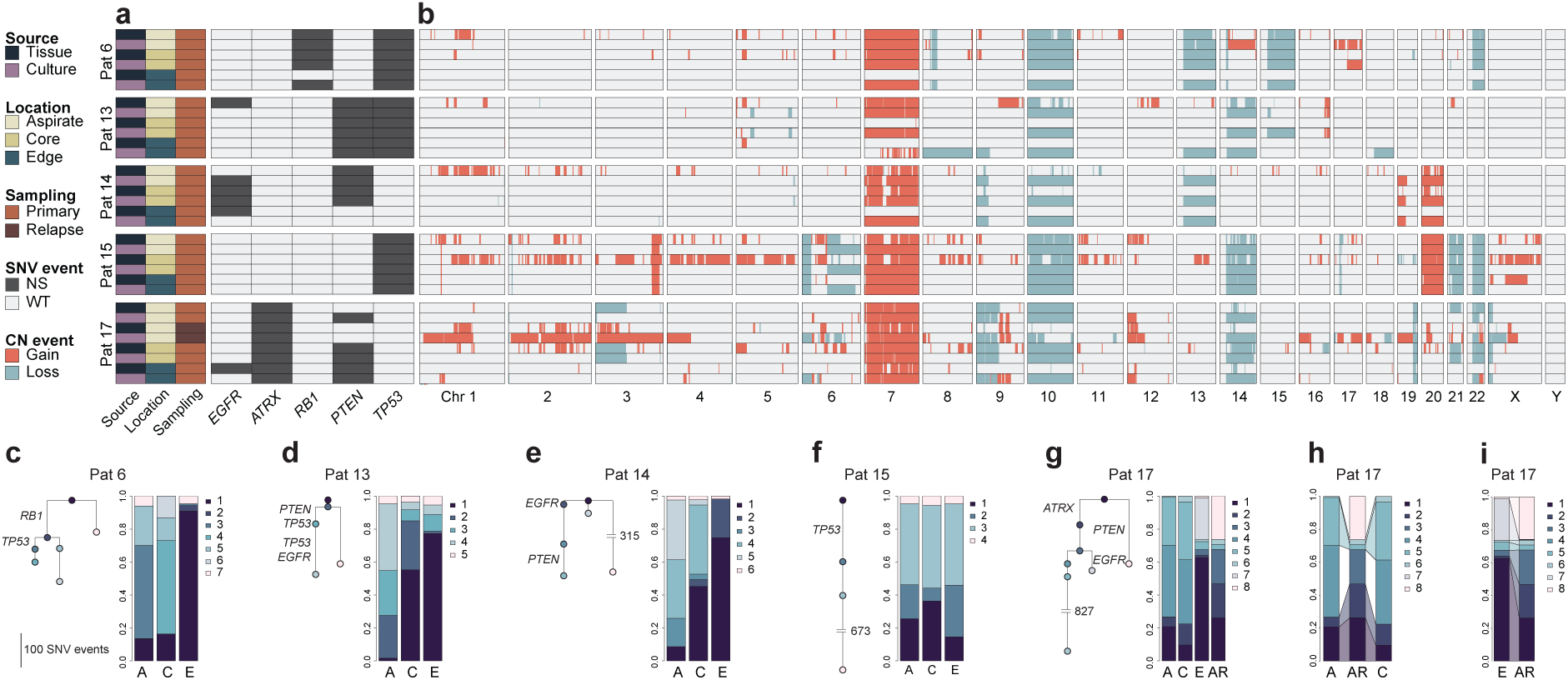
GB Edge samples exhibit common GB alterations and evolutionary early subclones in relation to their matched bulk tumor samples. **(a)** Oncoprint of GB relevant genes from *Wang et al.* in five individual patients across different sample locations. SNV, single nucleotide variant; NS, non-synonymous. **(b)** Copy Number (CN) variation profile of five patients across different sample locations. **(c-g)** Phylogenetic tree inference of mutational relationships in clones of individual patients based on data from Aspirate (A), Core (C) and Edge (E) tumor samples. The branch length is equal to the number of mutations (left). The bar plots represent the composition of the subclonal populations in each sample (right). **(h-i)** Comparisons of subclones across samples of patient 17. Only subclones with a population frequency of at least 1% are shown. (**h**) Primary bulk tumor Aspirate and Core samples compared with their matched relapse sample Aspirate recurrence (AR). (**i**) Primary Edge tumor sample compared with its matched Aspirate Recurrence.

To understand clonal diversity and hierarchies, particularly the relation of Edge tumor cells with their matched bulk tumor, we used all tumor-specific somatic mutations and CNAs of the tissue samples of each patient in a phylogenetic reconstruction analysis (Fig. 2c-g, Table S4). The resulting tree topology and clonal fraction distribution revealed intra-tumoral heterogeneity and evolutionary paths of tumor cells. A cluster of variant allele frequencies (VAFs) adjusted by local copy number status defined a clonal population. Additional mutations with lower VAFs identified subclones and potential branches in the phylogenetic tree with the branch length denoting evolutionary distance. The presence and distribution of subclones showed that all tumor samples of the same patient displayed similar subclonal diversities but with different relative proportions (Fig. 2c-g, Table S4). In general, Edge samples had a higher fraction of evolutionary early subclones, and in all but one patient (patient 14) all Aspirate subclones were present in their matched Edge. Perhaps even more unexpectedly, in only one patient (patient 17) we could find an Edge-unique subclone (clone 8, Table S4, Fig. 2g). Clonal analysis of a relapse-sample of this patient showed that all subclones present in the primary bulk tumor persisted in the relapse sample (Table S4). Comparing subclones of Aspirate and Core with Aspirate relapse (AR) showed that clone 8 was exclusive to AR and that clone 3 was heavily expanded in AR (Table S4, Fig. 2h), while all Edge subclones were present in the AR sample, and vice versa (Table S4, Fig. 2i). This supported previous reports that recurrence is not primarily driven by new mutations^16,24,25^ and emphasized the importance of local peritumoral cells in the relapse process.

In all, the genomic analyses uncovered intra-tumoral subclonal hierarchies proposing that Edge cells populate the tumor periphery early during tumor development. However, the subclonal heterogeneity of Edge samples suggested a continuous addition of bulk tumor cells to the Edge regions.

### Single-cell transcriptomics reveals a greater heterogeneity and enrichment of MES and immune-activated phenotypes in Edge cells

The rarity of Edge-unique genomic subclones implied that tumor progression in local peritumoral regions was non-genomically driven. To investigate mechanisms underlying the functional and phenotypic differences between bulk and peritumoral GB cells, single nucleus (sn) multiomic (snRNA-seq and snATAC-seq) profiling was performed on six early passage matched Core and Edge cell cultures from three patients. After processing 27670 snRNA-seq and 22741 combined snRNA-seq and snATAC-seq cells passed the quality filters (Extended Data Fig. 3).

To assess phenotypic heterogeneity within and across the cell cultures and relate our data to established GB cell states^13,14^, all qualified snRNA-seq profiles were analyzed. All *Neftel* cell states^13^ were present in all cultures (Fig. 3a-c). Comparing matched cultures revealed a clear shift from less (neural precursor cell (NPC), oligodendrocyte precursor cell (OPC)) to more differentiated (AC, MES) states in Core versus Edge cells (Fig. 3a-c) and overall (Fig. 3d). In alignment, *Richards* cell states^14^ showed a transition from developmental to injury response in matched Core versus Edge cells (Fig. 3e-g) and overall (Fig. 3h). To validate these findings, we analyzed the distribution of *Neftel* (Extended Data Fig. 4a-d) and *Richards* (Extended Data Fig. 4e-h) cell states using scRNA-seq tumor cell profiles from matched tissue samples of the tumor Center and Periphery of three GB patients^21^. These samples contained lower numbers of tumor cells, especially Periphery samples. Still, this data exhibited a similar result of less differentiated and more developmental states in Center GB cells and more differentiated and injury response states in Periphery GB cells, in line with the result of our cell cultures.

**Figure 3.**
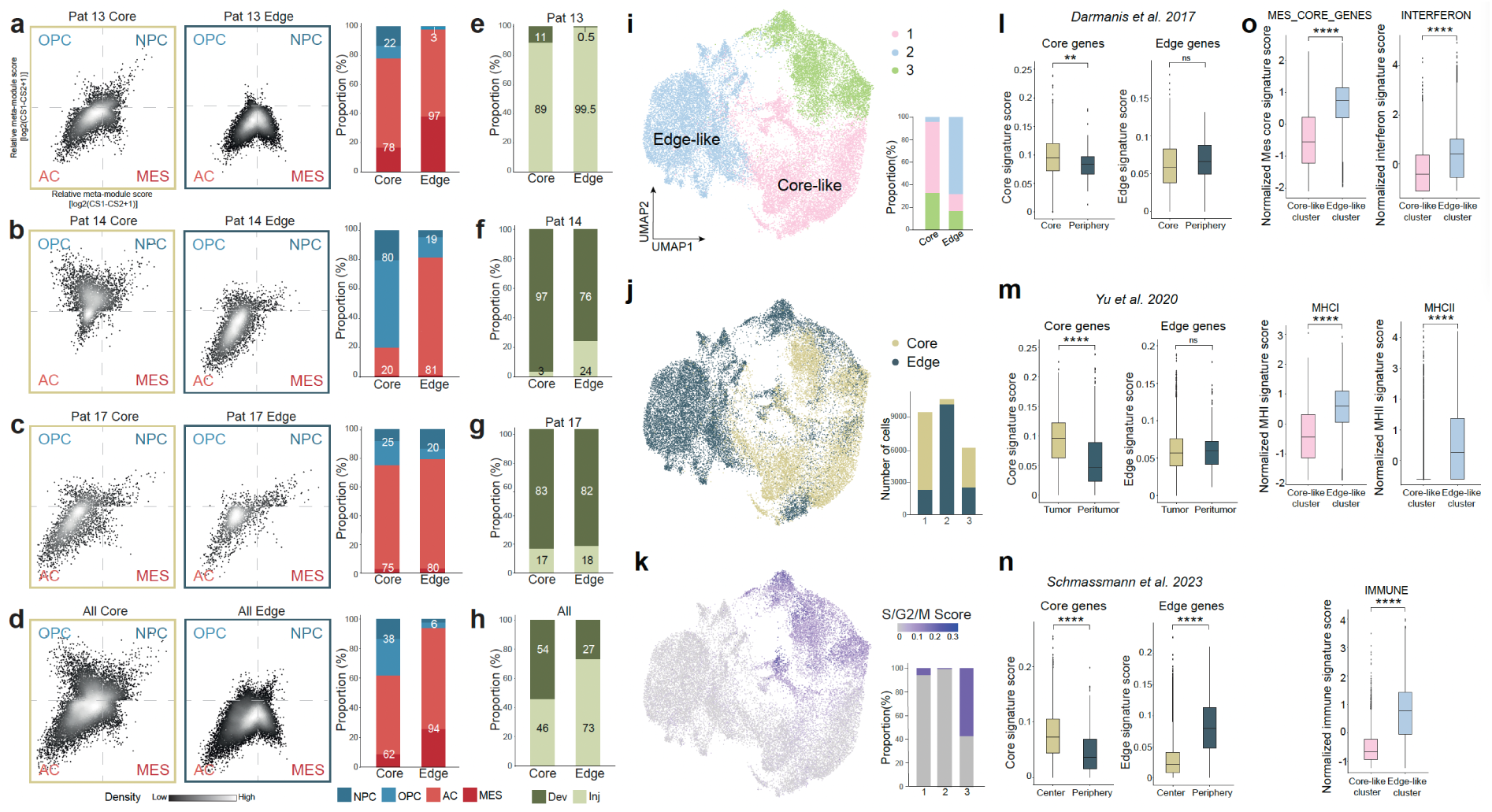
Single-cell transcriptomics reveals enrichment of mesenchymal cell states and a higher heterogeneity of Edge cells. **(a-d)** Visual representation of *Neftel* cell state heterogeneity of Core profiles (left) and Edge profiles (middle), and proportion of combined NPC+OPC and AC+MES cell states (right) in individual patients (**a-c**) and combined data (**d**). **(e-h)** Proportion of *Richards* Developmental (Dev) and Injury response (Inj) cell states in individual patients (**e-g**) and combined data (**h**). **(i)** Uniform manifold approximation and projection (UMAP) representation of major populations from snRNA-seq of all qualified cells (n=27670) (left). Proportion of clusters in Core and Edge cells (right) **(j)** Core and Edge cells projected on the snRNA UMAP (left). Absolute numbers of Core and Edge cells in each cluster (right): 1 (Core, n=7534; Edge, n=2380), 2 (Core, n=547; Edge, n=10709), 3 (Core, n=3904; Edge, n=2596). **(k)** Cycling cells projected on the UMAP (left). Proportion of cycling cells in each cluster (right). **(l-n)** Core and Edge gene signature expression in public scRNA-seq profiles. Statistical significance was assessed with the Mann-Whitney U test. (**l**) *Darmanis* neoplastic cells of Core (p=0.0053) and Periphery samples (p=0.054), (**m**) *Yu* neoplastic cells of the Tumor (p<2.22e-16) and Peritumor samples (p=0.88), and (**n**) *Schmassmann* CD45-negative cells of Center (p<4.72e-67) and Periphery samples (p<3.25e-146). **(o)** Gene scores of *Chanoch-Myers* mesenchymal and immune gene signatures in the Core-like and Edge-like clusters. Statistical significance was assessed with the Mann-Whitney U test, p < 2.22^-16^.

Graph-based clustering of all qualified snRNA-seq profiles was performed in an unbiased approach to understand the differences between Core and Edge GB cells. This identified three clusters (Fig. 3i) where the majority of Core cells were in cluster 1 (Core-like cluster) and Edge cells in cluster 2 (Edge-like cluster) (Fig. 3i-j). Cluster 3 was enriched for cycling cells (Fig. 3k). Thus, Core cells were essentially divided between the Core-like cluster 1 (62%) and the Cycling cluster 3 (33%) with only 5% in the Edge-like cluster 2, while almost equal fractions of Edge cells were found in the Core-like (15%) and Cycling (17%) clusters with the rest in the Edge-like cluster (68%). This corroborated the interpretation of the phylogeny results (Fig. 2c-g) suggesting that the major transfer of tumor cells would be in the direction of bulk to Edge regions.

Differential gene expression (DGE) analysis between all cells of the Core-like cluster 1 and the Edge-like cluster 2 identified a Core-Edge gene signature of 49 Core genes and 73 Edge genes (Table S5). This signature was applied on five publicly available transcriptome data sets of spatially separated GB tissue samples. In the *Darmanis*^17^, *Yu*^20^ and *Schmassmann*^21^ scRNA-seq data, expression of the Core and Edge genes were analyzed separately for each spatial sample. Core genes could in all cases separate bulk tumor cells from peripheral tumor cells (Fig. 3l-n, left), while Edge genes showed a trend towards higher expression in peripheral tumor cells with significance in one data set (Fig. 3l-n, right). The bulk RNA-seq data^18,19^ were analyzed with non-negative matrix factorization (NMF) clustering using the Core-Edge gene signature which with high precision could stratify bulk tumor from peripheral tumor samples (Extended Data Fig. 4i-j). The fact that Core genes could more precisely separate bulk and peritumoral GB cells aligned with the finding that Core cells were more homogenous than Edge cells.

Gene set enrichment analysis (GSEA) of the Core and Edge genes showed an upregulation of RAS, MYC and stemness markers in the Core-like cluster and immune-related and mesenchymal pathways in the Edge-like cluster (Extended Data Fig. 4k). This was in line with the cell state analyses of Core and Edge cells (Fig. 3a-d) and was further supported by analysis of additional mesenchymal cell state gene signatures^26^ comparing expression in the Core-like and Edge-like clusters (Fig. 3o).

### Single-cell epigenomics identifies Edge-unique subclusters

Mechanisms shaping the differential phenotypes of Core and Edge cells were explored by analyzing the multiome data, i.e. nuclei with dual snRNA-seq and snATAC-seq profiles (n=22,741). Cluster analysis of this subset of snRNA-seq profiles (Extended Data Fig. 5a-c) recapitulated previous clusters (Fig. 3i-k). This result confirmed that Core cells predominantly belonged to either the Core-like (71%) or Cycling (24%) clusters with only 5% in the Edge-like cluster, while Edge cells were divided with 64% in the Edge-like, 23% in the Cycling and 13% in the Core-like cluster, respectively (Extended Data Fig. 5b). Clustering of the corresponding snATAC-seq data identified two clusters, A1 and A2 (Fig. 4a). The distribution of Core and Edge cells within these clusters revealed that almost all Core cells (98%) belonged to cluster A1 (Fig. 4b), which also included the cycling cells (Fig. 4c), and that Edge cells were divided between both clusters with 60% in cluster A2 (Fig. 4b), again pointing to a higher heterogeneity of Edge cells displaying a range of Core to Edge phenotypes.

**Figure 4.**
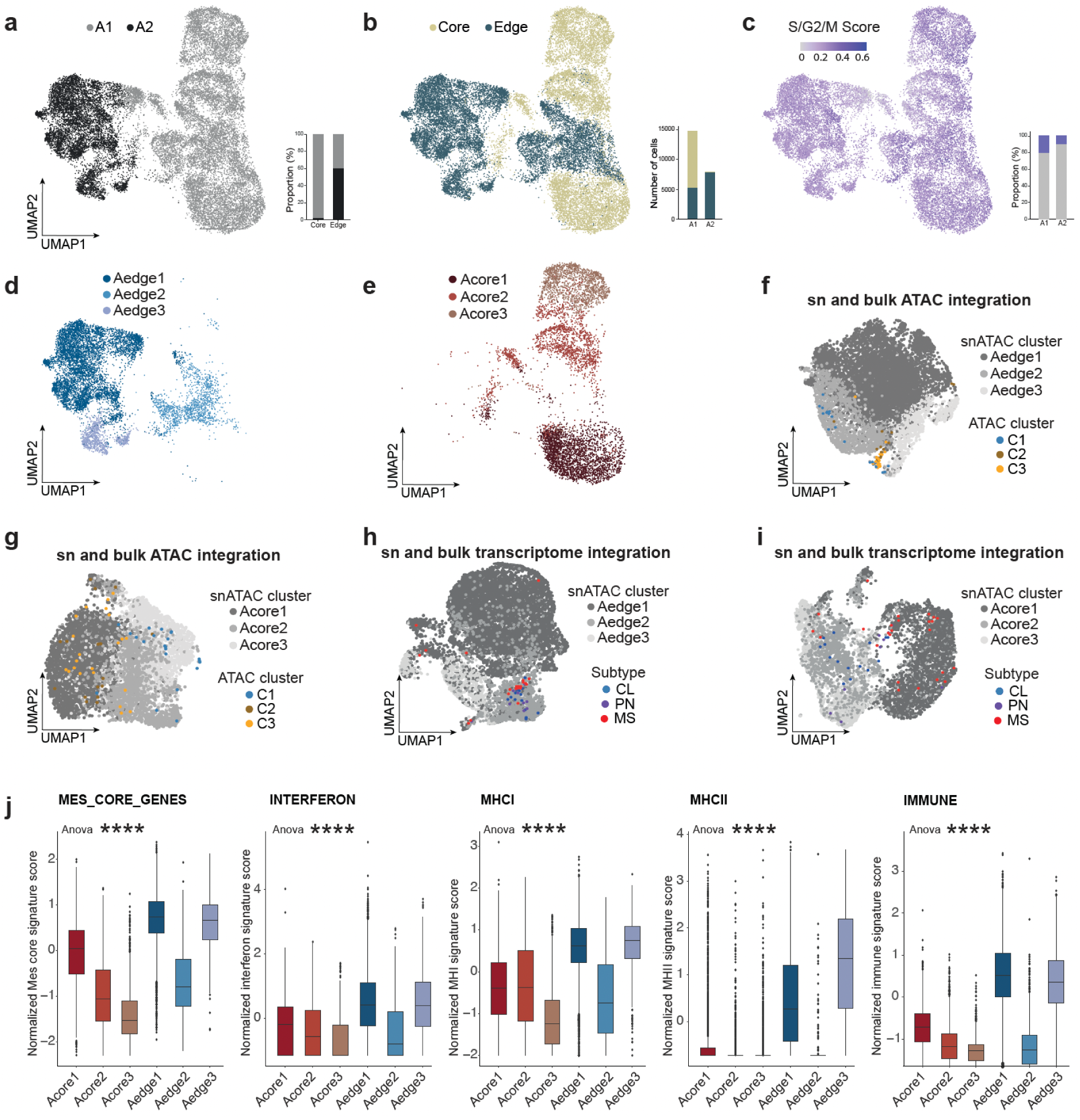
Single-cell epigenomics identifies two Edge-unique subclusters. **(a)** UMAP representation of major populations A1 (n=14811) and A2 (n=7930) of snATAC-seq profiles from nuclei qualified for both snRNA-seq and snATAC-seq (n=22741). The proportion of clusters in Core and Edge cells (right). **(b)** Core and Edge cells projected on the snATAC UMAP clusters (left). Absolute numbers of Core and Edge cells in each cluster (right): A1 (Core, n=9658; Edge, n=5153), A2 (Core, n=166; Edge, n=7764). **(c)** Projection of cycling cells on the snATAC clusters (left). Proportion of cycling cells in each cluster (right). **(d-e)** UMAP representation of subcluster analysis of (**d**) Edge profiles (n=8998) of the Edge-like cluster R2 into Aedge1, Aedge2, and Aedge3, and (**e**) Core profiles (n=6126) of the Core-like cluster R1 into Acore1, Acore2, and Acore3. **(f-g)** Integration of snATAC-seq profiles partitioned in pseudobulk and denoted by their subcluster assignment with *Lu* ATAC-seq data of 60 GCCs derived from bulk tumor samples denoted with their ATAC cluster assignment (C1, C2, C3). GCC ATAC profiles are projected on snATAC Edge pseudobulk profiles (**f**), and on snATAC Core pseudobulk profiles (**g**). **(h-i)** Integration of snRNA-seq profiles partitioned in pseudobulk and denoted by their subcluster assignment with *Lu* transcriptome data of 50 GCCs derived from bulk tumor samples denoted with their TCGA subtype assignment (classical (CL), proneural (PN), mesenchymal (MS)). GCC transriptome profiles are projected on snRNA Edge pseudobulk profiles (**h**), and on snRNA Core pseudobulk profiles (**i**). **(j)** Expression of *Chanoch-Myers* mesenchymal and immune gene signatures in the Core and Edge subclusters. Statistical significance was assessed using analysis of variance (One-way ANOVA) with p<0.0001.

The regulatory landscapes underlying the Core-like R1 and Edge-like R2 clusters were analyzed using the corresponding snATAC-seq profiles. Unique ATAC peaks associated with each cluster were identified (Table S6), and transcription factor (TF) motif enrichment analysis of differential peaks revealed distinct regulatory signatures between the two cell populations (Table S7). The Core-like cluster R1 was characterized by motifs of basic helix-loop-helix (bHLH) (e.g. OLIG1, OLIG2, NEUROD1, ASCL1), homeodomain (e.g. POU3F2, POU5F1, HOXA1), and high-mobility group (e.g. SOX2, SOX4, SOX10) TFs associated with stem-like, tumor-propagating GB cells^23^. The Edge-like cluster R2 exhibited enrichment for motifs linked to the p53 family (e.g. TP53, TP63, TP73), TEA domain factors (e.g. TEAD1-4), runt domain proteins (e.g. RUNX2, RUNX3), Rel homology domain-containing factors (e.g. NFKB1, NFKB2, REL) and interferon regulatory factors (e.g. IRF2, IRF4, IRF8), implicated in cell invasion, stress response, inflammation, and immune activation. These findings were consistent with the results of functional and transcriptomic analyses and supported that the distinct characteristics of Core and Edge cells were largely driven by epigenetic mechanisms. To further investigate the underlying basis of the phenotypic heterogeneity of Edge cells we performed a subcluster analysis selecting cells based on transcriptome clustering (Extended Data Fig. 5a). Edge profiles of the Edge-like cluster R2 (n=8998) were analyzed and for reference the same analysis was performed on Core profiles of the Core-like cluster R1 (n=6126) (Extended Data Fig. 5b). Analyses of the snRNA profiles produced no meaningful subclusters, while re-clustering the snATAC profiles produced three subclusters for both Edge (Fig. 4d, Aedge1-3) and Core (Fig. 4e, Acore1-3). The subclusters were related to our previously identified and thoroughly characterized patient-derived GB cell cultures stratified based on ATAC-seq^27^. Integration of ATAC-seq data of 60 GCCs (derived from patient bulk tumor samples) with the snATAC data was performed by partitioning the snATAC profiles into pseudo bulk of 50 cells. ATAC-seq profiles labeled by their ATAC cluster (C1, C2, C3) were projected onto the snATAC-seq pseudo bulk data denoted by their subcluster (Fig. 4f-g). For Edge, this showed that the ATAC-seq profiles (C1-C3) essentially only overlapped with the Aedge2 subcluster (Fig. 4f), while there was a wider overlap with the Acore subclusters (Fig. 4g). Differential ATAC peak analysis of snATAC subclusters (Extended Data Fig. 6a) followed by TF motif enrichment analysis of differential peaks (Exteded Data Fig. 6b-d) uncovered for Core cells genes and TFs previously connected with GB bulk tumors^28–33^. The same analyses of Edge subclusters showed that Aedge2 displayed typical GB bulk tumor genes and TF motifs whereas Aedge1 and Aedge3 showed novel GB cell phenotypes (Extended Data Fig. 6e-h). Aedge1 was enriched for interferon regulatory factor motifs (e.g. IRF1, IRF4, IRF7, IRF8), which can be associated with macrophage-mediated tumor regulation^34,35^. Aedge3 showed enrichment of forkhead box family motifs, of which many have been linked to stress adaptation^36,37^ and cancer^36,38^.

Additional characterization of the snATAC subclusters was performed using the snRNA profiles. Similarly to the integration of bulk and sn ATAC data, transcriptome data of 50 GCCs with previously assigned TCGA subtype^39^ was integrated with snRNA-seq pseudo bulk data of Edge (Fig. 4h) or Core (Fig. 4i) cells. Also here, the projection of GCC data on the Edge subclusters was centered over the Aedge2 subcluster, while there was a broader distribution over the Core subcluster landscape. The subcluster phenotypes were further analyzed by DGE analysis of Core (Table S8) and Edge (Table S9) subclusters, followed by gene ontology (GO) analysis. Core subclusters showed enrichment of neuronal development and cell adhesion in Acore1, and nervous system development in Acore3 (Extended Data Fig. 6i). The Edge subcluster Aedge1 was enriched for extracellular matrix structural constituents and Aedge3 for several immune activity terms (Extended Data Fig. 6j). Cell state analyses showed, in line with previous data (Figure 3a-h), that the Core subclusters were more enriched for less differentiated and more developmental states (Extended Data Fig. 6k) compared with the Edge subclusters (Extended Data Fig. 6j). The most Core-like Aedge2 subcluster also had the highest enrichment of immature cell states. Lastly we analyzed expression of the mesenchymal cell state signatures^26^ (Fig. 4j) which convincingly showed that Aedge1 and Aedge3 cells had a significantly different phenotype than the bulk-like Acore1-3 and Aegde2 subclusters.

Taken together, reclustering snATAC profiles of Core and Edge cells identified three subclusters in each analysis of which all Core and one Edge aligned with previously described GB bulk tumor phenotypes^10,13,27^. The Aedge1 and Aedge3 subclusters did not associate with previous molecular classifications and were characterized by highly elevated mesenchymal-and immune-related gene expression proposing that they had developed through epigenetic reprogramming.

### OSM-activation is a potential mechanism shaping the Edge-unique phenotype

To investigate possible underlying mechanisms shaping the distinct Edge cell phenotype we turned to the *Schmassmann* scRNA-seq data of tissue samples collected from the tumor Center and Periphery of three GB patients^21^. CellChat^40^ was applied to analyze potential cell-cell communication patterns in CD45-tumor cells and CD45+ immune cells of Center and Periphery samples separately (Extended Data Fig. 7a). This identified vascular endothelial growth factor (VEGF), Oncostatin-M (OSM) and platelet derived growth factor (PDGF) as unique to Periphery. Analysis of outgoing signaling in Periphery samples showed that OSM was the only truly paracrine factor, produced by macrophages, microglia, monocytes and neutrophils (Extended Data Fig. 7b).

Next we wished to analyze putative communication of our Core and Edge cells with the microenvironment by integrating our snRNA-seq Core and Edge profiles with the *Schmassmann* Center and Periphery CD45+ profiles, respectively. Before doing so we combined the tumor cell profiles of both cohorts in an UMAP analysis which showed a high overlap of Core with Center, and Edge with Periphery GB cells (Fig. 5a), supporting the applicability of an integrated analysis. Separate CellChat analyses were performed for Core with CD45+ Center (Fig. 5b) and Edge with CD45+ Periphery (Fig. 5c) and resulting interactions were curated using the *Schmassmann* only results (Extended Data Fig. 7a) as a help to highight the most tissue-relevant interactions. This identified OSM as the only exclusively paracrine incoming signal in Edge cells, and more specifically in the Egde-unique subclusters Aedge1 and Aedge3.

**Figure 5.**
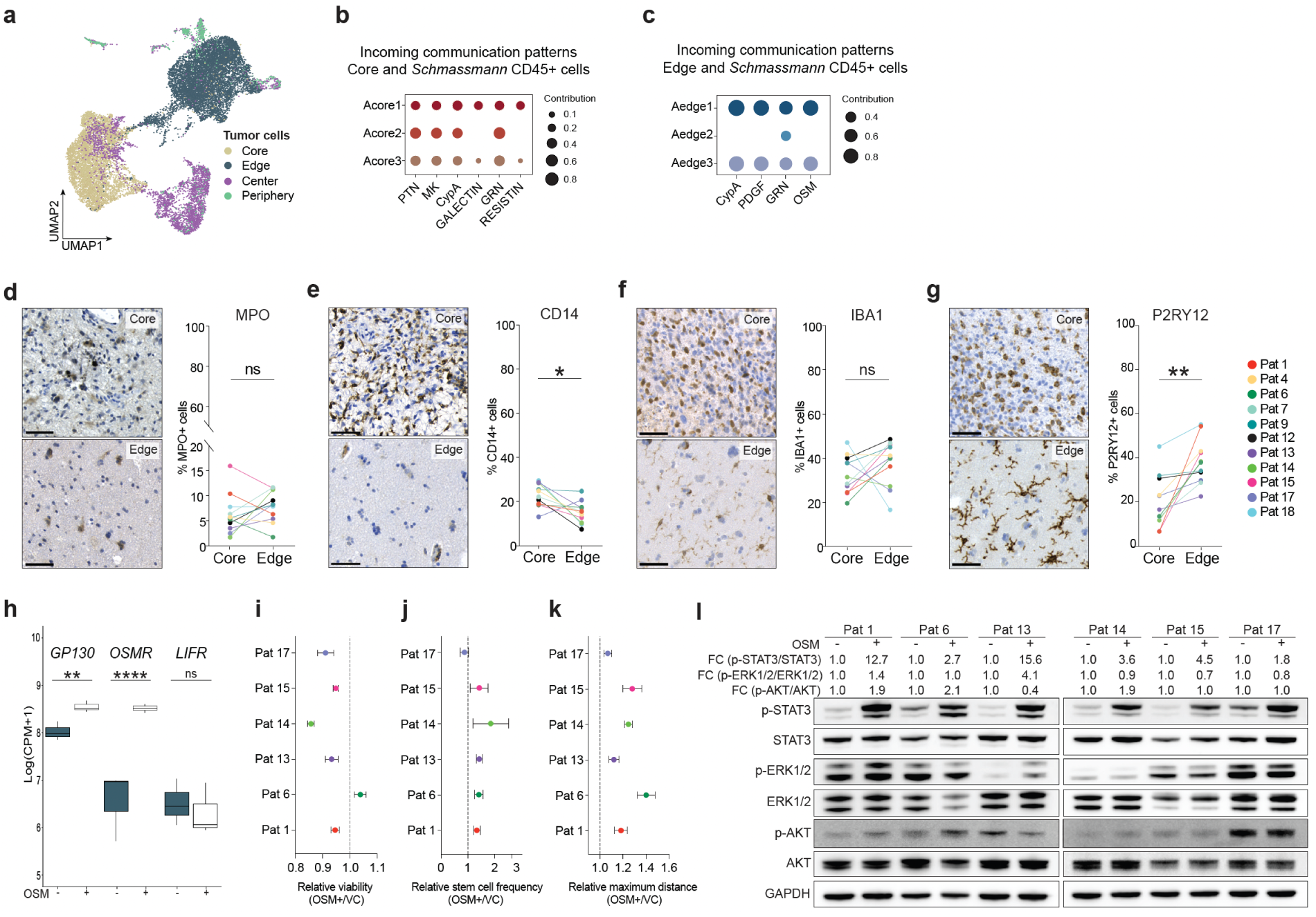
CellChat identifies a potential immune to GB cell communication via OSM that sustains the distinct Edge cell phenotype. **(a)** UMAP representation of the integration of snRNA profiles of Core and Edge cells with scRNA profiles of *Schmassmann* CD45- Center (n=3740) and Periphery (n=628) tumor cells. **(b)** CellChat incoming signaling pattern integrating snRNA profiles of Core cells with *Schmassmann* CD45+ Center cells. **(c)** CellChat incoming signaling pattern integrating snRNA profiles of Edge cells with *Schmassmann* CD45+ Periphery cells. **(d-g)** Representative images of immunostainings of Core and Edge tissue samples of patient 18 (left, scale bar 50 µM), and proportions of marker+ cells related to all cells (hematoxylin+) in 11 GB patients (right, paired t-test). **(d)** MPO, p=0.3036 **(e)** CD14, p=0.0109 **(f)** IBA1, p=0.3364 **(g)** P2RY12, p=0.0022 **(h)** Gene expression of OSM receptors in Edge cells of patient 13, 15 and 17 cultured in presence or absence of OSM. DESeq2 Wald test was performed: GP130, p=0.004; OSMR, p=5.96e-07; LIFR, p=0.70. **(i-k)** Functional consequences of 7 days OSM stimulation in Edge cells normalized to vehicle control (VC). **(i)** Relative viability shown as mean ± SEM of three biological replicates. Paired t-test, p=0.0689. **(j)** Relative stem cell frequency shown as mean ± SEM of three biological replicates. Paired t-test, VC vs OSM, *p=0.0255. **(k)** Relative maximum invasion distance in collagen at 24 hours shown as mean ± SEM of three biological replicates. Paired t-test, VC vs OSM, **p=0.0049. **(l)** Western blot analysis of STAT3, ERK and AKT signaling in Edge cells cultured in absence or presence of OSM.

OSM is mainly produced by macrophages and microglia in the human central nervous system^41,42^. To analyze the frequency of potential OSM-producing cell types in our cohort we performed immunostainings of matched Core and Edge tissue samples of all patients for myeoloperoxidase (MPO, neutrophils), CD14 (monocytes and macrophages), ionized calcium-binding adapter molecule 1 (IBA-1, macrophages and microglia) and purinergic receptor P2Y12 (P2RY12, microglia). There was no difference between Core and Edge for MPO (Fig. 5d), a decrease in Edge for CD14 (Fig. 5e), no difference for IBA-1 (Fig. 5f), and an increase in Edge of P2RY12 positive cells (Fig. 5g). Collectively this pointed to a significantly higher fraction of microglia in Edge samples, in line with the *Schmassmann* data of Center and Periphery samples (Extended Data Fig. 7j) and other investigations of peritumoral GB regions^43–45^.

To investigate the responsiveness of GB Edge cells to OSM, three cultures were subjected to continuous OSM stimulation or vehicle control (VC) for 7 days after which RNA sequencing was performed. OSM signaling can occur via two receptor complexes, where glycoprotein 130 (GP130) is an obligatory partner which, together with either the leukemia inhibitory factor receptor (LIFR, type 1 receptor complex) or the OSM receptor (OSMR, type 2 receptor complex)^46^, bind to OSM and activate downstream signaling pathways, including STAT3, ERK and PI3K, depending on cell type and context^47,48,49–52^. Analysis of RNA-seq data showed that *GP130* was expressed in VC cells and further induced upon OSM stimulation. Both *OSMR* and *LIFR* were lowly expressed in VC cells, where *OSMR* was strongly induced while *LIFR* remained low after 7 days of OSM stimulation (Fig. 5h). This proposed that OSM signaling occurred via the type 2 receptor complex in GB Edge cells. Next, all six Edge cultures were subjected to OSM treatment and cell viability measured which had little effect related to VC control (Fig. 5i), while self-renewal (Fig. 5j and invasion were significantly enhanced by OSM stimulation (Fig. 5k). Western blot analyses uncovered a clear activation of STAT3 signaling with ERK and AKT pathways essentially unaffected (Fig. 5l).

The OSMR-STAT3 signaling appeared to have little effect on the Edge molecular phenotype, with small changes in expression of *Neftel* cell states^13^ (Extended Data Fig. 7d), mesenchymal cell states^26^ (Extended Data Fig. 7e) and the Edge-like genes that had been derived from all profiles of the Edge-like and Core-like cluster (Extended Data Fig 7f). To generate a more defined Edge gene signature DEGs were extracted of Edge profiles of the Edge-like cluster and Core profiles of the Core-like cluster (Table S10). These genes were highly expressed in the Edge-unique subclusters (Aedge1 and Aedge3) (Extended Data Fig. 7g), validating the signature as Edge-unique. Nor this signature could distinguish OSM-activated from non-activated GB Edge cells (Extended Data Fig. 7h). This showed that the Edge cells were primed to functionally respond to OSM signaling which induced minor changes in Edge-defining genes, proposing that the Edge cell phenotype had remained stable under culture conditions.

### Edge-related signatures can predict patient survival

The majority of GB relapses are local and in contact with the resection cavity of the primary tumor^7–9^. Our paired primary and recurrent genomic data showed that all subclones detected in the local peritumoral primary sample constituted all subclones detected in the recurrent tumor sample (Fig. 2g-i), demonstrating the importance of local peritumoral cells in the relapse process. To explore the relation between the GB Edge cell phenotype and patient outcome we analyzed the distribution of Edge cell subclusters in each patient (Fig. 6a) and related the frequency of Edge-unique cells (Aedge1 and Aedge3) to patient survival (Table S1). Acknowledging the small sample size this showed a strong (R=-0.9243) inverse relation (Fig. 6b). There is very limited profiling data of GB peritumoral regions with connected survival data. Therefore, to investigate the relation of the Edge cell phenotype and patient outcome we performed survival analyses on three publicly available GB cohorts^25,53,54^, including only IDH wildtype patients that were stratified at median expression of the Edge-unique signature (Table S11). In the TCGA cohort^53^ (primary bulk GB samples, n=140) this could predict progression-free interval (PFI) (Fig. 6c) but not overall survival (OS) (Fig. 6d), indicating that a higher frequency of Edge-like tumor cells in the bulk tumor could influence therapy response. The other two cohorts had profiling data of primary and recurrent GB tumors^25,54^. In the GLASS cohort^25^ the Edge-unique signature could predict survival in both primary (Fig. 6e, n=163) and recurrent tumor data (Fig. 6f, n=93), with higher prediction value for recurrent tumors, in line with our correlation analysis (Fig. 6b). The *Spitzer* data^54^, however, showed no significance for neither primary tumors (n=121) (Fig. 6g) nor recurrences (n=59) (Fig. 6h), although a separation of the curves in the same direction as for the TCGA and GLASS cohorts could be noted.

**Figure 6.**
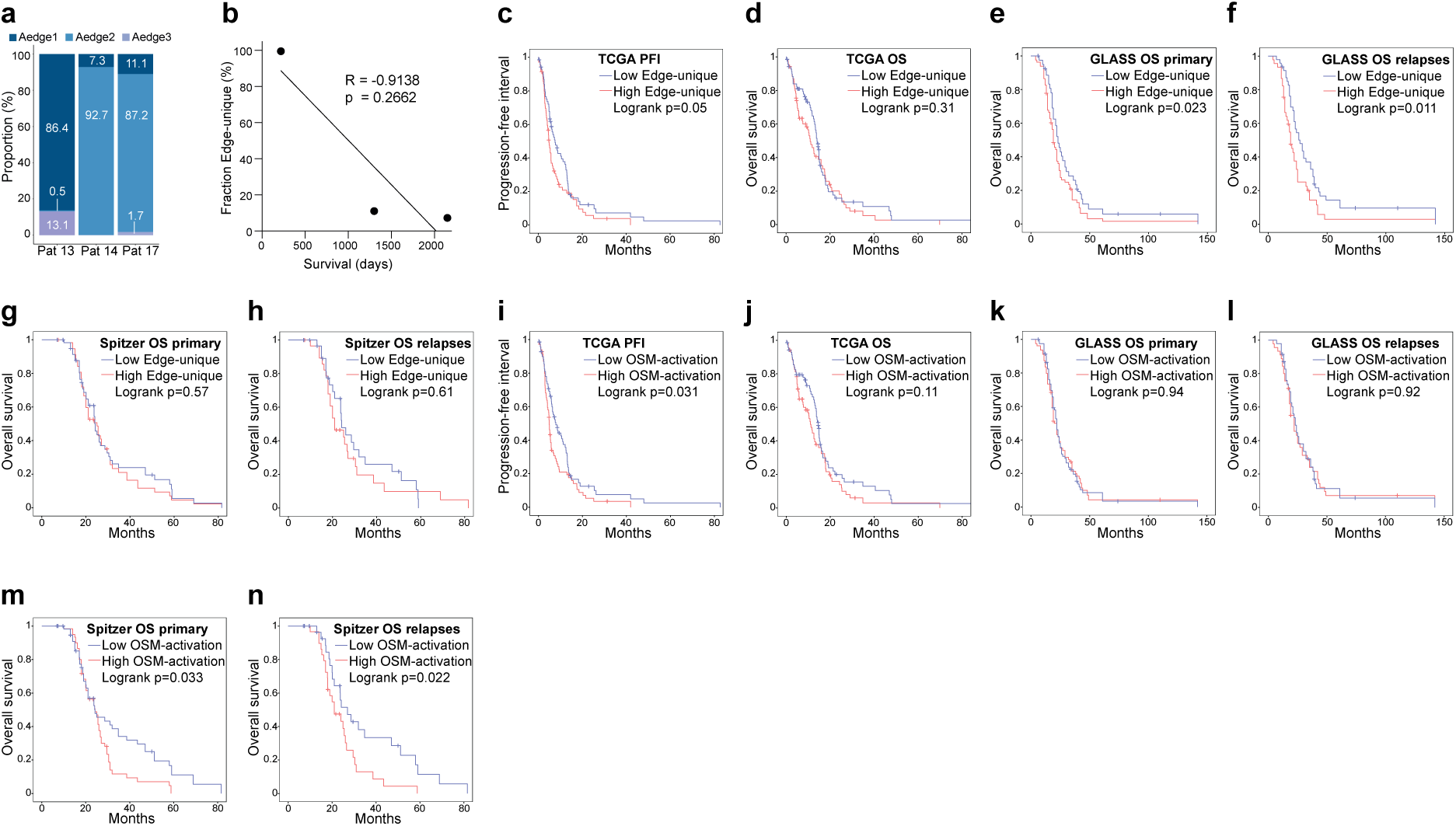
Edge-unique and OSM-activation signatures can predict survival in independent IDH wildtype GB patient cohorts. **(a)** Proportions of Edge subcluster cells (Aedge1-3) in early passage cultures of patient 13, 14 and 17. **(b)** Correlation analysis of fraction of Edge-unique cells (Aedge1 and Aedge3) and patient survival. **(c-n)** Kaplan-Meier analyses of progression-free interval (PFI) or overall survival (OS) of IDH wildtype GB patients of independent cohorts. Log-rank (Mantel-Cox) test was used to calculate p-values. **(c)** PFI of TCGA patients stratified by the Edge-unique signature. **(d)** OS of TCGA patients stratified by the Edge-unique signature. **(e)** OS of GLASS patients (primary tumors) stratified by the Edge-unique signature. **(f)** OS of GLASS patients (recurrent tumors) stratified by the Edge-unique signature. **(g)** OS of *Spitzer* patients (primary tumors) stratified by the Edge-unique signature. **(h)** OS of *Spitzer* patients (recurrent tumors) stratified by the Edge-unique signature. **(i)** PFI of TCGA patients stratified by the OSM-activation signature. **(j)** OS of TCGA patients stratified by the OSM-activation signature. **(k)** OS of GLASS patients (primary tumors) stratified by the OSM-activation signature. **(l)** OS of GLASS patients (recurrent tumors) stratified by the OSM-activation signature. **(m)** OS of *Spitzer* patients (primary tumors) stratified by the OSM-activation signature. **(n)** OS of *Spitzer* patients (recurrent tumors) stratified by the OSM-activation signature.

The Edge-unique signature had been derived from cells maintained in culture for a few passages. To produce a signature that tentatively would mimick Edge cells in a peritumoral microenvironment (immune cell-activated) a DGE analysis of OSM-stimulated versus non-stimulated Edge cell transcriptome data was performed (Table S12). This produced the OSM-activation signature of 22 genes (its small size supporting previous notions of minor transcriptomic changes in OSM-stimulated Edge cells). Applying this signature (Table S13) on the TCGA data showed a similar result as with the Edge-unique signature, significant for PFI but not for OS (Fig. 6i-j). In the GLASS cohort the signature had no prediction value in either primary nor recurrent data (Fig. 6k-l), while it significantly separated patients for both primary and recurrent data in the *Spitzer* cohort (Fig. 6m-n). Overall, both Edge-related signatures showed prediction value in two of three GB cohorts supporting a central role of peritumoral GB cells for therapy response and tumor relapse.

## DISCUSSION

The therapeutic progress for GB has been sparse despite decades of immense scientific advances. One critical and understudied area is that of local peritumoral GB cells, particularly in direct comparison with their matched bulk tumor. These cells reside in regions that are generally unavailable to surgery and where most relapses occur^7–9,55^, making them essential targets for new therapies. The scarcity of peritumoral samples has also led to a lack of functional models representing these regions. Here we aimed to collect, culture and analyze matched patient samples from the bulk tumor and a local peritumoral site. The tumor border delineating the bulk tumor was determined by 5-ALA fluorescence which extends beyond the MRI margin^5^, and all Edge samples were collected at least 10 mm beyond the fluorescent border. We took careful measures to reduce the risk of contaminating Edge samples with bulk tumor cells, to maintain tissue viability during transport and to process each sample immediately after resection. These actions likely contributed to the high (50%) success rate of Edge cell cultures despite their often minute sample sizes and low tumor-cell content. Functional analyses of patient-matched bulk and Edge cell cultures revealed distinct and similar differences for central cancer cell characteristics across patients highlighting preservation of the Edge-specific phenotype under culture conditions.

Genomic analyses failed to elucidate Edge-specific aberrations. Phylogeny proposed that Edge cells had populated their niche early with limited subsequent genetic diversification. Only one patient displayed an Edge-unique subclone relative to the matched bulk tumor, and the corresponding recurrent tumor contained the same set of subclones as the primary Edge sample. We concluded that Edge cell tumor progression was not primarily driven by genomic aberrations but likely shaped by epigenetic alterations induced by the microenvironment. Single cell multiome profiling supported this notion, uncovering distinct transcriptomic and epigenomic landscapes of matched bulk and Edge GB cells. The molecular profiles of Edge cells exhibited greater heterogeneity and occurred in a continuum of bulk-like to Edge-like phenotypes. Across patients, Edge cells consistently displayed a shift toward more differentiated cell states, accompanied by distinct chromatin accessibility signatures and gene regulatory circuits. Chromatin-accessibility-based subcluster analysis identified two subgroups of Edge cells that were also recognizable by their high expression of the Edge-unique signature. Moreover, both subgroups displayed significantly elevated expression of a range of mesenchymal and immune cell state signatures relative to bulk GB cells. Integrated computational, phenotypic and functional analyses suggested microglia-derived OSM as a driver of the Edge-unique phenotype. The fact that Edge cell cultures remained primed to functionally respond to OSM while cell state and Edge-related molecular changes were modest provided additional support for their phenotypic stability and relevance as local peritumoral GB cell models.

Several longitudinal studies of matched primary and recurrent GB tumors have described an increased proportion of mesenchymal-like cells in relapse samples^16,25,56,57^. This evolutionary trajectory has been suggested to be treatment-induced, while our data propose an alternative explanation. We show that our treatment-naïve GB Edge cells isolated from relapse-prone peritumoral regions are more mesenchymal-like than their matched bulk tumor cells across patients. We also show, for one patient, that Edge cells are central in the recurrence. Thus, the commonly observed MES trajectory in recurrent tumors could have been produced by comparing the relapse sample with its incorrect source, i.e. the primary bulk tumor rather than the peritumoral compartment. A recent study describing more diverse and unpredictable trajectories^54^ aligns with this notion and with the broader challenge of identifying generalizable clinically relevant knowledge across different GB patient cohorts. This was further emphasized in our study where the Edge-unique and OSM-activation signatures could predict patient outcome in two out of three cohorts. The fact that the Edge-related signatures were produced from a small number of GB patients and most likely had not captured the full complexity and heterogeneity of the peritumoral GB cell landscape would also have had an impact on the survival analyses.

Our investigation describes novel insights into the biology, phenotype and regulation of peritumoral GB cells and presents a unique resource of functional models. Given the inter-patient and intra-tumor heterogeneity of GB, and the complexity of its microenvironment some caveats are the limited number of patients and that the single cell multiomic profiling was performed cell cultures and not tissue samples. To compensate for these limitations we validated our results in publicly available independent GB cohorts^17–20,54,58^. Yet, our work underscores the need for placing peritumoral GB cells at the center of GB research with continued investigations of matched primary Edge and recurrent tumor samples to delineate progression mechanisms and identify therapeutic vulnerabilities. Sampling efforts should also be broadened to include multiple peritumoral and relapse samples, and more patient-derived models are needed that reflect these spatial and temporal GB cells.

## METHODS

### GB patient cohort and sample collection

The collection and use of human samples was carried out at Uppsala University Hospital according to the ethical permit 2007/353 and approved addendums 2013, 2016, 2019, 2020, 2021, 2022, 2024, and following informed written consent. GB tissue and peripheral blood samples were collected from 11 patients with radiologically suspected GB that were eligible for total or supratotal 5-ALA (Gliolan^®^) fluorescence-guided resection. Patient samples were deidentified before processing. All patients were diagnosed with GB IDH-wildtype according to the 2021 WHO classification^1^. Table S1 provides a summary of the patient cohort and clinical data.

Gliolan was administered 4-6 hours before surgery (20 mg/kg). The neuronavigation system (Brainlab), ultrasound device (BK medical), and fluorescent filter operating microscope (Leica) were used. After craniotomy and exposure of the brain cortex, the tumor was localized and confirmed with the neuronavigation system. Cortical-subcortical resection was achieved with a cavitron ultrasonic surgical aspirator (CUSA). When the central part of the tumor was reached, according to anatomical and intraoperative findings, the fluorescent filter was activated, revealing the hyperintense pink color distinctive for high-grade gliomas. The highest fluorescent tumor area was located and marked on MRI sequences with neuronavigation probes and resected *en bloc*. This sample was called Core. Tumour resection continued using the CUSA and was guided by the 5-ALA fluorescence. All tissue resected with CUSA was collected and labeled Aspirate. When no fluorescence could be detected, the resection cavity was filled with a physiological solution, and navigated ultrasound images were used to confirm total resection compared to preoperative images. The cavity was generously irrigated with water to remove blood clots. To avoid cell contamination between the bulk (Aspirate and Core) and Edge samples, a new set of instruments was used to collect the Edge sample from a peritumoral non-fluorescent and non-eloquent region of the surgical cavity to minimize the risk of postoperative neurological sequelae. The sampled area was recorded on the preoperative MRI images with the neuronavigation probe. The surgical cavity was then irrigated one more time, and hemostasis was performed with hemostatic material (Surgicel) and fibrin glue (Tisseel). Images of patient tumors with annotated samples were created using the Brainlab Element software (Brainlab AG).

Samples were placed in Hibernate-A medium (Gibco, A1247501) and immediately transported to the lab on ice where it was divided and processed. Some was frozen in RNAlater Stabilization Solution (Invitrogen, AM7020), some formalin-fixed and paraffin-embedded (FFPE), and some was processed into a single-cell suspension using Mitenyi’s Biotech Brain Tumor Dissociation Kit (130-107-677) following the manufacturer’s instructions.

### Establishment of patient-derived GB cell cultures

Single-cell suspensions were explanted and maintained in on DMEM/F12 Glutamax (Gibco, 11320033) and Neurobasal medium (Gibco, 21103049) 1:1 supplemented with 1% B27 (Gibco, 12587010), 0.5% N2 (Gibco, 17502001), 1% penicillin/streptomycin (Sigma, P4333), 10ng/mL EGF (Peprotech, AF-100-15-1MG) and FGF (Peprotech, 100-18B-1MG) and defined as NSC medium, as described previously^59^. Briefly, single-cell suspensions derived from patient GB tumors were seeded in NSC media and left undisturbed in the incubator. After 3 days, fresh media was added and 7-10 days after seeding primary spheres were collected, gently resuspended, and seeded adherently in laminin-coated flasks.

### Authentication and mycoplasma testing of patient-derived cell cultures

All established cell cultures (that reached passage 7 and beyond) were authenticated through STR profiling (Table S2). DNA from patient tissue and corresponding explanted cell culture was extracted using Qiagen DNeasy® Blood & Tissue Kit (Qiagen) following manufacturer’s instructions. DNA was analyzed by Eurofins Genomics (Germany) using the Cell Line Authenticity Basic Service testing 16 DNA markers.

Cultures were also regularly tested for mycoplasma, and all cultures used in this study were negative. Conditioned culture medium was collected, centrifuged at 250g to remove debris, and pelleted by centrifugation at 13,000rpm for 10 min. Pellets, when present, were resuspended in PCR-grade water and incubated at 95°C for 3 min. PCR was performed using primers Mico1 (5′-GGCGAATGGGTGAGTAACACG-3′) and Mico2 (5′-CGGATAACGCTTGCGACTATG-3′) at 0.4 μM each in reactions containing PCR buffer, 0.6 mM dNTPs, 3 mM MgCl₂, and 0.03 U/μL GoTaq polymerase (Promega). Cycling conditions were 94°C for 4 min; 35 cycles of 94°C for 30s, 60°C for 45s, and 72°C for 30s; and 72 °C for 7 min. PCR products were analyzed on 2% agarose gels, with the presence of a ∼0.5 kb amplicon indicating mycoplasma contamination.

### Sphere formation assay of acutely dissociated cell suspensions

Primary tumor spheres were established by seeding dissociated cells on laminin-coated plates and maintained in NSC medium with media replacement every 3 days. The duration of primary culture varied across samples (1–4 weeks), depending on culture viability and growth, Adherent spheres were detached using TrypLE (Invitrogen, 12563011), dissociated and seeded (1000 cells/well) in a 24-well low attachment plate (Corning) in 8 technical replicates for secondary sphere assays. After 7 days, the number of secondary spheres was counted, collected, dissociated into single-cell suspensions (TrypLE), and seeded (1000 cells/well) for tertiary sphere assays. Spheres were counted 7 days later.

### Extreme Limiting Dilution Assay (ELDA) of established cultures

Patient-derived cell cultures grown adherently were seeded as serial dilutions in ultralow attachment 96-well plates (Corning, #3474) with the highest density at 200 cells per well and the lowest at 1 cell per well. Each cell dose was seeded in 10 technical replicates. Each ELDA plate was counted as a technical replicate. After plating, the ELDA plates were incubated at 37°C and 5% CO2 for 7 or 10 days when all wells were scored for the presence or absence of spheres. The minimal number of cells needed to form a sphere was calculated using the extreme limit dilution analysis software^60^ in the statmod package for the R programming environment.

### Collagen invasion assay

1000 cells at passages 4-7 were seeded in a 96-well Nunclon Sphera U-Shaped-Bottom Microplate (Thermo Scientific, 174925) at 10-16 technical replicates and centrifuged at 130g for 10 minutes. The plate was incubated at 37°C, 5% CO_2_ for 3 days. A collagen gel mixture composed of 5% 0.1M NaOH, 2% 1M HEPES (Invitrogen, 15630056), 3% 7.5% NaHCO3 and 50% bovine collagen type I (Cell systems, 5005-100ML) in 40% NSC media was added to the top of the spheroid. The photos were taken at 0 hours, 24 hours, and 48 hours after collagen addition, and Fiji 1.53f was used to measure the invasion area of each sphere.

### Proliferation analysis through BrdU incorporation

Patient-derived cell cultures grown as adherent were dissociated using TrypLE and seeded on laminin-coated coverslips (VWR, 631-1577P). 1μg/μl BrdU (Sigma, B5002) was added at 70-80% cell confluency for a pulse of 16 hours. Cells were permeabilized with 2M HCl for one hour at room temperature. After washing with PBS, cells were permeabilized using a solution containing 0.2% TritonX-100 and 3% bovine serum albumin (Sigma, A7906) in PBS. Nonspecific binding was blocked using a solution containing 0.2% TritonX-100 and 5% bovine serum albumin (Sigma, A7906) in PBS containing 0.2% TritonX-100 for 1 hour at room temperature. The anti-BrdU primary antibody (Abcam, ab6326) was diluted in a blocking solution and incubated overnight at 4°C. After washing in PBS-T, cells were incubated with secondary antibody dilution – Alexa Fluor™ Plus 488 anti-Rat (Invitrogen, A48269) - dilution in a blocking buffer for 1 hour at room temperature. After washing with PBS-T, cells were mounted on a microscope slide on a microscope slide with Fluoromount-G mounting medium, with DAPI (Invitrogen, 00-4959-52). Following staining, up to 10 pictures with at least 20 cells were taken using a Leica DMi8 microscope. Images were analyzed using Fiji 1.53f.

### Immunohistochemical (IHC) analyses of tissue sections

FFPE blocks were sectioned (6**μ**m) using a microtome (HM325 rotary microtome, Thermo Fisher Scientific). Slides were dried at 37°C overnight and stored at 4°C until use. Tissue sections were deparaffinized in xylene and hydrated in decreasing concentrations of EtOH. Sections were heated in citric acid buffer (Vector laboratories, H-3300-250) supplemented with 0.05% Tween-20 (v/v) and allowed to cool for 2 hours. The tissue sections were outlined with an ImmEdge® Hydrophobic Barrier PAP Pen (Vector Laboratories, H-4000) and washed with TBS containing 0.05% Tween-20 (v/v) (TBS-T). Nonspecific binding was blocked using the BrightVision 2-step detection system, Step 1 (Immunologic, DPVB55HRP) for 1 hour at room temperature. The tissues were incubated in primary antibodies – Stem121 (Takara Bio, Y40410), Sox2 (Millipore, AB5603), MPO (Proteintech, 22225-1-AP), CD14 (CST, #75181), IBA1 (Wako, 0119-19741), and P2RY12 (Abcam, ab254347) - diluted in normal antibody diluent (Immunologic, BD09-125) overnight at 4°C. After washing in TBS-T, the sections were incubated for 10 minutes in 3% H_2_0_2_ (Sigma, H1009) diluted in TBS-T. The tissue sections were washed in TBS-T and incubated with secondary antibody solution BrightVision 2-step detection system, step 2 (Immunologic, DPVB55HRP) for at least 30 minutes. After washing with TBS-T, DAB substrate (Vector Laboratories, SK-4100) was added. The tissue sections were then counterstained with hematoxylin, dehydrated at increasing concentrations of EtOH, and mounted in Pertex. For histology analysis, tissue sections were deparaffinized in Xylene and hydrated at increasing concentrations of EtOH. The tissue sections were counterstained with hematoxylin, followed by dehydration at increasing concentrations of EtOH and Pertex mount.

### Immunofluorescence analyses of tissue sections

FFPE blocks were sectioned, dried, and stored as outlined for IHC analyses. Slides were incubated at 60°C for 1 hour prior to deparaffinization, rehydration, antigen retrieval, hydrophobic barrier generation and washing as outlined for IHC analyses. Sections were permeabilized with solution containing 0.2% TritonX-100 and 3% bovine serum albumin (Sigma, A7906) for 5 minutes at room temperature. Nonspecific binding was blocked with blocking buffer prepared in PBS, containing 0.2% TritonX-100 and 5% bovine serum albumin for 1 hour at room temperature. Primary antibody against Sox2 (R&D, AF2019) was diluted in blocking buffer and incubated overnight at 4°C. Slides were washed with 0.2% TritonX-100 in PBS (PBS-T) and incubated with secondary antibody (Alexa Fluor™ Plus 555 anti-Goat (Invitrogen, A32816) diluted in blocking buffer for 1 hour at room temperature. After washing with PBS-T, slides were mounted with a glass coverslip and Fluoromount-G mounting medium with DAPI (Invitrogen, 00-4959-52). A minimum of 10 randomly selected regions were imaged the following day with LSM700 confocal microscope at 40X magnification. Representative images were acquired at 60X magnification. Fiji 1.53f Point Tool was used for subpopulation analysis with manual cell count.

### Immunofluorescence analyses of fixed cells

Cells grown as adherent cultures were dissociated with TrypLE, seeded on laminin-coated coverslips (VWR, 631-1577P), and fixed after 24 hours with 4% formaldehyde (Histolab, 02176). The cells were washed with PBS. Cells were permeabilized using a solution containing 0.2% TritonX-100 and 3% bovine serum albumin (Sigma, A7906) in PBS. Nonspecific binding was blocked using a solution containing 0.2% TritonX-100 and 5% bovine serum albumin (Sigma, A7906) in PBS containing 0.2% TritonX-100 for 1 hour at room temperature. Primary antibodies Sox2 (Millipore, AB5603) and were diluted in a blocking solution and incubated overnight at 4°C. After washing in PBS-T, cells were incubated with secondary antibodies (Alexa Fluor™ Plus 555 anti-Rabbit (Invitrogen, A32732)) diluted in blocking buffer for 1 hour at room temperature. After washing with PBS-T, cells were mounted on a microscope slide on a microscope slide with Fluoromount-G mounting medium with DAPI (Invitrogen, 00-4959-52).

### Patient-derived orthotopic xenograft models

All animal experiments were conducted following the guidelines and regulations of Uppsala University and were approved by the local animal ethics committee (ethical permits 08446-2019 and 5.8.18-02391/2024). Six to eight weeks old NOD.CB17-Prkdcscid/NCrHsd mice (Envigo) of both sexes were orthotopically transplanted with patient-derived cell cultures (P<10) using a stereotaxic injector (Stoelting CO). A total of 200,000 cells were injected in the subventricular zone using the coordinates: 0,5 mm anterior of bregma, 1,5 mm right of the midline, and 2 mm deep. Mice were housed in groups of up to 5 animals and maintained on a 12-hour light-dark schedule with gradual dawn and rise for 30 minutes with a temperature of 22 ± 2°C and relative humidity of 55 ± 10%. Animals were monitored several times a week for disease-symptoms and weight-loss and were sacrificed when they reached the humane endpoint or at the experimental endpoint of 1 year. The brains were collected, formalin-fixed, and paraffin-embedded.

### Microscopy and image processing

The sphere formation, ELDA, and invasion assays were imaged using an Eclipse TS 100 Nikon microscope using a 10X or 20X objective with the software. Images were analyzed using Fiji 1.53f. Tumor tissue and xenografted mouse brains were imaged using a ZEISS AxioScan for Brightfield and ZEISS LSM700 confocal microscope for immunofluorescence using a 10X, 20X, 40X or 60X objective with the Zen software (Zeiss). Images were cropped and edited using Adobe Photoshop (Adobe) and Adobe Illustrator (Adobe). The BrdU pulse and immunofluorescent staining of fixed cells on coverslips were imaged using a Leica DMi8 microscope. Images were analyzed using Fiji 1.53f or QuPath^61^. For immunohistochemistry analysis using QuPath, positive cells were identified with DAB mean threshold range of 0.15-0.35.

### Whole exome sequencing and analysis

DNA was extracted from frozen tumor tissue using Qiagen DNeasy® Blood & Tissue Kit (Qiagen) following the manufacturers’ instructions with the modification of incubating in ATL buffer supplemented with 10% proteinase K at 56°C overnight until completely lysed. DNA was extracted from patient-derived cell cultures (passages 4-6) grown adherently using Qiagen DNeasy® Blood & Tissue Kit and following manufacturers’ instructions. Quality control of the extracted DNA was performed using TapeStation (Agilent), and the concentration was determined using Qubit (Invitrogen). Library preparation was performed using the Twist Comprehensive Exome Kit (Twist Biosciences) following the manufacturer’s instructions. Libraries were sequenced (SNP&SEQ Technology Platform, Uppsala University) using cluster generation and 100 cycles of paired-end sequencing in NovaSeq SP flowcell using the NovaSeq 6000 system and v1.5 sequencing chemistry (Illumina Inc.) aiming at a minimum of 650 million read pairs per flow cell, with at least 80% of the bases having had a base quality score of 30 or higher. The raw reads were aligned to the human genome version GRCh38.13 using the Burrows-Wheeler Aligner BWA-MEM^62^ (v.0.7.17). Duplicate reads were then marked using Samtools^63^ (v.1.14). WES data is deposited at EGA with ID: EGAD50000002097.

#### Variant calling

Somatic variants were called using Mutect2 from the GATK suite^64^ (v.4.2.0.0). To assess technical artifacts, a panel of normals was provided from the 1000 genomes project^65^. According to GATK best practices, GnomAD was used as a germline resource during the calling process^66^. Both the normal panel and the somatic resource were downloaded from https://console.cloud.google.com/storage/browser/gatk-best-practices/somatic-hg38. The variants that passed post-processing by GATK FilterMutectCalls were kept for downstream analyses. The remaining variants were annotated using snpEff and depth filtered using snpSift^67^. The resulting vcf files were then translated to maf format and oncoprints were generated using Maftools^68^. Variant calling data is deposited at EGA with ID: EGAD50000002096.

#### CN calling

Copy number profiles for matched tumor-normal samples were obtained using the Python package CNVkit batch pipeline^69^. Since we lacked normal samples for patient 1 and edge tissue samples are mainly composed of healthy cells, they were used as a paired normal reference in the analysis of aspirate and core tissue samples for that patient. The resulting copy number estimates were then visualized in a heatmap using the R library KaryoplotR^70^.

#### Phylogenetic tree inference and clonal hierarchies

Clonal hierarchies were investigated in a two-step process, where first PyClone^71^ was used to generate gene clusters and calculate the cellular prevalence of these clusters based on variant allele frequencies and copy number status from all samples within the same patient. Secondly, to model the clonal dynamics of the gene clusters and infer a phylogenetic relationship, we used CITUP with standard parameters^72^.

### 10X multiome single-nucleus (sn) sequencing

Fresh single-cell suspensions were generated from patient-derived matched Core and Edge cell cultures of patients 13, 14 and 17 grown adherently at passages 4-5. Cells were resuspended in PBS with 0.04% BSA and lysed in lysis buffer (10mM Tris-Hcl pH 7.4, 10mM NaCl, 3mM MgCl2, 0.1% Tween-20, 0.1% Nonidet P40 Substitute, 0.01% Digitonin, 1% BSA, 1mM DTT, 1U/uL RNase inhibitor 40 U/uL). Cells were filtered through a 40**μ**m cell strainer before their integrity and quantity were evaluated with SYBR Green stain. Cells were resuspended according to 10X Genomics concentration guidelines to obtain a target of 6,000 to 10,000 nuclei per sample. Construction of the sn Assay for Transposase-Accessible Chromatin using sequencing (ATAC) and RNA libraries was performed using the Chromium Next GEM Single Cell Multiome ATAC + Gene Expression kit, according to the manufacturer’s instructions. Libraries were quality controlled using Agilent Technologies 2100 bioanalyzer using a DNA 1000 chip, whereafter they were sent to Macrogen Europe (Amsterdam) for sequencing using 150bp paired-end cycles (Illumina), with the aim of a minimum sequencing depth of 20k reads/nucleus.

The snRNA-seq and snATAC-seq data were processed simultaneously using 10X Genomics Cell Ranger ARC software (v1.0.1). Reads were demultiplexed and aligned to the human reference transcriptome (GRCh38) using the STAR aligner. Raw data is deposited at EGA with ID: EGAD50000002098.

### snRNA-seq analyses

#### Quality control

Cells were identified based on unique cell barcodes within the library. Cells containing the number of counts < 1000 and the number of detected feature genes < 1000 and mitochondrial percentage > 0.20 were filtered. R package DoubletFinder^73^ was used to distinguish between empty droplets and droplets containing multiple cells. The raw matrix of gene counts versus cells generated by Cell Ranger was subsequently filtered to exclude unqualified cells. After preprocessing and quality control we obtained 11985 snRNA-seq profiles from core cells and 15685 profiles from edge cells.

#### Pre-processing

Seurat package (v.4.3.0)^74^ was used to perform the downstream analysis. Expression normalization was performed using the LogNormalize function. To account for inter-patient variability, datasets from three patients were integrated using canonical correlation analysis (CCA)^75^. Integration anchors were identified using FindIntegrationAnchors with the first 30 canonical correlation vectors. The integrated assay was generated using IntegratedData. To correct for potential effects in library size, the number of UMIs, mitochondrial content, and cell-cycle difference, a linear model was applied during gene scaling and centering. Unwanted variations associated with cell cycle phases were removed using the “Alternate Workflow” outlined in the Seurat vignette (https://satijalab.org/seurat/v2.4/cell_cycle_vignette.html) based on the expression level of previously published G2/M and S-phase gene signatures. Expression values were scaled across all the cells within the dataset. Principle components analysis (PCA) was performed on the top 2000 most variable genes. Significant principal components were used as inputs for nonlinear dimensionality reduction techniques via Uniform Manifold Approximation and Project (UMAP) as well as cell clustering.

#### Cell cycle analysis

Cell-cycle activity was quantified in the scRNA-seq dataset using a curated panel of canonical S-phase and G2/M-phase gene signatures^76^. Normalized expression matrices were scored at the single-cell level using the AUCell^77^ package, applying a combined S-phase and G2/M-phase gene signatures to generate a cell-cycle activity score for each cell. The resulting cell-cycle activity pattern was visualized using UMAP embedding, and cell-cycle score distributions were summarized across snRNA-defined clusters using barplots.

#### Clustering analysis

To identify clusters of transcriptional similar cells, a shared nearest neighbor (SNN) graph was constructed using the FindNeighbors function with the first 30 canonical correlation vectors. Graph-based clustering was performed using the Louvain algorithm implemented in the FindClusters with clustering resolutions of 0.1 to define distinct clusters.

#### Differential gene test

Differentially expressed genes (DEGs) between clusters were calculated using the wilcoxauc function from the presto package (https://github.com/immunogenomics/presto). Genes were considered differentially expressed with the following criteria: p-value < 0.01, log2FoldChange >0.5, AUC > 0.6.

#### Gene Set Enrichment Analysis

Differentially expressed genes ranked by log2 (fold change) and FDR were analyzed using GSEA^78^. Pathways were selected from hallmark gene sets and filtered for a minimum size of 15 and a maximum size of 200 genes. Enriched pathways with FDR < 0.05 were considered significantly enriched.

#### Gene Signature Analysis across public datasets

Differentially expressed genes from core-like and edge-like clusters were used as gene signatures. Gene activity in single cells was quantified using the AUCell^77^ package, with scores normalized to the [0,1] range. Processed data matrices from published studies (D*armanis et al., 2017, Yu et al., 2020, Schmassmann et al., 2023*)^17,20,21^ were downloaded, and core-like and edge-like gene activity was accessed using AUCell and displayed with z-score. Gene signatures, including MES_CORE_GENES, interferon, MHCI, and MHCII modules (*Chanoch-Myers et al. 2022*)^26^, were assessed in both core-like and edge-like datasets. For bulk RNA-seq datasets (*Andrieux et al., 2023, Bastola et al., 2020*)^18,19^, matrix counts were normalized using CPM (count-per-million) and log2 transformation. Gene signatures from core-like and edge-like clusters were integrated using a non-negative matrix factorization (NMF) algorithm to assess clustering consistency across these datasets.

#### Analyses of *Neftel* and *Richards* cell states

Cell states were mapped following *Neftel et al. 2019*^13^. Cells were scored for NPC1/2 and MES1/2 gene signatures using the AUCell package, and scores were averaged to represent NPC and mesenchymal states. Cells were divided into OPC/NPC and astrocyte/mesenchymal lineages using a transcriptional program score, D = max(SC_OPC_, SC_NPC_) - max(SC_AC_, SC_MES_). Dimensional coordinates for OPC/NPC and astrocyte/mesenchymal states were computed based on log-transformed differences in signature scores. Gene signatures associated with developmental and injury-related cellular status were analyzed in cells from our datasets using a similar scoring method. Barplots were generated to visualize the distribution of these signatures across cell states.

### snATAC-seq analyses

#### Quality control

For each patient-derived culture sample, a list of unique ATAC-seq fragments with associated barcodes was generated using Cell Ranger. The list of unique fragments per barcode for each patient-derived culture sample was read into the Signac^79^ package to perform quality control and doublet removal for each patient dataset individually. To enrich cellular barcodes, the bimodal distribution of log10(TSS enrichment+1) and log10(number of unique fragments) were used to distinguish cellular from non-cellular barcodes. Barcode cutoff thresholds were set as 2 for log10(TSS enrichment+1) and 3 for log10(number of unique fragments). Only barcodes exceeding these thresholds in both metrics were retained as cellular barcodes for downstream analysis. The peak-barcode matrices generated after this filtering step were used for further analysis. After preprocessing and quality control, we obtained 9824 profiles from core cells and 12917 profiles from edge cells for snATAC-seq.

#### Cell cycle analysis

Cell-cycle activity in the scATAC-seq dataset was quantified by scoring chromatin accessibility at regulatory regions with S-phase and G2/M-phase gene programs^76^. Following TF-IDF normalization and LSI-based dimensionality reduction, peaks around promoter-proximal peaks (-2000 to +2000 bp around transcription start site) were annotated to their corresponding cell-cycle genes. For each cell, phase-specific chromatin accessibility scores were computed by aggregating normalized accessibility values across promoter peaks belonging to each signature. The resulting cell-cycle activity patterns were visualized using UMAP embedding, and their distribution across snATAC-defined clusters was summarized using barplots.

#### Clustering analysis

Chromatin accessibility was analyzed in all nuclei for which there were dual qualified profiles, i.e. both ATAC and RNA. Normalization was performed using Term Frequency-Inverse Document Frequency (TF-IDF) transforms using the function RunTFIDF, which adjusts peak counts based on their frequency across cells to emphasize informative features. Top accessible peaks were identified using FindTopFeatures with a cutoff of 20 peaks per cell (min.cutoff =20). Dimensionality reduction was performed using Singular Value Decomposition (SVD) via RunSVD(), generating Latin Semantic Indexing (LSI) components. UMAP embedding was computed based on LSI reduction with the top 50 components. To correct for patient-specific batch effects, a harmony algorithm was used based on the LSI-reduced dimensions with the function RunHarmony. The harmony integration generated a batch-corrected low-dimensional embedding, which was used for downstream clustering and visualization. A shared nearest neighbor (SNN) graph was constructed using FindNeighbors. The clustering of cells was performed using the Louvain algorithm via FindClusters.Differential peak accessibility analysis was performed using the wilcoxauc function from the presto package. This procedure identified differential accessibility peaks (DEPs) between two groups of cells with the threshold: Benjamini-Hochberg corrected p-value < 0.01, log2FoldChange >= 0.1, and AUC > 0.6. Marker peaks were annotated to the nearby gene using clusterProfiler, with a q-value cutoff of 0.05 after multiple comparison adjustments with the Benjamini-Hochberg procedure.

#### TF motif enrichment analysis

Transcription factor (TF) motif enrichment analysis was performed using chromVar^80^ with the JASPAR2020 motif database^81^. Significant TFs enriched in each cluster were identified to represent unique features. The chromatin accessibility matrix was used as input to calculate TF accessibility deviation scores for each cell across the entire sample set. TF motifs that were positively correlated with a specific group were selected to represent that cluster. Finally, deviation z-scores were visualized as a heatmap for marker TFs.

### Integration analyses of sn multiome data with public datasets

#### Integration of snRNA-seq profiles with TCGA subtype-defined bulk RNA-seq data

From each dataset (*Lu, Maturi, et al., 2020*^27^ and present study), genes exhibiting high variability across cells were selected to represent diverse biological features. These high-variability genes were independently identified and ranked in each dataset, and the shared top 2000 genes chosen from both datasets were used for further analysis. The FindVariableFeatures function in the Seurat package was used to analyze these selected genes. Subsequently, anchors were sought using FindIntegrationAnchors. Anchors were identified as pairs of cells (one from each dataset) that exhibit a common biological state. To facilitate this process, Canonical Correlation Analysis^82^ was initially conducted for dimensionality reduction. Following dimensionality reduction, the K-nearest neighbors for each cell within its respective dataset were computed. Finally, the mutual nearest neighbors were identified.

#### Integration of snATAC-seq profiles with ATAC subcluster-defined bulk ATAC-seq data

Peaks with more than 20 counts were selected from each dataset to capture diverse biological features. The FindTopFeatures function in Signac was used to identify these peaks. Integration was performed using FindIntegrationAnchors, which identifies anchor pairs as cells from different datasets sharing a common biological state. Dimensionality reduction was performed using Harmony to address batch effects. Following dimensionality reduction, K-nearest neighbors were computed for each cell within its dataset, and mutual nearest neighbors were identified to facilitate integration.

#### Integration of snRNA-seq profiles with *Schmassman* scRNA-seq data of GB samples

The processed *Schmassmann et al.,* 2023^21^ single-cell data was downloaded from GEO accession GSE197543^21^. Samples included in this analysis represented IDH wt samples with ID BTB_604, BTB_616, and BTB_617. To integrate our in-house dataset with publicly available single-cell RNA-seq data, the default assay for both datasets was set to use their normalized RNA assay. Features were identified for integration using SelectIntegrationFeatures with default parameters. After that, integration anchors were identified using FindIntegrationAnchors with parameters: dims = 1:30, k.anchor=50 and k.score=50. Integration was performed using the IntegrateData function, with LogNormalize as the normalization method and k.weight=50 to control the contribution of anchors. The integrated dataset was scaled and centered using ScaleData and then subjected to dimensionality reduction using PCA. UMAP was applied to the PCA-reduced space using the first 30 principal components. To analyze cell-cell communication, we used the CellChat^40^ package with the integrated Seurat object. The CellChatDB.human was loaded and secreted signalling interaction was selected using subsetDB. The Seurat object was then converted to a CellChat object using the createCellChat function. The recommended workflow was followed to infer the cell state-specific communications (using identifyOverExpressedGenes, and identifyOverExpressedInteraction with the default parameters). Finally, cell-cell communication networks were generated across ligand-receptor pairs to generate a summary of overall communication strength with population size scaling enabled (using computeCommunProb with population.size = TRUE). Cell-type proportions were calculated as the fraction of M1/M2-polarized macrophages or activated/surveilling microglia relative to the total number of cells within the Center or Periphery region of the IDH wt *Schmassmann* samples.

### Analyses of OSM-stimulated Edge cell cultures

#### OSM stimulation

GCCs derived from edge samples were seeded at a density of 5 × 10⁵ cells per well in 6-cm culture dishes pre-coated with laminin. After a 24-hour incubation period, half of the wells were treated with 50 ng/mL (diluted in PBS) of human Oncostatin M (Peprotech, #300-10-50UG) and control cells were given PBS only (vehicle control (VC)). New OSM was added every 48 hours, at day 1, 3, 5, and 7, and cells were used at day 8 for RNA and protein extraction, Extreme Limiting Dilution Assay (ELDA), invasion assays, and cell viability assessment.

#### Viability assay

Edge cells were seeded at a density of 5 x 10^4^ cells per well in clear-bottom 96-well plates (Corning, #3610) with 10 technical replicates for VC and OSM-stimulation respecitvely. Cells were treated as described above and cell viability was assessed on day 8 using CellTiter-Glo® 2.0 Reagent (Promega, G9248) according to the manufacturer’s instructions. Briefly, CellTiter-Glo® 2.0 was equilibrated to room temperature and added to the wells in at a 1:1 ratio with the existing media. Plates were incubated in dark at 37°C for 10 minutes and luminescence was recorded using a CLARIOstar Plus microplate reader (BMG Labtech). Luminescence values from OSM-treated wells were normalized to the matched VC wells and expressed as relative viability (OSM+/VC) with mean ± SEM. The assay was repeated three times.

#### Extreme Limiting Dilution Assay

OSM-treated and control (VC) Edge cells were harvested at the end of the treatment period after dissociation with TrypLE (Gibco™, 12563011). Matched treated and VC cells were seeded, incubated, scored and the estimated number of cells needed to form a sphere calculated as previously detailed for the ELDA assay. For each independent patient-derived culture, ELDA estimates were averaged across biological replicates. ELDA-derived stem cell frequencies were calculated as the inverse of the estimated number of cells required for sphere formation and normalized to matched VC. Data are expressed as relative stem cell frequency (OSM+/VC) with mean ± SEM.

#### Collagen invasion assay

OSM-treated and control (VC) Edge cells were harvested at the end of the treatment period after dissociation with TrypLE (Gibco™, 12563011). Matched treated and VC cells were seeded at a density of 500 cells per well in a 96-well Nunclon Sphera U-Shaped-Bottom Microplate (Thermo Scientific, 174925) in 8 technical replicates and centrifuged at 1000 rpm for 4 minutes. The plate was incubated and spheroids embedded with a collagen mixture as detailed previously. Images were taken at 0 hours, 24 hours and 48 hours using the Eclipse TS 100 Nikon microscope with a 10X objective, with one pixel corresponding to 0.49 mm. Image analysis was conducted in Fiji 1.53f, where pixel-to-length ratio was calibrated globally. For each technical replicate, the 0-hour spheroid borders were overlayed to define the centre of the matched spheroid at 24 hours. Distance to the farthest invaded cell in collagen was measured from the centre. For each independent patient-derived culture, maximum invasion distances were summarized using the mean. Mean distances from OSM-treated spheroids were normalized to matched VC and visualized with mean ± SEM per patient.

#### Western blot analysis

Edge cells from three independent OSM-stimulations have been analyzed with similar results. Cells were lysed using RIPA buffer (Thermo Scientific, 89900) supplemented with protease (Roche, CO-RO) and phosphatase inhibitors (Roche, PHOSS-RO). Protein concentrations were measured with the Pierce™ BCA Protein Assay Kit (Thermo Scientific, 23225). Equal amounts of protein samples were resolved on 4–12% Bis-Tris polyacrylamide gradient gels (NuPAGE, Thermo Scientific) and transferred onto nitrocellulose membranes (Novex, Thermo Scientific). Membranes were blocked with EveryBlot Blocking Buffer (BioRad, 12010020) and incubated overnight at 4°C with the following primary antibodies: rabbit anti-pSTAT3 (#9145, CST), mouse anti-STAT3 (#9139, CST), rabbit anti-pERK1/2 (#9101, CST), rabbit anti-ERK1/2 (#9102, CST), rabbit anti-pAKT(#4060, CST), rabbit anti-AKT(#4691, CST) and rabbit anti-GAPDH (#5174, CST). Following primary antibody incubation, membranes were incubated with HRP-conjugated secondary antibodies (anti-mouse, GE Healthcare, #NXA931; anti-rabbit, GE Healthcare, #NA9340) and developed using SuperSignal™ West Femto Maximum Sensitivity Substrate (Thermo Scientific, 34095). Signals were visualized with the Amersham™ Imager 680 system (GE Healthcare). For membrane reprobing, Restore™ Western Blot Stripping Buffer (Thermo Scientific, 21059) was used according to the manufacturer’s protocol. Quantitative densitometry was performed using ImageJ2 software (Version 2.14.0).

### RNA sequencing

RNA was extracted from Edge cell cultures using the DNeasy Blood and Tissue Kit (QIAGEN following the manufacturers’ instructions. Sequencing libraries were prepared by polyA selection, and sequencing was performed using NovaSeq (2×150bp, 20 million pair-end reads) by Genewiz from Azenta Life Sciences (Germany).

Paired-end reads were aligned to the human reference genome (GRCh38) using HISAT, and transcripts were annotated following the StringTie pipeline^83^. Raw counts generated from this workflow were used for downstream analysis. Genes with fewer than five counts across samples were excluded. DEGs were identified using DESeq2 (v1.22.2)^84^, with size factors calculated for normalization. DEGs were defined by an adjusted false discovery rate (FDR) < 0.05 and |log2 fold change| > 1. Raw data is deposited at EGA with ID: EGAD50000002099.

### Survival analyses

Bulk RNA-seq expression data and matched clinical information from the TCGA^53^, GLASS^25^ and Spitzer^54^ GB cohorts were processed to evaluate overall survival and progression-free interval in relation to the Edge-unique (Table S11) and OSM-activation (Table S13) gene signatures. Gene alias mapping was performed to maximize the correspondence between the gene signature and previously published datasets, accounting for outdated or non-standard gene symbols. For each cohort, per-sample signature scores were computed as the mean expression of constituent genes and patients were stratified into high- or low-signature groups using cohort-specific medians. Kaplan–Meier curves were generated using the survminer package in R and group differences were assessed using Mantel–Cox test.

### Quantification and statistical analysis

All statistical analyses were performed using GraphPad Prism v10 (GraphPad Software) and R v4.3.2. The statistical tests, *n* values, and exact p values are indicated in the corresponding figure legends. For *in vitro* experiments, *n* refers to the number of biological replicates, each plated in technical triplicates unless otherwise specified. Data are presented as mean ± SD unless otherwise noted. Statistical significance was defined as p<0.05. Significance is indicated with asterisks (*) and defined as follows: ns, p>0.05; * p≤0.05; ** p<0.01; *** p<0.001; **** p<0.0001. Assumptions of the statistical tests (e.g., normal distribution, equal variance) were not formally tested, but parametric analyses were applied to datasets with comparable variance and distribution across replicates.

## Data availability

All sequence data generated and analyzed in this study has been deposited in EGA, under accession number EGAS50000001455. Accession to public data used are for Schmassmann^58^ GSE197543, TCGA (https://www.cbioportal.org/study/summary?id=gbm_tcga), GLASS (https://www.synapse.org/Synapse:syn26465623) and Spitzer^54^ GSE274546. Original western blot images have been deposited at Mendeley Data at DOI: 10.17632/6s94kxnhfw.2 and are publicly available as of the date of publication. Microscopy data reported in this paper will be shared by the lead contact upon request. The patient-derived cell cultures generated in this study will be made available upon reasonable request following approval by an internal review board, the completion of a Materials Transfer Agreement and after compensation for processing and shipping.

## Code availability

All code used in this work is available on https://github.com/shindowfeather/HGCC5000-2/tree/main. The code used to reproduce the presented analysis is indexed on Zenodo. All scripts run on jupyter notebooks are available as “.ipynb” files and scripts executed in the command line are available as “.sh” files. R scripts are available as “R”.

## ACKNOWLEDGEMENTS

We are grateful to the patients for donating their tissue to our HGCC sample collection. This work was supported by grants to L.U. from the Swedish Cancer Society (CAN 2018/777, 21 1518 Pj, 24 3723 Pj), the Swedish Research Council (2022-01114) and the Swedish Brain Foundation (FO2025-0221), to F.J.S. from the Swedish Cancer Society (24 3881 Pj), Sjöbergstiftelsen (2020-162) and the Swedish Research Council (2022-00969), and to X.C. from the Swedish Cancer Society (24 3484 Pj, 22 0491 JIA), the Swedish Research Council (2022-00658, 2024-03756) and Knut and Alice Wallenberg Foundation (KAW 2023.0046). WES was performed by the SNP&SEQ Technology Platform at Uppsala University. The facility is part of the National Genomics Infrastructure (NGI) Sweden and Science for Life Laboratory and supported by the Swedish Research Council and the Knut and Alice Wallenberg Foundation. The data handling was enabled by resources provided by the National Academic Infrastructure for Supercomputing in Sweden (NAISS) and the Swedish National Infrastructure for Computing (SNIC) at Uppsala Multidisciplinary Center for Advanced Computational Science (UPPMAX) partially funded by the Swedish Research Council (2022-06725, 2018-05973). We thank Dr. Verónica Rendo for providing helpful feedback on the manuscript.

## AUTHOR CONTRIBUTIONS

Conceptualization: L.U. Data curation: X.L., V.J., A.S. Formal analysis: I.N., X.L., V.J., N.M., A.S., P.X., L.U. Funding acquisition: X.C, F.J.S., L.U. Investigation: I.N., V.J., F.L., N.M., Y.D., I.U., T.B., M.R. Methodology: I.N., F.L., N.M., Y.D., I.U., T.B., M.R., L.U. Project administration: I.N., X.L., V.J., F.L, N.M., A.S., Y.D., I.U., T.B., P.X., M.R., L.U. Software: X.L. Resources: F.L., M.R. Supervision: I.N., N.M., Y.D., M.R., X.C., F.J.S., L.U. Validation: I.N., V.J. Vizualization: I.N., X.L., V.J., F.L., A.S., L.U. Writing – original draft: L.U. Writing – review and editing: I.N., X.L., V.J., F.L., A.S., X.C., F.J.S., L.U.

## EXTENDED DATA

**Extended Data Figure 1.**
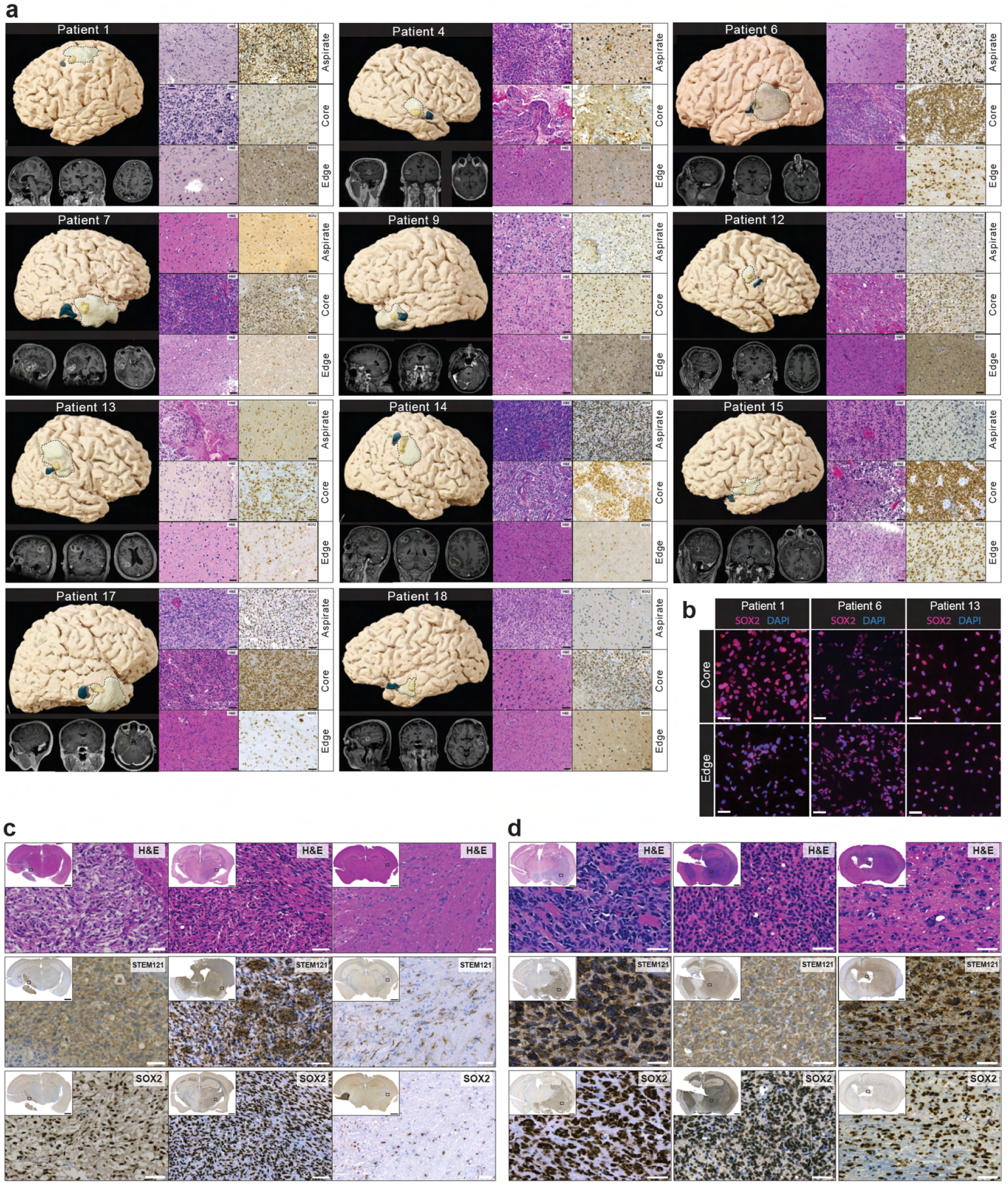
Characterization of the patient cohort and of patient-derived orthotopic xenografts (PDOXs). **(a)** Panels showing for each patient locations of specimens, histology and SOX2 expression. **Top left**, MRI-based sagittal illustration of sample locations reconstructed from T1+C sequences. Aspirate is outlined by a dashed black line, the annotated Core by a dashed red line, and the annotated Edge in blue. **Bottom left**, T1-weighted MRI images (left and middle - coronal, right - axial). **Middle**, H&E-stained sections. **Right**, SOX2 immunostainings. Scale bar, 50 µM. **(b)** Representative images of SOX2 immunostainings in matched Core and Edge cultures of patient 1, patient 6 and patient 13. Scale bar, 50 µM. **(c-d)** Examples of PDOXs generated after intracranial injections of freshly dissociated tissue samples or established GCCs. Sections were stained with H&E, or immunostained for STEM121 or SOX2. Scale bar, 1 mm for the inset and 50 µM. **(c)** Freshly dissociated tissue samples of Aspirate (patient 13, left column), Core (patient 12, middle column) and Edge (patient 12, right column). **(d)** Matched GCCs of patient 6, Aspirate (left column), Core (middle column) and Edge (right column).

**Extended Data Figure 2.**
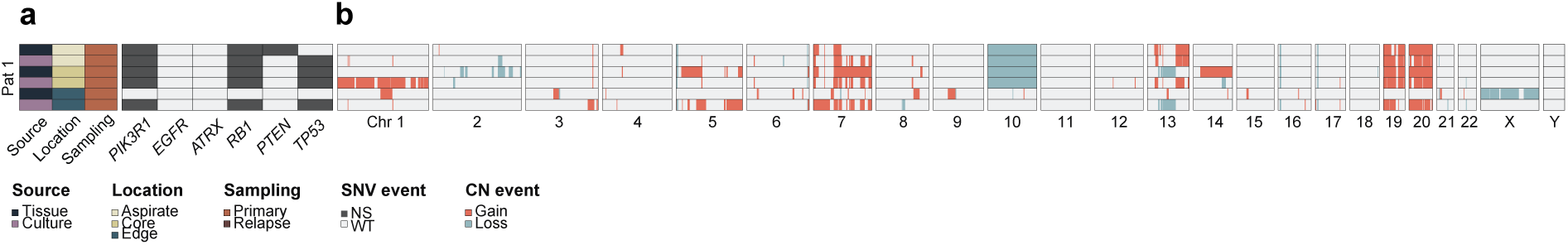
Tumor-only mode analyses of patient 1 WES data. **(a)** Oncoprint of GB relevant genes from *Wang et al.* in patient 1 across different sample locations. SNV, single nucleotide variant; NS, non-synonymous. **(b)** Copy Number (CN) variation profile of patient 1 across different sample locations.

**Extended Data Figure 3.**
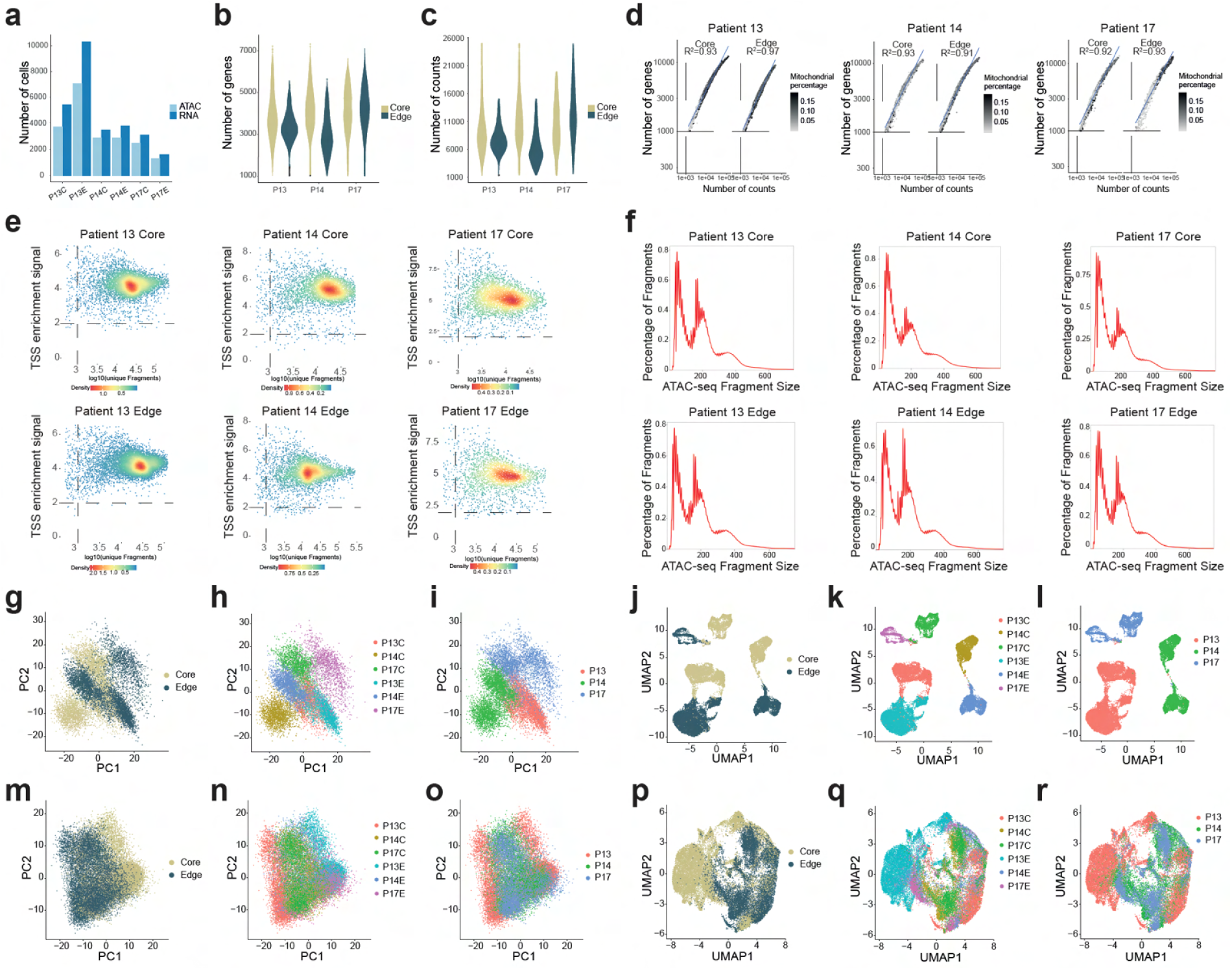
Preprocessing and quality control of the snRNA-seq and snATAC-seq data. **(a)** Number of cells that passed quality control (QC) for snRNA-seq (P13C n=4468, P14C n=2868, P17C n=2488, P13E n=8510, P14E n=3100, P17E n=1307), and snATAC-seq (P13C n=5403, P14C n=3499, P17C n=3083, P13E n=10243, P14E n=3811, P17E n=1631). **(b)** Total number of genes per cell in each snRNA-seq sample. Median number of genes in each sample is above 2000. **(c)** Number of counts per cell in each snRNA-seq sample, with a threshold of more than 1000. **(d)** Pearson correlation analysis between genes per cell and counts per cell. The percentage of mitochondrial RNA is indicated. **(e)** QC of snATAC-seq profiles (cells) using the thresholds counts per cell >1000 (y-axis) and TSS enrichment score >8 (x-axis), indicated with dotted lines. **(f)** Distribution of insert sizes between paired reads derived from sequencing of snATAC-seq libraries indicating nucleosome patterning. **(g-i)** PCA before batch-effect correction labeled by Core and Edge (**g**), sample (**h**), and individual patients (**i**). **(j-l)** UMAP before batch-effect correction labeled by Core and Edge (**j**), sample (**k**), and patients (**l**). **(m-o)** PCA after batch-effect correction labeled by Core and Edge (**m**), sample (**n**), and patients (**o**). **(p-r)** UMAP after batch-effect correction labeled by Core and Edge (**p**), sample (**q**), and patients (**r**).

**Extended Data Figure 4.**
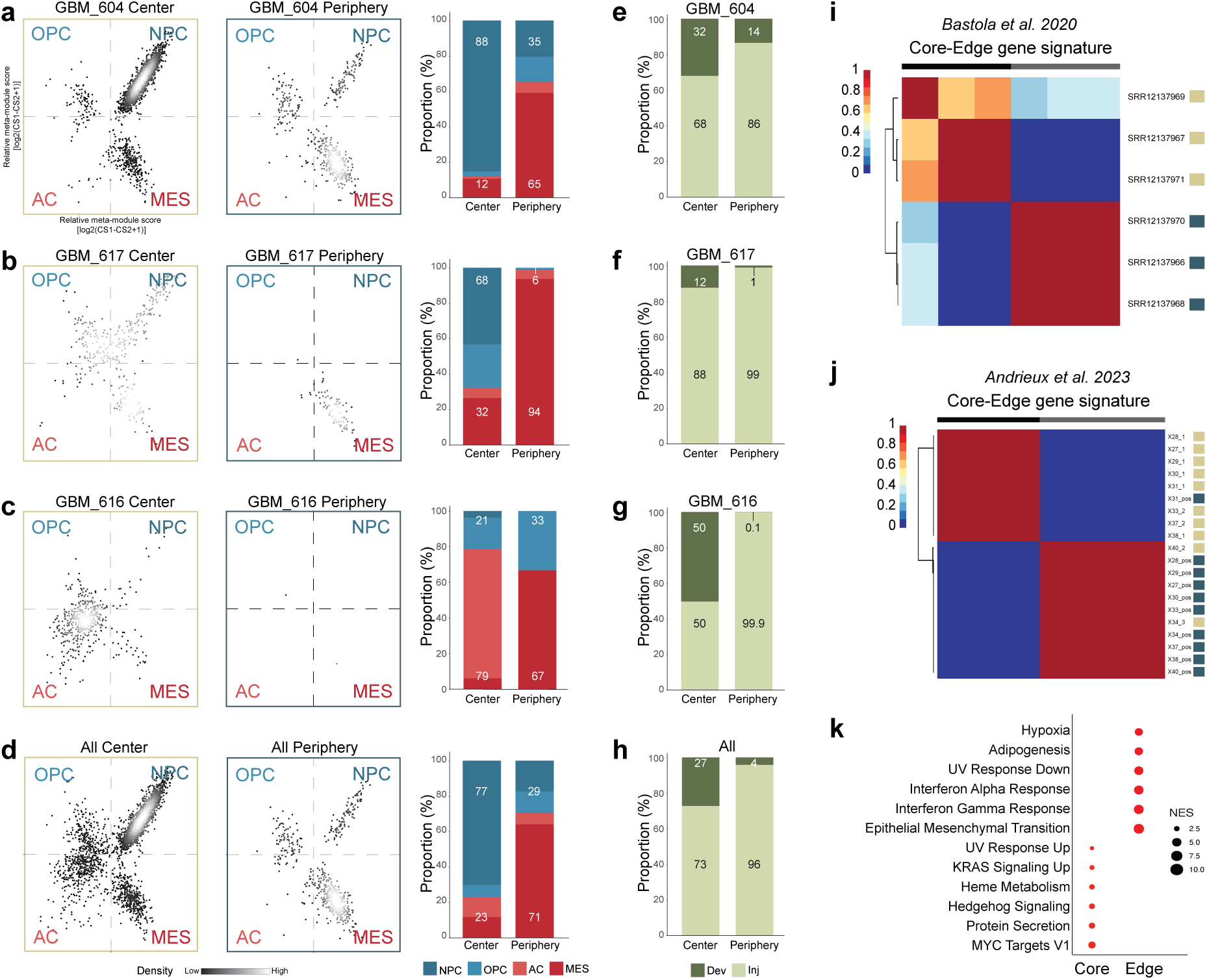
Cell state analyses of *Schmassman* Center and Periphery GB cells and validation of the Core-Edge gene signature in Bastola and Andrieux transcriptome data. **(a-d)** Visual representation of *Neftel* cell state heterogeneity of *Schmassmann* Center profiles (left) and Periphery profiles (middle), and percentage of combined NPC+OPC and AC+MES cell states (right) in individual patients (**a-c**) and all data combined (**d**). **(e-h)** Percentage of *Richards* Developmental (Dev) and Injury response (Inj) cell states in *Schmassmann* Center and Periphery profiles in individual patients (**e-g**) and all data combined (**h**). **(i-j)** NMF clustering using the Core-Edge gene signature on the *Andrieux* RNA-seq data of tumor core (yellow squares) and 5-ALA+ GB cells of invasive margin (blue squares) **(i)**, and the *Bastola* RNA-seq data of Core (yellow squares) and Edge (blue squares) samples **(j)**. **(k)** Gene set enrichment analysis (GSEA) scores for the Core-Edge gene signature with a false discovery rate (FDR) threshold of < 0.025.

**Extended Data Figure 5.**
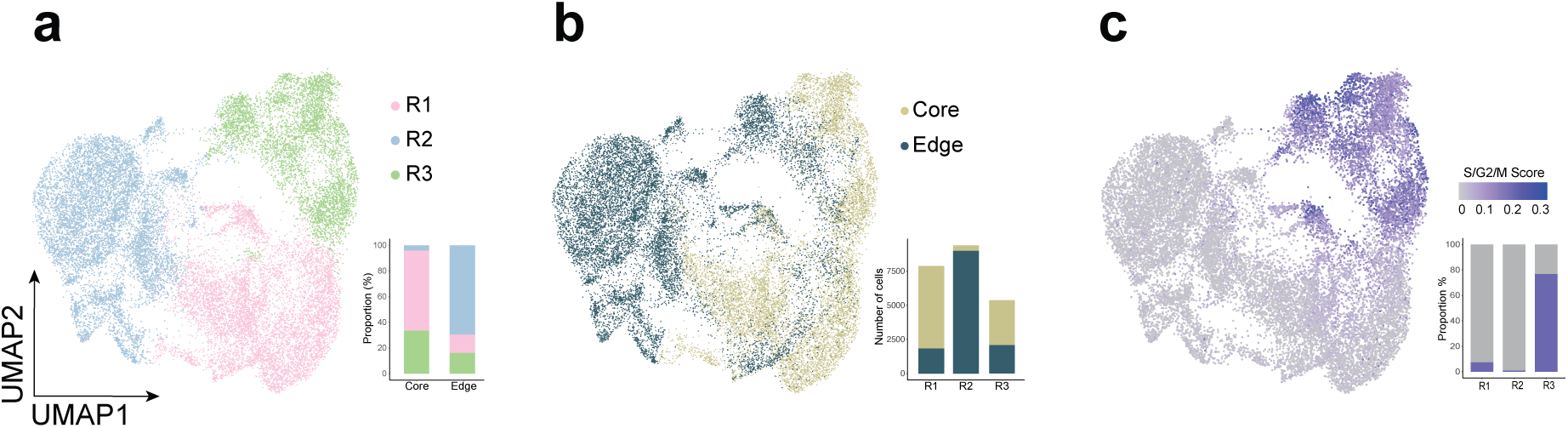
Clustering analysis of all nuclei with qualified dual (snRNA-seq and snATAC-seq) profiles. **(a)** UMAP representation performed on snRNA-seq profiles from nuclei qualified for both snRNA-seq and snATAC-seq (n=22741) showing the R1, R2, R3 clusters (left). Proportion of clusters in each sample (right). **(b)** Core and Edge cells projected in the snRNA UMAP (left). The absolute numbers of Core and Edge cells in each cluster (right). 1 (Core, n=6126; Edge, n=1845), 2 (Core, n=405; Edge, n=8998), 3 (Core, n=2074; Edge, n=3293). **(c)** Cycling cells projected in the snRNA UMAP (left). Proportion of cycling cells in the snRNA clusters (right).

**Extended Data Figure 6.**
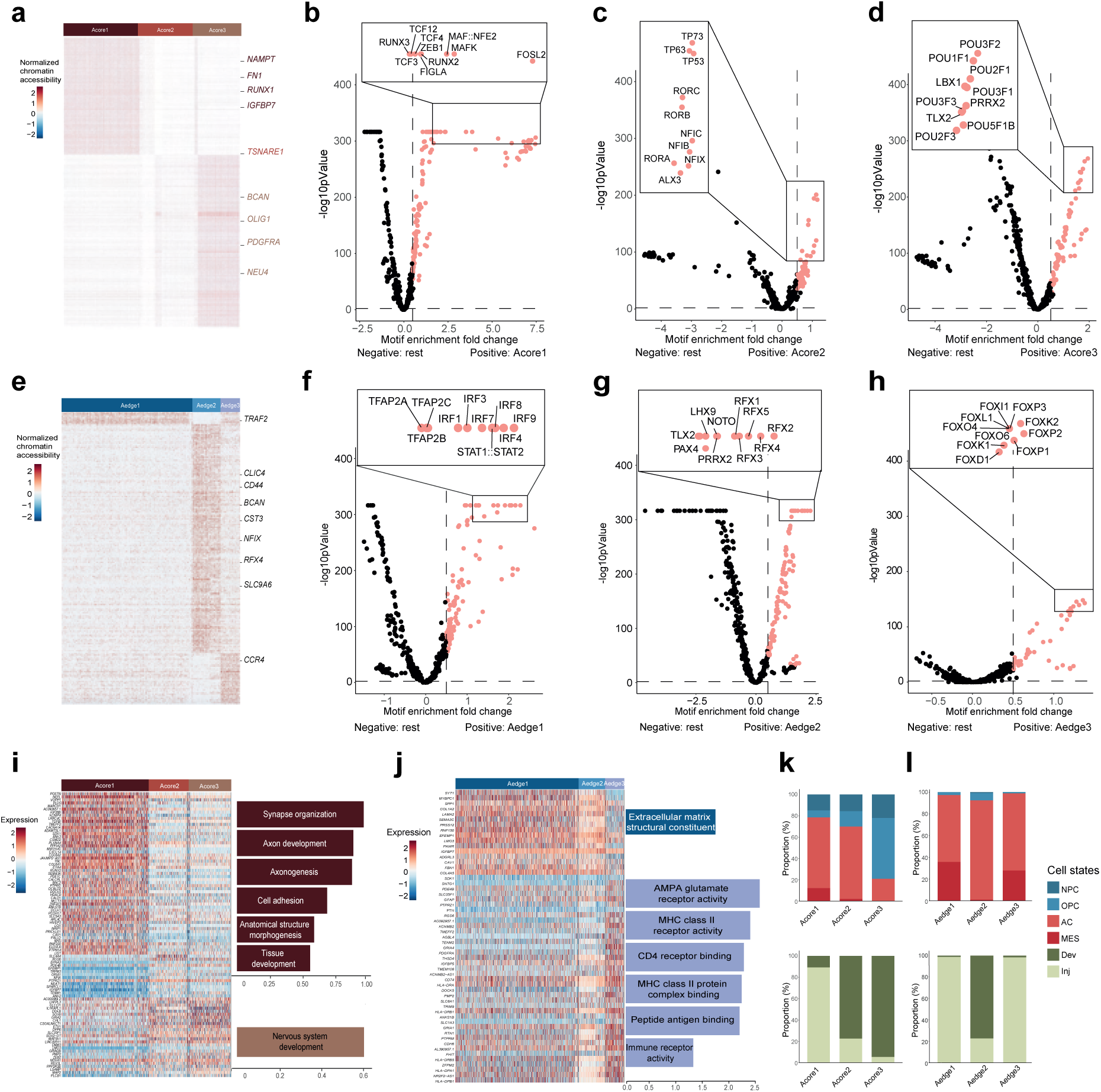
snATAC-based subcluster analysis of Core and Edge cells identifes distinct regulatory networks and tumor cell phenotypes. **(a)** Differential ATAC peaks in the Acore1, Acore2 and Acore3 subclusters. **(b-d)** Volcano plots of TF motif enrichment analysis of differential peaks in the snATAC Core subclusters. Threshold is set to p-value <0.01, log2FC>0.5, AUC>0.6. **(e)** Differential ATAC peaks of Aedge1, Aedge2, and Aedge3 subclusters. **(f-h)** Volcano plots of TF motif enrichment analysis of differential peaks in the snATAC Edge subclusters. Threshold is set to p-value <0.01, log2FC>0.5, AUC>0.6. **(i)** DEGs (left) and GO analysis (right) of Core subclusters. **(j)** DEGs (left) and GO analysis (right) of Edge subclusters. **(k-l)** Proportions of *Neftel* (top), and *Richards* (bottom) cell states in the Core subclusters (**k**) and Edge subclusters (**l**).

**Extended Data Figure 7.**
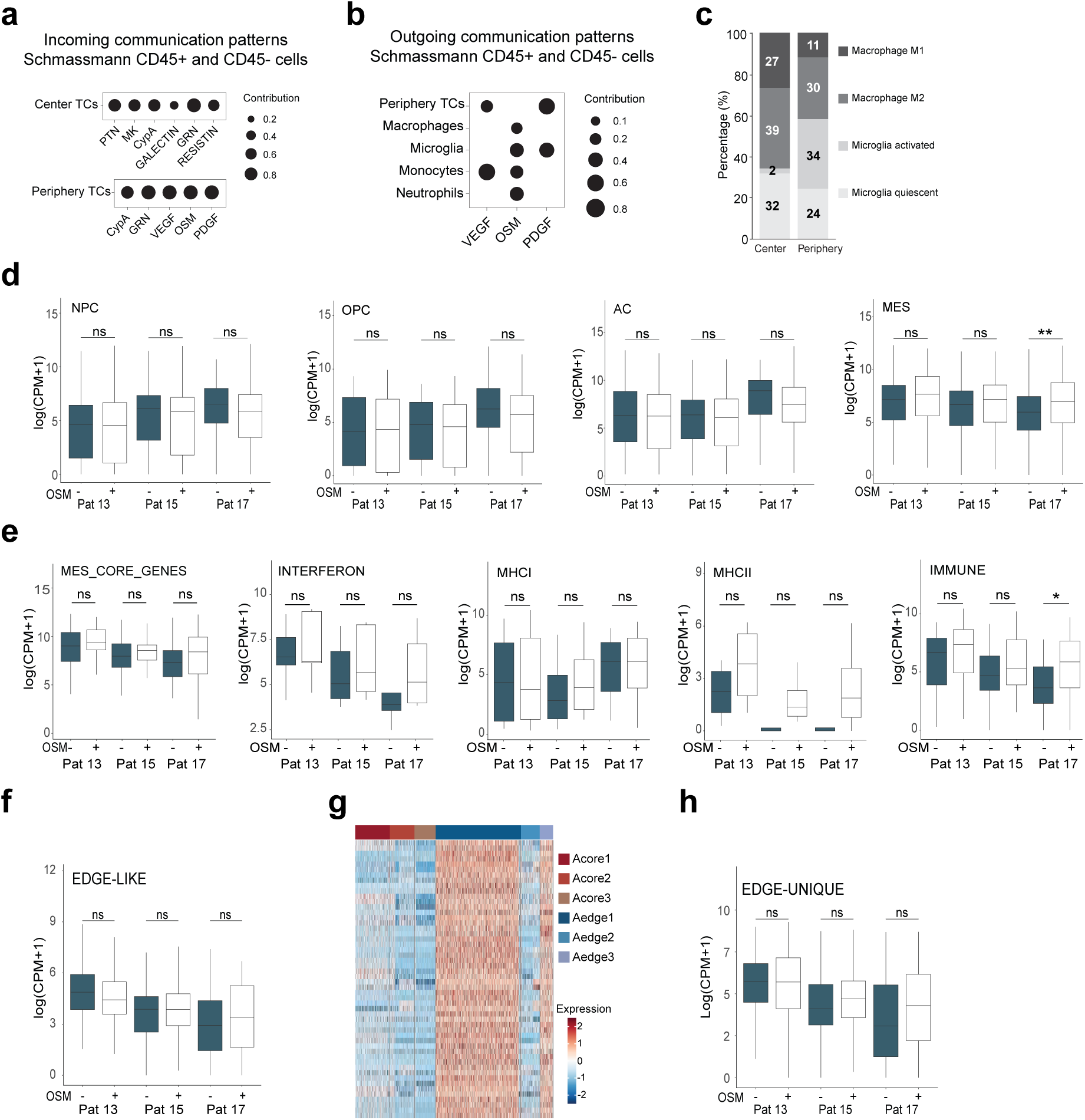
OSM-stimulation of Edge cells produces small changes in cell state and Edge-related gene expression. **(a)** CellChat incoming signaling pattern of scRNA profiles of *Schmassmann* CD45+ and CD45-cells, analyzed separately for Center (top) and Periphery (bottom). **(b)** CellChat outgoing signaling pattern of *Schmassmann* CD45+ and CD45- Periphery cells. **(c)** Proportions of M1/M2-polarized macrophages and activated/surveilling microglia in *Schmassmann* Center and Periphery samples relative to the total number of cells. **(d)** Expression of *Neftel* cell state gene signatures in RNA-seq data of Edge cells cultured in absence or presence of OSM. Mann-Whitney U test, **p = 0.0018. **(e)** Expression of *Chanoch-Myers* mesenchymal cell state gene signatures in RNA-seq data of Edge cells cultured in absence or presence of OSM. Mann-Whitney U test was performed, *p= 0.014. **(f)** Expression of the Edge gene signature (Table S5) in RNA-seq data of Edge cells cultured in absence or presence of OSM. Mann-Whitney U test was performed. **(g)** Expression of the Edge-unique gene signature in the snRNA profiles organized in Acore1-3 and Aedge1-3 subclusters. **(h)** Expression of the Edge-unique gene signature (Table S10) in RNA-seq data of Edge cells cultured in absence or presence of OSM. Mann-Whitney U test was performed.

