## Supplemental Table 1 for "Identification of an epigenetically and phenotypically distinct peritumoral glioblastoma cell population linked to inferior patient outcome"

**Table S1. The patient cohort**

| Patient ID | Sex | Age[1] (years) | Survival[2] (days) | Diagnosis[3] | Primary/ recurrent | Tumor location | Aspirate-derived culture | Core-derived culture | Edge-derived culture |
|---|---|---|---|---|---|---|---|---|---|
| 1 | F | 68 | 123 | Glioblastoma | Primary | Frontal left | U5000A | U5001C | U5002E |
| 4 | M | 68 | 321 | Glioblastoma | Primary | Temporal right | U5007A | FAIL | FAIL |
| 6 | M | 72 | 765 | Glioblastoma | Primary | Temporal left | U5013A | U5014C | U5015E |
| 7 | M | 69 | 634 | Glioblastoma | Primary | Temporal right | U5016A | FAIL | FAIL |
| 9 | F | 67 | 385 | Glioblastoma | Primary | Temporal left | U5022A | U5023C | FAIL |
| 12 | M | 72 | 20 | Glioblastoma | Primary | Frontal right | U5030A | U5031C | U5032E to p7 |
| 13 | M | 58 | 219 | Glioblastoma | Primary | Frontal right | U5033A | U5034C | U5035E |
| 14 | M | 62 | 2150 | Glioblastoma | Primary | Parietal right | U5036A | U5037C | U5038E |
| 15 | F | 62 | 292 | Glioblastoma | Primary | Temporal left | U5039A | U5040C | U5041E |
| 17 | F | 46 | 1300 | Glioblastoma | Primary | Temporal right | U5045A | U5046C | U5047E |
| | | N/A | 362 | N/A | Recurrent | Temporal right | U5045A_R | N/A | N/A |
| 18 | M | 66 | 413 | Glioblastoma | Primary | Temporal left | FAIL | FAIL | FAIL |

[1]at surgery, [2]from date of surgery, [3]WHO 2021 classification
