## Supplemental Table 2 for "Identification of an epigenetically and phenotypically distinct peritumoral glioblastoma cell population linked to inferior patient outcome"

**Table S2. STR profiling of the parental tumor and cell cultures derived thereof**

| | Marker | D8S1179 | D21S11 | D7S820 | CSF1PO | D3S1358 | TH01 | D13S317 | D16S539 | D2S1338 | D19S433 | vWA | TPOX | D18S51 | AMEL | D5S818 | FGA |
|---|---|---|---|---|---|---|---|---|---|---|---|---|---|---|---|---|---|
| **Patient** | **Sample** | | | | | | | | | | | | | | | | |
| **1** | **Aspirate tissue** | 12,14 | 30,31 | 10,11 | 11,11 | 15,18 | 7,8 | 10,11 | 8,11 | 17,20 | 14,14 | 14,17 | 8,10 | 13,17 | X,X | 11,13 | 24,25 |
| | **Aspirate culture** | 12,14 | 30,31 | 10,11 | 11,11 | 15,18 | 7,8 | 10,11 | 8,11 | 17,20 | 14,14 | 14,17 | 8,10 | 13,17 | X,X | 11,13 | 24,25 |
| | **Core culture** | 12,14 | 30,31 | 10,11 | 11,11 | 15,18 | 7,8 | 10,11 | 8,11 | 17,20 | 14,14 | 14,17 | 8,10 | 13,17 | X,X | 11,13 | 24,25 |
| | **Edge culture** | 12,14 | 30,31 | 10,11 | 11,11 | 15,18 | 7,8 | 10,11 | 8,11 | 17,20 | 14,14 | 14,17 | 8,10 | 13,17 | X,X | 11,13 | 24,25 |
| **6** | **Aspirate tissue** | 12,13 | 28,30 | 7,11 | 10,11 | 16,16 | 7,9.3 | 11,12 | 12,12 | 19,24 | 14,15 | 17,19 | 8,12 | 12,13 | X,Y | 12,13 | 20,20 |
| | **Aspirate culture** | 12,13 | 28,30 | 7,11 | 10,11 | 16,16 | 7,9.3 | 11,11 | 12,12 | 19,24 | 14,15 | 17,19 | 8,12 | 12,13 | X,Y | 12,13 | 20,20 |
| | **Core culture** | 12,13 | 28,30 | 7,11 | 10,11 | 16,16 | 7,9.3 | 11,11 | 12,12 | 19,24 | 14,15 | 17,19 | 8,12 | 12,13 | X,Y | 12,13 | 20,20 |
| | **Edge culture** | 12,13 | 28,30 | 7,11 | 10,11 | 16,16 | 7,9.3 | 11,11 | 12,12 | 19,24 | 14,15 | 17,19 | 8,12 | 12,13 | X,Y | 12,13 | 20,20 |
| **13** | **Aspirate tissue** | 8,13 | 28,34.2 | 11,12 | 11,12 | 16,18 | 7,9.3 | 11,14 | 12,13 | 20,20 | 13,15 | 17,18 | 8,9 | 16,18 | X,Y | 12,12 | 19,22 |
| | **Aspirate culture** | 8,13 | 28,34.2 | 11,12 | 11,12 | 16,18 | 7,9.3 | 11,14 | 12,13 | 20,20 | 13,15 | 17,18 | 8,9 | 16,18 | X,Y | 12,12 | 19,22 |
| | **Core culture** | 8,13 | 28,34.2 | 11,12 | 11,12 | 16,18 | 7,9.3 | 11,14 | 12,13 | 20,20 | 13,15 | 17,18 | 8,9 | 16,18 | X,Y | 12,12 | 19,22 |
| | **Edge culture** | 8,13 | 28,34.2 | 11,12 | 11,12 | 16,18 | 7,9.3 | 14,14 | 12,13 | 20,20 | 13,15 | 17,18 | 8,9 | 16,16 | X,Y | 12,12 | 19,22 |
| **14** | **Aspirate tissue** | 14,16 | 30,31.2 | 9,10 | 9,10 | 15,17 | 7,9.3 | 11,12 | 9,11 | 24,25 | 14,16 | 17,19 | 10,12 | 13,18 | X,Y | 11,12 | 18,22.2 |
| | **Aspirate culture** | 14,16 | 30,31.2 | 9,10 | 9,10 | 15,17 | 9.3,9.3 | 11,11 | 9,11 | 24,25 | 14,16 | 17,19 | 10,12 | 13,18 | X,Y | 11,12 | 18,22.2 |
| | **Core culture** | 14,16 | 30,31.2 | 9,10 | 9,10 | 15,17 | 9.3,9.3 | 11,11 | 9,11 | 24,25 | 14,16 | 17,19 | 10,12 | 13,18 | X,Y | 11,12 | 18,22.2 |
| | **Edge culture** | 14,16 | 30,31.2 | 9,10 | 9,10 | 15,17 | 7,9.3 | 11,11 | 9,11 | 24,25 | 14,16 | 17,19 | 10,12 | 13,18 | X,Y | 11,12 | 18,22.2 |
| **15** | **Aspirate tissue** | 13,16 | 29,29 | 9,10 | 10,11 | 16,16 | 9.3,9.3 | 11,11 | 9,9 | 19,23 | 12.2,14 | 16,17 | 8,11 | 12,13 | X,X | 12,13 | 20,20 |
| | **Aspirate culture** | 13,16 | 29,29 | 9,10 | 10,11 | 16,16 | 9.3,9.3 | 11,11 | 9,9 | 19,23 | 12.2,14 | 16,17 | 8,11 | 12,13 | X,X | 12,13 | 20,20 |
| | **Core culture** | 13,16 | 29,29 | 9,10 | 10,11 | 16,16 | 9.3,9.3 | 11,11 | 9,9 | 19,23 | 12.2,14 | 16,17 | 8,11 | 12,13 | X,X | 12,13 | 20,20 |
| | **Edge culture** | 13,16 | 29,29 | 9,10 | 10,11 | 16,16 | 9.3,9.3 | 11,11 | 9,9 | 19,23 | 12.2,14 | 16,17 | 8,11 | 12,13 | X,X | 12,13 | 20,20 |
| **17** | **Aspirate tissue** | 10,13 | 29,29 | 9,10 | 11,12 | 15,16 | 9.3,9.3 | 11,12 | 11,11 | 25,25 | 13,13 | 16,17 | 8,11 | 13,18 | X,X | 12,13 | 19,24 |
| | **Aspirate culture** | 10,13 | 29,29 | 9,10 | 11,12 | 15,16 | 9.3,9.3 | 11,12 | 11,11 | 25,25 | 13,13 | 16,17 | 8,11 | 13,18 | X,X | 12,13 | 19,24 |
| | **Core culture** | 10,13 | 29,29 | 9,9 | 11,12 | 15,16 | 9.3,9.3 | 11,12 | 11,11 | 25,25 | 13,13 | 16,17 | 8,11 | 13,18 | X,X | 12,13 | 19,24 |
| | **Edge culture** | 10,13 | 29,29 | 9,10 | 11,12 | 15,16 | 9.3,9.3 | 11,12 | 11,11 | 25,25 | 13,13 | 16,17 | 8,11 | 13,18 | X,X | 12,13 | 19,24 |
