## Supplemental Table 3 for "Identification of an epigenetically and phenotypically distinct peritumoral glioblastoma cell population linked to inferior patient outcome"

**Table S3. Evaluation of orthotopic tumor cell growth**

| Patient ID | IC injected Aspirate cells | | | IC injected Core cells | | | IC injected Edge cells | | |
|---|---|---|---|---|---|---|---|---|---|
| | No mice | IC tumor | IC cells | No mice | IC tumor | IC cells | No mice | IC tumor | IC cells |
| 1 | 3 | + | N/A | 4 | + | N/A | 6 | - | - |
| 6 | 4 | + | N/A | 5 | + | N/A | 6 | + | N/A |
| 13 | 3 | + | N/A | 9 | + | N/A | 9 | - | + |
| 14 | 5 | + | N/A | 10 | + | N/A | 11 | - | + |
| 15 | 4 | + | N/A | 6 | - | + | 5 | + | N/A |
| 17 | 4 | + | N/A | 9 | - | + | 9 | - | + |

**Note**: IC = intracranial, no = number of, IC tumor + = tumor detected in ≥ one brain, IC cells + = tumor cells detected in ≥ one brain
