## Supplemental Table 4 for "Identification of an epigenetically and phenotypically distinct peritumoral glioblastoma cell population linked to inferior patient outcome"

**Table S4. Subclonal fractions in Aspirate, Core, Edge and recurrent Aspirate tumor samples**

| | | Aspirate | Core | Edge | Recurrent Aspirate |
|---|---|---|---|---|---|
| Patient 6 | 1 | 0,1328 | 0,162987 | 0,91 | ND |
| | 2 | 0,00235001 | 1,0126E-09 | 0,0330634 | ND |
| | 3 | 0,565476 | 1,5638E-10 | 0,0108787 | ND |
| | 4 | 2,36545E-05 | 0,567237 | 0,000021348 | ND |
| | 5 | 0,238777 | 0,138875 | 2,28862E-05 | ND |
| | 6 | 0,000023164 | 0,1309 | 0,000013697 | ND |
| | 7 | 0,06055 | 0 | 0,046 | ND |
| Patient 13 | 1 | 0,0173 | 0,5531 | 0,7724 | ND |
| | 2 | 0,258267 | 0,296833 | 0,0149333 | ND |
| | 3 | 0,272867 | 0,0673 | 0,100533 | ND |
| | 4 | 0,405567 | 0,0468 | 0,0587 | ND |
| | 5 | 0,046 | 0,0359667 | 0,0534333 | ND |
| Patient 14 | 1 | 0,0862 | 0,4517 | 0,747233 | ND |
| | 2 | 1,31127E-09 | 0,0418 | 0,23585 | ND |
| | 3 | 0,171967 | 0,0325 | 2,89504E-05 | ND |
| | 4 | 0,3559 | 0,4205 | 0,00002094 | ND |
| | 5 | 0,3625 | 0,0308 | 0 | ND |
| | 6 | 0,0234333 | 0,0227 | 0,0168667 | ND |
| Patient 15 | 1 | 0,2557 | 0,362933 | 0,145933 | ND |
| | 2 | 0,2057 | 0,0783667 | 0,313217 | ND |
| | 3 | 0,49202 | 0,5028 | 0,49481 | ND |
| | 4 | 0,04658 | 0,0559 | 0,04604 | ND |
| Patient 17 | 1 | 0,2073 | 0,0965 | 0,6273 | 0,2626 |
| | 2 | 0,06025 | 0,12715 | 0,0137 | 0,2057 |
| | 3 | 7,55015E-10 | 3,5859E-11 | 0,03515 | 0,208876 |
| | 4 | 0,43455 | 0,38965 | 8,89337E-11 | 9,0552E-11 |
| | 5 | 0,291167 | 0,351867 | 0,0480833 | 0,03015 |
| | 6 | 0,00673333 | 0,0348333 | 0,0134667 | 0,0303 |
| | 7 | 7,55014E-10 | 3,5859E-11 | 0,2522 | 7,379E-05 |
| | 8 | 0 | 0 | 0,0101 | 0,2623 |
