## Supplemental Table 5 for "Identification of an epigenetically and phenotypically distinct peritumoral glioblastoma cell population linked to inferior patient outcome"

**Table S5. The Core-Edge gene signature. Differentially expressed genes in the Core-like and Edge-like clusters**

| Gene | Cluster | RNA average expression | Log2 fold-change | AUC | p-val | FDR |
|---|---|---|---|---|---|---|
| COL3A1 | Edge-like | 0,893902286 | 0,690106919 | 0,719899818 | 0 | 0 |
| FN1 | Edge-like | 1,083443125 | 0,536155353 | 0,70676366 | 0 | 0 |
| PDE1A | Edge-like | 0,872086607 | 0,711009947 | 0,664465017 | 0 | 0 |
| ROBO2 | Edge-like | 1,058393581 | 0,557127407 | 0,62809218 | 1,38E-232 | 1,76E-232 |
| ACTG2 | Edge-like | 0,85466481 | 0,622600328 | 0,711591081 | 0 | 0 |
| CHSY3 | Edge-like | 0,954026914 | 0,682353385 | 0,688917898 | 0 | 0 |
| GABRG3 | Edge-like | 1,491704066 | 1,062168743 | 0,838043322 | 0 | 0 |
| ARHGAP24 | Edge-like | 0,914164127 | 0,576963625 | 0,663788947 | 0 | 0 |
| MYBPC1 | Edge-like | 1,104957193 | 0,891375582 | 0,745322854 | 0 | 0 |
| EPHA3 | Edge-like | 1,68851208 | 1,387737612 | 0,820913817 | 0 | 0 |
| SPP1 | Edge-like | 1,784180413 | 1,056906084 | 0,816686259 | 0 | 0 |
| THBS1 | Edge-like | 1,036594617 | 0,730491513 | 0,747093272 | 0 | 0 |
| PCDH7 | Edge-like | 1,322932755 | 0,649133331 | 0,71379623 | 0 | 0 |
| KCNK2 | Edge-like | 0,859111308 | 0,641725008 | 0,683529195 | 0 | 0 |
| PI15 | Edge-like | 0,666984647 | 0,505325877 | 0,656720831 | 0 | 0 |
| COL1A2 | Edge-like | 2,000423358 | 1,111960841 | 0,842748028 | 0 | 0 |
| THSD4 | Edge-like | 0,723077751 | 0,501114437 | 0,669768691 | 0 | 0 |
| KIRREL3 | Edge-like | 0,917018064 | 0,546537312 | 0,644285301 | 4,68E-297 | 6,23E-297 |
| LAMA2 | Edge-like | 1,494388723 | 1,125751049 | 0,810793955 | 0 | 0 |
| PRSS12 | Edge-like | 0,863799859 | 0,704923612 | 0,732586263 | 0 | 0 |
| ITGBL1 | Edge-like | 1,426309197 | 1,02400821 | 0,84282906 | 0 | 0 |
| ALPK2 | Edge-like | 1,10906831 | 0,595467039 | 0,674585847 | 0 | 0 |
| ITPR2 | Edge-like | 1,012058403 | 0,516934392 | 0,69188286 | 0 | 0 |
| RNF150 | Edge-like | 1,49375843 | 1,039699347 | 0,823835734 | 0 | 0 |
| CPA6 | Edge-like | 0,88357423 | 0,640052038 | 0,62622507 | 9,73E-231 | 1,24E-230 |
| EFEMP1 | Edge-like | 2,003963459 | 1,529967839 | 0,927347282 | 0 | 0 |
| PRUNE2 | Edge-like | 2,046863835 | 1,078725582 | 0,8432883 | 0 | 0 |
| COL21A1 | Edge-like | 2,470314477 | 1,438685593 | 0,892440832 | 0 | 0 |
| PDE1C | Edge-like | 2,5329095 | 0,767676384 | 0,697003254 | 0 | 0 |
| BICC1 | Edge-like | 1,312023554 | 0,924886532 | 0,816460159 | 0 | 0 |
| LMO3 | Edge-like | 1,574026904 | 1,219802235 | 0,883719529 | 0 | 0 |
| CD74 | Edge-like | 1,094589743 | 0,729734429 | 0,792993052 | 0 | 0 |
| TRPM3 | Edge-like | 1,723825507 | 1,072287539 | 0,782619296 | 0 | 0 |
| HLA-DRA | Edge-like | 1,299469513 | 0,684739993 | 0,772462205 | 0 | 0 |
| NTRK2 | Edge-like | 1,360065264 | 1,020376551 | 0,829043518 | 0 | 0 |
| GRIN2B | Edge-like | 1,396621131 | 0,599082146 | 0,663229332 | 0 | 0 |
| NR3C2 | Edge-like | 0,901397203 | 0,656405251 | 0,708538438 | 0 | 0 |
| TGFB2 | Edge-like | 1,224886484 | 0,551538386 | 0,677668676 | 0 | 0 |
| TMTC2 | Edge-like | 1,621085722 | 0,655457729 | 0,652211726 | 0 | 0 |
| NAV3 | Edge-like | 1,157200091 | 0,526013225 | 0,675504654 | 0 | 0 |
| SLC7A11 | Edge-like | 0,951436448 | 0,53928699 | 0,674837236 | 0 | 0 |
| HLA-DRB1 | Edge-like | 1,030077819 | 0,666038043 | 0,77177522 | 0 | 0 |
| CALD1 | Edge-like | 3,60163454 | 0,818456834 | 0,769162617 | 0 | 0 |
| LTBP1 | Edge-like | 1,786349631 | 0,951371262 | 0,799100238 | 0 | 0 |
| NFIA | Edge-like | 1,709530999 | 0,86383069 | 0,773357937 | 0 | 0 |
| COL4A6 | Edge-like | 0,641051379 | 0,509564113 | 0,658431617 | 0 | 0 |
| GRIA1 | Edge-like | 1,088767375 | 0,801871672 | 0,68647265 | 0 | 0 |
| PAWR | Edge-like | 0,7340403 | 0,534555881 | 0,679664594 | 0 | 0 |
| AL096854.1 | Edge-like | 1,092616251 | 0,751865107 | 0,791349251 | 0 | 0 |
| NNMT | Edge-like | 1,138687124 | 0,845395863 | 0,839276439 | 0 | 0 |

| | | | | | | |
|---|---|---|---|---|---|---|
| AHR | Edge-like | 1,994257558 | 0,991112208 | 0,849958367 | 0 | 0 |
| NEAT1 | Edge-like | 3,188701432 | 2,004385749 | 0,93415044 | 0 | 0 |
| GCNT1 | Edge-like | 1,235970823 | 0,6652647 | 0,733170839 | 0 | 0 |
| PDE4D | Edge-like | 2,773239326 | 0,84105934 | 0,739175835 | 0 | 0 |
| SYNM | Edge-like | 0,992539434 | 0,681979609 | 0,755420569 | 0 | 0 |
| CTNNA2 | Edge-like | 2,115498071 | 0,770268829 | 0,720360622 | 0 | 0 |
| SPARCL1 | Edge-like | 1,769044963 | 0,917101659 | 0,81899406 | 0 | 0 |
| IGFBP7 | Edge-like | 3,011071896 | 1,785557945 | 0,951407818 | 0 | 0 |
| AL390957.1 | Edge-like | 1,109684444 | 0,729233433 | 0,763326786 | 0 | 0 |
| PRICKLE1 | Edge-like | 2,137518208 | 0,583356291 | 0,74780805 | 0 | 0 |
| PALLD | Edge-like | 2,309157105 | 0,703451196 | 0,738522818 | 0 | 0 |
| TENM1 | Edge-like | 1,710365399 | 1,048526432 | 0,813578052 | 0 | 0 |
| MAP2 | Edge-like | 2,580191554 | 0,760921959 | 0,751070628 | 0 | 0 |
| FBN1 | Edge-like | 1,188118107 | 0,784320938 | 0,811397739 | 0 | 0 |
| CCDC102B | Edge-like | 1,291843008 | 0,534937648 | 0,648429613 | 8,66E-307 | 1,16E-306 |
| CRIM1 | Edge-like | 1,548239794 | 0,727780589 | 0,723673266 | 0 | 0 |
| SCG2 | Edge-like | 1,755724851 | 0,543352272 | 0,674140806 | 0 | 0 |
| AC090371.2 | Edge-like | 0,737058128 | 0,580625782 | 0,745902143 | 0 | 0 |
| ARHGAP29 | Edge-like | 1,26440658 | 0,545372957 | 0,671469019 | 0 | 0 |
| ABCC3 | Edge-like | 1,061267317 | 0,587678941 | 0,714938463 | 0 | 0 |
| ANK2 | Edge-like | 2,791494308 | 1,660081138 | 0,914803867 | 0 | 0 |
| PLCB1 | Edge-like | 1,60398135 | 0,806149826 | 0,750254799 | 0 | 0 |
| COL4A5 | Edge-like | 1,649838128 | 1,255793643 | 0,87291507 | 0 | 0 |
| DCC | Core-like | 0,800144038 | 0,739216909 | 0,849698299 | 0 | 0 |
| AC002069.2 | Core-like | 0,837616519 | 0,577696168 | 0,738928703 | 0 | 0 |
| GRIK1 | Core-like | 1,357859161 | 0,731573674 | 0,721878374 | 0 | 0 |
| LRRTM4 | Core-like | 0,586070067 | 0,526451915 | 0,796916564 | 0 | 0 |
| NEFL | Core-like | 1,335330845 | 1,237821204 | 0,93916175 | 0 | 0 |
| NXPH1 | Core-like | 1,29504609 | 1,173203135 | 0,859085474 | 0 | 0 |
| PLD5 | Core-like | 1,198504224 | 0,863513153 | 0,822699429 | 0 | 0 |
| GRIK2 | Core-like | 1,076186655 | 0,743482219 | 0,786038937 | 0 | 0 |
| ZFPM2-AS1 | Core-like | 0,679848832 | 0,583320402 | 0,832831577 | 0 | 0 |
| AC092957.1 | Core-like | 1,692237141 | 0,853357033 | 0,752527525 | 0 | 0 |
| CENPF | Core-like | 0,944406328 | 0,576411613 | 0,709220646 | 0 | 0 |
| LRRC4C | Core-like | 1,566410734 | 1,255508386 | 0,848590872 | 0 | 0 |
| TGFBI | Core-like | 0,834656722 | 0,765693561 | 0,913787705 | 0 | 0 |
| CADM2 | Core-like | 0,799422948 | 0,502374416 | 0,731308864 | 0 | 0 |
| CACNA1A | Core-like | 1,800620187 | 1,203096389 | 0,83089232 | 0 | 0 |
| SDK1 | Core-like | 2,203380643 | 1,238397852 | 0,824284377 | 0 | 0 |
| ANK3 | Core-like | 1,173612764 | 0,562227741 | 0,697197215 | 0 | 0 |
| PPFIA2 | Core-like | 1,079507837 | 0,691608491 | 0,779088075 | 0 | 0 |
| CSMD2 | Core-like | 0,908944377 | 0,809237679 | 0,845724084 | 0 | 0 |
| PLXNA4 | Core-like | 1,126082837 | 0,920020599 | 0,844806622 | 0 | 0 |
| NKAIN3 | Core-like | 2,021911906 | 0,767799912 | 0,69514477 | 0 | 0 |
| SNTG1 | Core-like | 1,867977868 | 1,258513287 | 0,822072305 | 0 | 0 |
| JAKMIP2-AS1 | Core-like | 1,358416918 | 0,835713813 | 0,793077229 | 0 | 0 |
| COL6A2 | Core-like | 1,621270073 | 0,765335895 | 0,797296574 | 0 | 0 |
| HIP1 | Core-like | 1,008383045 | 0,519063514 | 0,719164774 | 0 | 0 |
| ARL6IP1 | Core-like | 1,038531199 | 0,583249767 | 0,733561122 | 0 | 0 |
| TNC | Core-like | 1,307597927 | 0,821992417 | 0,794455236 | 0 | 0 |
| EYA4 | Core-like | 1,188216099 | 0,623598887 | 0,743596722 | 0 | 0 |
| COL6A1 | Core-like | 1,701794838 | 0,763439716 | 0,762768673 | 0 | 0 |
| AL117329.1 | Core-like | 0,585619602 | 0,505633041 | 0,850296873 | 0 | 0 |
| LINC00511 | Core-like | 1,110847984 | 0,703329723 | 0,774188453 | 0 | 0 |
| NAV2 | Core-like | 0,81875186 | 0,519652162 | 0,745088518 | 0 | 0 |
| SOX4 | Core-like | 1,112697276 | 0,711176389 | 0,762932488 | 0 | 0 |

| | | | | | | |
|---|---|---|---|---|---|---|
| MMP16 | Core-like | 1,537950764 | 0,818149639 | 0,763733567 | 0 | 0 |
| FREM2 | Core-like | 0,995823177 | 0,735536707 | 0,798827862 | 0 | 0 |
| PTPRG | Core-like | 1,64919649 | 0,56052114 | 0,671364002 | 0 | 0 |
| NCAM1 | Core-like | 0,80736724 | 0,569862671 | 0,792193582 | 0 | 0 |
| NOVA1 | Core-like | 1,841839455 | 0,810074864 | 0,766135155 | 0 | 0 |
| HDAC9 | Core-like | 1,196675376 | 0,733458826 | 0,776017384 | 0 | 0 |
| TMEM158 | Core-like | 0,642665205 | 0,502182102 | 0,811318938 | 0 | 0 |
| PTPRZ1 | Core-like | 2,095181678 | 0,61641383 | 0,690285213 | 0 | 0 |
| MLLT3 | Core-like | 0,921119659 | 0,575432173 | 0,758549799 | 0 | 0 |
| LDLRAD3 | Core-like | 0,969461441 | 0,69937832 | 0,809546499 | 0 | 0 |
| SLC1A3 | Core-like | 1,152283232 | 0,661089965 | 0,756365699 | 0 | 0 |
| PCDH17 | Core-like | 0,915427405 | 0,654164977 | 0,801928147 | 0 | 0 |
| NEDD4L | Core-like | 0,820168209 | 0,577891721 | 0,784603758 | 0 | 0 |
| PTN | Core-like | 2,52305109 | 1,279566617 | 0,872278201 | 0 | 0 |
| ARL4C | Core-like | 0,929418315 | 0,661603159 | 0,812426764 | 0 | 0 |
| CASC15 | Core-like | 1,028716773 | 0,713596847 | 0,791476165 | 0 | 0 |

Note: AUC = Area under curve, FDR = False discovery rate
