## Supplemental Table 6 for "Identification of an epigenetically and phenotypically distinct peritumoral glioblastoma cell population linked to inferior patient outcome"

**Table S6. Peak annotation in the Core-like R1 and Edge-like R2 clusters**

| Gene | Cluster | Peak | Annotation | Transcript Id | Distance to TSS |
|---|---|---|---|---|---|
| **C1orf94** | Core-like | chr1_34166188_34167441 | Promoter (<=1kb) | ENST00000373374.7 | 0 |
| **NPL** | Core-like | chr1_182751871_182753217 | Distal Intergenic | ENST00000488424.5 | -36076 |
| **GACAT3** | Core-like | chr2_16012531_16014729 | Promoter (<=1kb) | ENST00000652394.1 | 0 |
| **MIR4788** | Core-like | chr3_134405961_134407472 | Distal Intergenic | ENST00000511263.1 | -30133 |
| **LINC01991** | Core-like | chr3_187989433_187990376 | Distal Intergenic | ENST00000446091.1 | -13026 |
| **BASP1** | Core-like | chr5_17216124_17219343 | Promoter (<=1kb) | ENST00000616743.1 | 0 |
| **TJAP1** | Core-like | chr6_43488720_43490033 | Promoter (<=1kb) | ENST00000438588.6 | 0 |
| **EGFR** | Core-like | chr7_54932400_54933848 | Exon (ENST00000407916.2/ENST00000407916.2, exon 1 of 1) | ENST00000275493.7 | -85169 |
| **EGFR-AS1** | Core-like | chr7_55179528_55181017 | Exon (ENST00000442411.1/100507500, exon 2 of 2) | ENST00000442411.1 | 7917 |
| **EGFR** | Core-like | chr7_55191612_55193452 | Promoter (<=1kb) | ENST00000485503.1 | 0 |
| **EGFR** | Core-like | chr7_55212173_55214099 | Distal Intergenic | ENST00000485503.1 | 19362 |
| **LANCL2** | Core-like | chr7_55364620_55367000 | Promoter (<=1kb) | ENST00000254770.2 | 0 |
| **LANCL2** | Core-like | chr7_55469086_55470094 | Intron (ENST00000462326.5/81552, intron 4 of 4) | ENST00000466041.1 | 41030 |
| **VOPP1** | Core-like | chr7_55501620_55502991 | Intron (ENST00000462326.5/81552, intron 2 of 4) | ENST00000453256.5 | 13086 |
| **VOPP1** | Core-like | chr7_55526032_55527332 | Promoter (<=1kb) | ENST00000428097.5 | 0 |
| **HIP1** | Core-like | chr7_75687782_75689082 | Intron (ENST00000336926.11/3092, intron 1 of 30) | ENST00000485723.1 | 49853 |
| **TMEM130** | Core-like | chr7_98809689_98810889 | Distal Intergenic | ENST00000450876.5 | 56057 |
| **LOC340357** | Core-like | chr8_12774554_12775657 | Intron (ENST00000534827.5/340357, intron 3 of 4) | ENST00000530228.1 | 12799 |
| **XKR4** | Core-like | chr8_55101014_55103239 | Promoter (<=1kb) | ENST00000327381.7 | 0 |
| **NTM** | Core-like | chr11_131584471_131585636 | Promoter (2-3kb) | ENST00000416661.1 | 2460 |
| **IGSF9B** | Core-like | chr11_133964770_133966711 | Distal Intergenic | ENST00000533871.7 | -7785 |
| **CPM** | Core-like | chr12_68932270_68933761 | Promoter (<=1kb) | ENST00000551568.6 | 0 |
| **SLC41A2** | Core-like | chr12_104742146_104743389 | Intron (ENST00000303694.6/50515, intron 2 of 2) | ENST00000549713.1 | 90790 |
| **SERPINE3** | Core-like | chr13_51324598_51325436 | Distal Intergenic | ENST00000524365.5 | -10337 |
| **SALL3** | Core-like | chr18_78978450_78981071 | Promoter (<=1kb) | ENST00000616649.4 | 0 |
| **NOVA2** | Core-like | chr19_45953794_45955130 | 5' UTR | ENST00000596784.1 | 7476 |
| **SLCO4A1-AS1** | Core-like | chr20_62666387_62667815 | Promoter (<=1kb) | ENST00000451648.1 | 0 |
| **LOC101928107** | Core-like | chr21_33115571_33117877 | Intron (ENST00000657804.1/101928107, intron 1 of 2) | ENST00000655501.1 | 4208 |
| **ZPLD1** | Edge-like | chr3_102360062_102361679 | Intron (ENST00000491959.5/131368, intron 4 of 17) | ENST00000306176.5 | -73336 |
| **ZNF556** | Edge-like | chr19_2866519_2867923 | Promoter (<=1kb) | ENST00000586470.5 | 0 |
| **ZNF273** | Edge-like | chr7_64890469_64892385 | Intron (ENST00000527278.5/10793, intron 1 of 7) | ENST00000527278.5 | 7976 |
| **VPS53** | Edge-like | chr17_632975_634145 | Intron (ENST00000437048.7/55275, intron 7 of 21) | ENST00000570650.1 | -4744 |
| **UQCRHL** | Edge-like | chr1_15798858_15800806 | Distal Intergenic | ENST00000483273.2 | 8542 |

| | | | | | |
|---|---|---|---|---|---|
| **UGP2** | Edge-like | chr2_63874774_63876326 | Intron (ENST00000394417.6/7360, intron 3 of 9) | ENST00000495020.1 | 18010 |
| **UBL7** | Edge-like | chr15_74453444_74454624 | Promoter (1-2kb) | ENST00000566365.5 | -1066 |
| **UACA** | Edge-like | chr15_70799678_70800891 | Intron (ENST00000470368.1/ENST00000470368.1, intron 1 of 2) | ENST00000322954.11 | -36120 |
| **TRIM29** | Edge-like | chr11_120055298_120056866 | Distal Intergenic | ENST00000526161.5 | 60240 |
| **TRAF2** | Edge-like | chr9_136901103_136902816 | Promoter (<=1kb) | ENST00000482854.5 | 0 |
| **TMEM238L** | Edge-like | chr17_10789697_10791260 | Intron (ENST00000647474.1/101101775, intron 3 of 5) | ENST00000578763.1 | 6664 |
| **TMED7-TICAM2** | Edge-like | chr5_115685239_115686999 | Distal Intergenic | ENST00000508420.1 | -52247 |
| **TLN2** | Edge-like | chr15_62485405_62486733 | Intron (ENST00000561197.5/ENST00000561197.5, intron 1 of 2) | ENST00000561311.5 | -74632 |
| **TENT5B** | Edge-like | chr1_27026642_27028306 | Distal Intergenic | ENST00000289166.6 | -13792 |
| **TBC1D2** | Edge-like | chr9_98222455_98223900 | Intron (ENST00000465784.7/55357, intron 5 of 12) | ENST00000493589.2 | 5183 |
| **TAF1L** | Edge-like | chr9_32644344_32645517 | Exon (ENST00000413291.1/ENST00000413291.1, exon 1 of 2) | ENST00000242310.4 | -8675 |
| **STRA6** | Edge-like | chr15_74178952_74180389 | Promoter (2-3kb) | ENST00000575272.1 | 2489 |
| **SPARC** | Edge-like | chr5_151619692_151620786 | Distal Intergenic | ENST00000537849.1 | 46798 |
| **SPARC** | Edge-like | chr5_151627746_151628886 | Distal Intergenic | ENST00000537849.1 | 38698 |
| **SNX33** | Edge-like | chr15_75660019_75661747 | 3' UTR | ENST00000569152.1 | 10263 |
| **SMIM20** | Edge-like | chr4_26014916_26015807 | Distal Intergenic | ENST00000522137.1 | 86701 |
| **SMAD3** | Edge-like | chr15_67080320_67081490 | Promoter (2-3kb) | ENST00000559937.1 | 2334 |
| **SLC4A7** | Edge-like | chr3_27503157_27504140 | Distal Intergenic | ENST00000428005.1 | -18737 |
| **SLC44A3-AS1** | Edge-like | chr1_94782922_94784154 | Intron (ENST00000634870.1/101928079, intron 1 of 5) | ENST00000635408.1 | 35905 |
| **SHANK2** | Edge-like | chr11_71286646_71287944 | Distal Intergenic | ENST00000608988.5 | -34069 |
| **SEMA6D** | Edge-like | chr15_47293144_47294306 | Intron (ENST00000558014.5/80031, intron 1 of 19) | ENST00000560636.5 | -44761 |
| **SEMA5A** | Edge-like | chr5_9349954_9350823 | Intron (ENST00000382496.10/9037, intron 3 of 22) | ENST00000509486.2 | 29200 |
| **SAP30BP** | Edge-like | chr17_75700304_75700925 | Promoter (1-2kb) | ENST00000581207.5 | -1555 |
| **RTP3** | Edge-like | chr3_46495567_46496477 | Promoter (1-2kb) | ENST00000296142.4 | -1499 |
| **RNF43** | Edge-like | chr17_58401841_58403849 | Promoter (<=1kb) | ENST00000577625.5 | 0 |
| **RCSD1** | Edge-like | chr1_167662971_167664481 | Exon (ENST00000472038.1/92241, exon 2 of 2) | ENST00000461790.1 | 32705 |
| **RABGAP1** | Edge-like | chr9_123068542_123069620 | Intron (ENST00000456584.5/23637, intron 15 of 27) | ENST00000474707.1 | -3940 |
| **PTH1R** | Edge-like | chr3_46930247_46931843 | Intron (ENST00000425441.5/151903, intron 4 of 8) | ENST00000422115.2 | 27648 |
| **PRDM11** | Edge-like | chr11_45088960_45089768 | Distal Intergenic | ENST00000530656.5 | -6038 |
| **PLB1** | Edge-like | chr2_28678051_28678789 | Distal Intergenic | ENST00000436775.1 | 47465 |
| **PIK3IP1-AS1** | Edge-like | chr22_31327653_31329428 | Exon (ENST00000266269.10/23598, exon 4 of 5) | ENST00000451161.1 | -5921 |
| **PDE9A** | Edge-like | chr21_42707191_42708536 | Intron (ENST00000291539.11/5152, intron 4 of 19) | ENST00000467162.5 | 20994 |
| **PDE1A** | Edge-like | chr2_182616188_182617478 | Distal Intergenic | ENST00000495511.1 | -92996 |
| **PBOV1** | Edge-like | chr6_138293156_138294327 | Exon (ENST00000251691.5/57221, exon 20 of 34) | ENST00000527246.3 | -74665 |
| **OSTN-AS1** | Edge-like | chr3_191234825_191236288 | Promoter (<=1kb) | ENST00000430375.1 | -220 |
| **NEO1** | Edge-like | chr15_73282891_73284364 | Exon (ENST00000339362.9/4756, exon 24 of 30) | ENST00000560808.1 | -4710 |
| **NDST1** | Edge-like | chr5_150503599_150505370 | Promoter (2-3kb) | ENST00000518346.1 | -2270 |

| | | | | | |
|---|---|---|---|---|---|
| **MIR585** | Edge-like | chr5_169164107_169165056 | Intron (ENST00000519560.5/6586, intron 4 of 35) | ENST00000384887.1 | 98638 |
| **MIR585** | Edge-like | chr5_169181812_169183312 | Intron (ENST00000519560.5/6586, intron 4 of 35) | ENST00000384887.1 | 80382 |
| **MIR3914-1** | Edge-like | chr7_71307560_71308821 | Promoter (<=1kb) | ENST00000584357.2 | 0 |
| **MIR216A** | Edge-like | chr2_55992075_55993102 | Intron (ENST00000606639.1/ENST00000606639.1, intron 1 of 6) | ENST00000385063.1 | -3016 |
| **MIR205HG** | Edge-like | chr1_209411234_209412528 | Distal Intergenic | ENST00000429156.6 | -16289 |
| **METTL7A** | Edge-like | chr12_50933488_50935631 | Exon (ENST00000614647.1/ENST00000614647.1, exon 1 of 1) | ENST00000550097.1 | 8447 |
| **METTL6** | Edge-like | chr3_15443208_15444750 | Promoter (2-3kb) | ENST00000598878.1 | -2642 |
| **MEGF6** | Edge-like | chr1_3542619_3543725 | Intron (ENST00000356575.9/1953, intron 4 of 36) | ENST00000294599.8 | -11171 |
| **MAP4K3-DT** | Edge-like | chr2_39511805_39512908 | Promoter (1-2kb) | ENST00000438649.5 | -1238 |
| **MAP3K14** | Edge-like | chr17_45345106_45346967 | Distal Intergenic | ENST00000617331.2 | -28077 |
| **MAN2A1** | Edge-like | chr5_109743490_109744529 | Intron (ENST00000261483.4/4124, intron 4 of 21) | ENST00000502261.5 | -49646 |
| **LTF** | Edge-like | chr3_46469560_46470520 | 5' UTR | ENST00000431944.1 | -4655 |
| **LSM3** | Edge-like | chr3_14210388_14211961 | Distal Intergenic | ENST00000306024.4 | 31571 |
| **LRP1B** | Edge-like | chr2_140376261_140377283 | Intron (ENST00000389484.7/53353, intron 68 of 90) | ENST00000437977.5 | -18193 |
| **LOXL1-AS1** | Edge-like | chr15_73899164_73900764 | Distal Intergenic | ENST00000565689.5 | 19106 |
| **LOXL1-AS1** | Edge-like | chr15_73930166_73931374 | Promoter (1-2kb) | ENST00000564194.5 | -1918 |
| **LOC401478** | Edge-like | chr8_138080185_138081755 | Promoter (1-2kb) | ENST00000518973.1 | 1815 |
| **LOC105377480** | Edge-like | chr4_149008062_149009004 | Intron (ENST00000661928.1/ENST00000661928.1, intron 3 of 3) | ENST00000657933.1 | -65647 |
| **LOC105377448** | Edge-like | chr4_138953338_138954808 | Intron (ENST00000511951.1/105377448, intron 2 of 2) | ENST00000507038.1 | -15357 |
| **LOC101929106** | Edge-like | chr3_187207256_187208551 | Promoter (<=1kb) | ENST00000356133.3 | 0 |
| **LOC101927421** | Edge-like | chr5_124995576_124997076 | Promoter (<=1kb) | ENST00000642715.1 | 0 |
| **LMO3** | Edge-like | chr12_16585974_16587234 | Intron (ENST00000539036.5/4257, intron 4 of 4) | ENST00000616247.4 | 13626 |
| **LINC02426** | Edge-like | chr12_81887892_81889203 | Distal Intergenic | ENST00000550506.2 | -64516 |
| **LINC02408** | Edge-like | chr12_67474847_67476047 | Intron (ENST00000650195.1/100507175, intron 2 of 4) | ENST00000650195.1 | 31742 |
| **LINC02205** | Edge-like | chr15_70474235_70475668 | Distal Intergenic | ENST00000559752.1 | 30260 |
| **LINC01861** | Edge-like | chr5_153937637_153938908 | Distal Intergenic | ENST00000509568.1 | -38650 |
| **LINC01854** | Edge-like | chr2_129301074_129302207 | Distal Intergenic | ENST00000375987.3 | -27226 |
| **LINC00593** | Edge-like | chr15_69793769_69795449 | Intron (ENST00000647319.1/104472713, intron 10 of 11) | ENST00000558385.2 | -39744 |
| **LINC00111** | Edge-like | chr21_41612301_41613401 | Distal Intergenic | ENST00000413718.1 | -65780 |
| **KRT5** | Edge-like | chr12_52521813_52522993 | Promoter (1-2kb) | ENST00000552629.5 | -1419 |
| **KRT5** | Edge-like | chr12_52523583_52525119 | Distal Intergenic | ENST00000552629.5 | -3189 |
| **KDM3A** | Edge-like | chr2_86481191_86482309 | Promoter (2-3kb) | ENST00000462197.1 | -2372 |
| **KCNIP4** | Edge-like | chr4_21031428_21032748 | Intron (ENST00000382152.7/80333, intron 1 of 8) | ENST00000382149.9 | -47387 |
| **JPH2** | Edge-like | chr20_44154366_44155204 | Intron (ENST00000372980.3/57158, intron 2 of 5) | ENST00000342272.3 | 31889 |
| **INSIG2** | Edge-like | chr2_118123083_118124318 | Distal Intergenic | ENST00000479999.1 | 16424 |
| **IL34** | Edge-like | chr16_70576320_70577817 | Promoter (2-3kb) | ENST00000429149.6 | -2078 |
| **IFT122** | Edge-like | chr3_129485257_129486727 | Promoter (<=1kb) | ENST00000511425.5 | -779 |

| | | | | | |
|---|---|---|---|---|---|
| **ID1** | Edge-like | chr20_31595506_31596505 | Distal Intergenic | ENST00000376105.4 | -8778 |
| **HS6ST1** | Edge-like | chr2_128368864_128369923 | Distal Intergenic | ENST00000259241.7 | -49996 |
| **HMX1** | Edge-like | chr4_8910774_8912272 | Distal Intergenic | ENST00000400677.5 | -38935 |
| **H2AZ1-DT** | Edge-like | chr4_100074800_100077081 | Intron (ENST00000515026.1/256880, intron 5 of 5) | ENST00000515026.1 | 66729 |
| **GRIN2B** | Edge-like | chr12_13953024_13954518 | Intron (ENST00000609686.3/2904, intron 1 of 12) | ENST00000637875.1 | 25465 |
| **GPR1** | Edge-like | chr2_206229646_206231030 | Intron (ENST00000648653.1/101669764, intron 1 of 5) | ENST00000411719.1 | -11599 |
| **GALNT17** | Edge-like | chr7_71131020_71132454 | Promoter (<=1kb) | ENST00000333538.10 | 0 |
| **GALNT17** | Edge-like | chr7_71303131_71304030 | Promoter (2-3kb) | ENST00000467723.1 | 2389 |
| **GALK2** | Edge-like | chr15_49270553_49271619 | Intron (ENST00000559883.5/2585, intron 6 of 6) | ENST00000558399.5 | -12087 |
| **FTH1** | Edge-like | chr11_62012670_62014140 | Distal Intergenic | ENST00000273550.12 | -45036 |
| **FST** | Edge-like | chr5_53490429_53491264 | Distal Intergenic | ENST00000497789.2 | 6213 |
| **FOXI1** | Edge-like | chr5_170150802_170151935 | Distal Intergenic | ENST00000449804.4 | 44889 |
| **FMO6P** | Edge-like | chr1_171126276_171128284 | Intron (ENST00000669750.1/ENST00000669750.1, intron 3 of 4) | ENST00000367754.3 | -9456 |
| **FMO6P** | Edge-like | chr1_171141252_171143317 | Exon (ENST00000639860.1/ENST00000639860.1, exon 2 of 3) | ENST00000236166.4 | 3040 |
| **FIBCD1** | Edge-like | chr9_130935974_130936920 | Promoter (<=1kb) | ENST00000372337.6 | 0 |
| **EXOSC9** | Edge-like | chr4_121784450_121786061 | Distal Intergenic | ENST00000243498.9 | -15256 |
| **EXOSC7** | Edge-like | chr3_45010674_45012552 | Promoter (<=1kb) | ENST00000459856.1 | 0 |
| **ESRG** | Edge-like | chr3_54557087_54558545 | Intron (ENST00000415676.6/55799, intron 6 of 38) | ENST00000583516.1 | 81312 |
| **EPS15L1** | Edge-like | chr19_16400726_16401609 | Promoter (2-3kb) | ENST00000599790.1 | 2136 |
| **DHRSX** | Edge-like | chrX_2182060_2183035 | Distal Intergenic | ENST00000464935.6 | 83902 |
| **DEPDC1B** | Edge-like | chr5_60632449_60634234 | Intron (ENST00000265036.10/55789, intron 7 of 10) | ENST00000509006.1 | -3758 |
| **DAP** | Edge-like | chr5_10677850_10678863 | Downstream (<1kb) | ENST00000508646.1 | 82238 |
| **CYP1B1-AS1** | Edge-like | chr2_38190208_38191225 | Intron (ENST00000627992.3/285154, intron 2 of 2) | ENST00000585654.1 | 68223 |
| **CST5** | Edge-like | chr20_23919673_23921415 | Exon (ENST00000656541.1/ENST00000656541.1, exon 4 of 5) | ENST00000304710.5 | -39925 |
| **CRTC3** | Edge-like | chr15_90559883_90560786 | Intron (ENST00000268184.11/64784, intron 2 of 14) | ENST00000560927.1 | 20266 |
| **CPEB2** | Edge-like | chr4_15117791_15119036 | Intron (ENST00000502344.5/101929095, intron 2 of 5) | ENST00000509684.1 | 63658 |
| **CPEB1** | Edge-like | chr15_82553731_82555195 | Promoter (1-2kb) | ENST00000618698.4 | 1341 |
| **CORO2B** | Edge-like | chr15_68511116_68512296 | Distal Intergenic | ENST00000566799.5 | -66673 |
| **CERS6** | Edge-like | chr2_168395611_168396815 | Distal Intergenic | ENST00000305747.10 | -59047 |
| **CDC42EP1** | Edge-like | chr22_37545780_37547592 | Distal Intergenic | ENST00000430687.1 | -12888 |
| **CD300H** | Edge-like | chr17_74568771_74570105 | Promoter (1-2kb) | ENST00000651881.1 | -1428 |
| **CCL2** | Edge-like | chr17_34254191_34255499 | Promoter (<=1kb) | ENST00000580907.5 | 0 |
| **CBLN4** | Edge-like | chr20_55977385_55978889 | Distal Intergenic | ENST00000064571.3 | 26630 |
| **CACNB4** | Edge-like | chr2_152054317_152056050 | Intron (ENST00000539935.6/785, intron 2 of 13) | ENST00000635743.1 | -5528 |
| **CACNA1C-IT3** | Edge-like | chr12_2301514_2302873 | Intron (ENST00000327702.12/775, intron 3 of 47) | ENST00000542680.1 | 31738 |
| **C20orf194** | Edge-like | chr20_3314456_3315631 | Promoter (<=1kb) | ENST00000619760.1 | 459 |
| **C1orf87** | Edge-like | chr1_59994222_59995384 | Intron (ENST00000486478.5/127795, intron 3 of 5) | ENST00000488027.5 | 13461 |

| | | | | | |
|---|---|---|---|---|---|
| **BCAT1** | Edge-like | chr12_24820902_24822755 | Intron (ENST00000261192.12/586, intron 10 of 10) | ENST00000543099.1 | 7573 |
| **AUTS2** | Edge-like | chr7_70258860_70260356 | Intron (ENST00000644939.1/26053, intron 4 of 18) | ENST00000475660.2 | -56560 |
| **ATP6AP1L** | Edge-like | chr5_82383773_82385079 | Exon (ENST00000508366.5/92270, exon 7 of 8) | ENST00000514672.1 | 73737 |
| **ARNT2** | Edge-like | chr15_80450318_80451590 | 5' UTR | ENST00000525103.1 | 9078 |
| **ARHGEF26** | Edge-like | chr3_154242713_154243775 | Intron (ENST00000356448.8/26084, intron 12 of 14) | ENST00000483068.1 | 16928 |
| **ARHGAP15** | Edge-like | chr2_143229971_143231146 | Promoter (1-2kb) | ENST00000474474.5 | 1873 |
| **AOAH** | Edge-like | chr7_36784849_36786385 | Distal Intergenic | ENST00000612871.4 | -60300 |
| **ANKRD7** | Edge-like | chr7_118244936_118246153 | Intron (ENST00000433239.6/56311, intron 6 of 15) | ENST00000634332.1 | 4961 |
| **ANKRD1** | Edge-like | chr10_90929881_90931797 | Distal Intergenic | ENST00000371697.3 | -8605 |
| **AKR1B1** | Edge-like | chr7_134443215_134444800 | Exon (ENST00000467251.1/231, exon 3 of 3) | ENST00000467251.1 | 3224 |
| **ADRA1B** | Edge-like | chr5_159904905_159906013 | Intron (ENST00000641205.1/ENST00000641205.1, intron 1 of 2) | ENST00000306675.5 | -10469 |

**Note:** TSS = transcription start site
