## Supplemental Table 7 for "Identification of an epigenetically and phenotypically distinct peritumoral glioblastoma cell population linked to inferior patient outcome"

| Table S7. Transcription factor motif enrichment analysis of differential peaks in the Core-like R1 and Edge-like R2 clusters | | | | | | |
|---|---|---|---|---|---|---|
| **Gene** | **Cluster** | **Log2 fold-change** | **AUC** | **p-val** | **FDR** | **TF family** |
| **IRF2** | Edge-like | 1.09918327374757 | 0.664822302712696 | 9.13547455163731e-308 | 3.09238256213177e-307 | Tryptophan cluster factors |
| **RORA** | Edge-like | 0.891985394919715 | 0.712127524703937 | 0 | 0 | Nuclear receptors with C4 zinc fingers |
| **REL** | Edge-like | 1.2311279127584 | 0.72219052386714 | 0 | 0 | Rel homology region (RHR) factors |
| **RELA** | Edge-like | 1.27464914836359 | 0.728441161531086 | 0 | 0 | Rel homology region (RHR) factors |
| **NR1H2::RXRA** | Edge-like | 0.615960517620717 | 0.655714170077314 | 6.34195431420055e-275 | 1.94876557324706e-274 | Nuclear receptors with C4 zinc fingers |
| **MAF::NFE2** | Edge-like | 0.833473158797471 | 0.600626942986309 | 5.26509363478236e-116 | 9.74504172753577e-116 | Basic leucine zipper factors (bZIP) |
| **STAT1::STAT2** | Edge-like | 1.35977765181816 | 0.714847570448833 | 0 | 0 | STAT domain factors |
| **ESR2** | Edge-like | 0.603267187181966 | 0.662061610581792 | 1.29382520943458e-297 | 4.15731653589893e-297 | Nuclear receptors with C4 zinc fingers |
| **STAT1** | Edge-like | 1.14787219911021 | 0.743687258687516 | 0 | 0 | STAT domain factors |
| **STAT3** | Edge-like | 1.14183559515977 | 0.748226385306953 | 0 | 0 | STAT domain factors |
| **DBP** | Edge-like | 0.563035926619048 | 0.640317008189036 | 1.20926648603023e-223 | 3.20278529563655e-223 | Basic leucine zipper factors (bZIP) |
| **HSF1** | Edge-like | 0.842343744097094 | 0.709894695240362 | 0 | 0 | Heat shock factors |
| **IRF8** | Edge-like | 1.45760232716817 | 0.695773361568729 | 0 | 0 | Tryptophan cluster factors |
| **IRF9** | Edge-like | 1.55504516445466 | 0.706711041873276 | 0 | 0 | Tryptophan cluster factors |
| **MEF2B** | Edge-like | 0.6189354125188 | 0.648034838295628 | 1.10632884843139e-248 | 3.18320982298669e-248 | MADS box factors |
| **NFIA** | Edge-like | 1.15628706124727 | 0.70000995179364 | 0 | 0 | SMAD/NF-1 DNA-binding domain factors |
| **NFIX** | Edge-like | 0.842004553707615 | 0.666050526693241 | 0 | 0 | SMAD/NF-1 DNA-binding domain factors |
| **ESRRB** | Edge-like | 0.961854161419103 | 0.730991010124132 | 0 | 0 | Nuclear receptors with C4 zinc fingers |
| **NR2F1** | Edge-like | 1.26981348458632 | 0.779266241806865 | 0 | 0 | Nuclear receptors with C4 zinc fingers |
| **RARA** | Edge-like | 0.773431329336582 | 0.697432471930145 | 0 | 0 | Nuclear receptors with C4 zinc fingers |
| **HIC2** | Edge-like | 0.66490715978853 | 0.675154950254174 | 0 | 0 | C2H2 zinc finger factors |
| **HSF2** | Edge-like | 0.947242922540243 | 0.727948715187951 | 0 | 0 | Heat shock factors |
| **HSF4** | Edge-like | 0.754419125794218 | 0.69123187341628 | 0 | 0 | Heat shock factors |
| **IRF7** | Edge-like | 1.28514637750277 | 0.695418277729171 | 0 | 0 | Tryptophan cluster factors |
| **MEF2D** | Edge-like | 0.506042110926183 | 0.624622159721205 | 7.48883442959435e-177 | 1.6810043240898e-176 | MADS box factors |
| **NFKB1** | Edge-like | 0.628163607490227 | 0.653551145928558 | 2.12655943726889e-267 | 6.41005773233907e-267 | Rel homology region (RHR) factors |
| **NFKB2** | Edge-like | 0.625545304300096 | 0.652131597747994 | 1.61941821949911e-262 | 4.79014828478008e-262 | Rel homology region (RHR) factors |
| **RUNX2** | Edge-like | 1.38622891109474 | 0.670632361036824 | 0 | 0 | Runt domain factors |
| **TEAD3** | Edge-like | 3.289187810675 | 0.802924980113424 | 0 | 0 | TEA domain factors |
| **TFAP2A(var.2)** | Edge-like | 0.579795772303313 | 0.663529196613273 | 5.45551743394262e-303 | 1.80803274119669e-302 | Basic helix-span-helix factors (bHSH) |
| **TFAP2B** | Edge-like | 0.898705899003838 | 0.710508193498892 | 0 | 0 | Basic helix-span-helix factors (bHSH) |
| **TFAP2C** | Edge-like | 0.81620267494316 | 0.696264413673447 | 0 | 0 | Basic helix-span-helix factors (bHSH) |
| **CEBPG** | Edge-like | 0.892443206896619 | 0.684159395579901 | 0 | 0 | Basic leucine zipper factors (bZIP) |
| **TEF** | Edge-like | 0.537885237745425 | 0.633756729518534 | 2.06013770660245e-203 | 5.09401237609122e-203 | Basic leucine zipper factors (bZIP) |
| **RXRB** | Edge-like | 0.698544693269215 | 0.672988430502879 | 0 | 0 | Nuclear receptors with C4 zinc fingers |
| **RXRG** | Edge-like | 0.7124403254806 | 0.675765486589941 | 0 | 0 | Nuclear receptors with C4 zinc fingers |
| **TP63** | Edge-like | 1.28427804876697 | 0.697348017105451 | 0 | 0 | p53 domain factors |
| **TP53** | Edge-like | 1.4753404187581 | 0.692577526960735 | 0 | 0 | p53 domain factors |

| | | | | | | |
|---|---|---|---|---|---|---|
| **TP73** | Edge-like | 1.30596035090243 | 0.676308619169887 | 0 | 0 | p53 domain factors |
| **GATA2** | Edge-like | 0.672816966932877 | 0.643910520953782 | 3.89652386216812e-235 | 1.08179807225983e-234 | Other C4 zinc finger-type factors |
| **NFIC** | Edge-like | 1.37814729119452 | 0.716950715726621 | 0 | 0 | SMAD/NF-1 DNA-binding domain factors |
| **NR2F2** | Edge-like | 1.19205096576988 | 0.7697030937937 | 0 | 0 | Nuclear receptors with C4 zinc fingers |
| **TEAD2** | Edge-like | 3.14248119051757 | 0.807921096725817 | 0 | 0 | TEA domain factors |
| **PPARA::RXRA** | Edge-like | 0.726040255779492 | 0.683761250453344 | 0 | 0 | Nuclear receptors with C4 zinc fingers |
| **RORB** | Edge-like | 0.819184023184632 | 0.688541733751882 | 0 | 0 | Nuclear receptors with C4 zinc fingers |
| **RORC** | Edge-like | 0.913235592368726 | 0.70391818219382 | 0 | 0 | Nuclear receptors with C4 zinc fingers |
| **IRF3** | Edge-like | 1.06776732731875 | 0.679203571790664 | 0 | 0 | Tryptophan cluster factors |
| **IRF4** | Edge-like | 1.56029380177193 | 0.710091996653881 | 0 | 0 | Tryptophan cluster factors |
| **IRF5** | Edge-like | 0.650632883808358 | 0.640358721934598 | 8.9286887804591e-224 | 2.3747310916095e-223 | Tryptophan cluster factors |
| **HNF4A(var.2)** | Edge-like | 0.610940247797697 | 0.657166739694073 | 4.89270172263457e-280 | 1.53320801506321e-279 | Nuclear receptors with C4 zinc fingers |
| **KLF3** | Edge-like | 0.567556986192888 | 0.674031607691782 | 0 | 0 | C2H2 zinc finger factors |
| **KLF6** | Edge-like | 0.601234130339529 | 0.675873743532685 | 0 | 0 | C2H2 zinc finger factors |
| **NR1I3** | Edge-like | 0.5198560182773 | 0.629119859447959 | 1.07310829880387e-189 | 2.52519536484332e-189 | Nuclear receptors with C4 zinc fingers |
| **NR2C2(var.2)** | Edge-like | 1.0000968416852 | 0.729594650329875 | 0 | 0 | Nuclear receptors with C4 zinc fingers |
| **NR2F1(var.2)** | Edge-like | 0.765615371778263 | 0.687172885150124 | 0 | 0 | Nuclear receptors with C4 zinc fingers |
| **NR6A1** | Edge-like | 0.685651658755789 | 0.679363435567833 | 0 | 0 | Nuclear receptors with C4 zinc fingers |
| **PPARD** | Edge-like | 0.667981378856978 | 0.668434387533678 | 0 | 0 | Nuclear receptors with C4 zinc fingers |
| **RARB(var.3)** | Edge-like | 0.63720435835319 | 0.666061046855844 | 0 | 0 | Nuclear receptors with C4 zinc fingers |
| **RARG(var.3)** | Edge-like | 0.66453158917393 | 0.671734016187281 | 0 | 0 | Nuclear receptors with C4 zinc fingers |
| **RXRB(var.2)** | Edge-like | 0.607097994794605 | 0.657243029218181 | 2.62856699419012e-280 | 8.27802441453903e-280 | Nuclear receptors with C4 zinc fingers |
| **RXRG(var.2)** | Edge-like | 0.611455468884692 | 0.66093425207908 | 1.61425071650515e-293 | 5.16071062397859e-293 | Nuclear receptors with C4 zinc fingers |
| **THRB** | Edge-like | 0.657716043810373 | 0.664783190656046 | 1.27568350284571e-307 | 4.27252728730865e-307 | Nuclear receptors with C4 zinc fingers |
| **ZBTB26** | Edge-like | 1.01225816492445 | 0.744966028813931 | 0 | 0 | C2H2 zinc finger factors |
| **ZBTB6** | Edge-like | 0.507287489741898 | 0.645227902812056 | 2.01889438996005e-239 | 5.65469092409163e-239 | C2H2 zinc finger factors |
| **ZNF135** | Edge-like | 1.00496983997273 | 0.73306875224454 | 0 | 0 | C2H2 zinc finger factors |
| **ZNF136** | Edge-like | 0.642997288060309 | 0.663118663444895 | 1.7590966155999e-301 | 5.73973277151927e-301 | C2H2 zinc finger factors |
| **ZNF460** | Edge-like | 0.532435964300076 | 0.695717485296088 | 0 | 0 | C2H2 zinc finger factors |
| **ZNF528** | Edge-like | 0.543235524966521 | 0.650242097826892 | 4.33444818302987e-256 | 1.26438050684696e-255 | C2H2 zinc finger factors |
| **CEBPG(var.2)** | Edge-like | 1.15506697675446 | 0.715596743448644 | 0 | 0 | Basic leucine zipper factors (bZIP) |
| **EBF3** | Edge-like | 0.748942942317381 | 0.699718489254484 | 0 | 0 | Rel homology region (RHR) factors |
| **ZBTB12** | Edge-like | 0.713812835643133 | 0.683558959133911 | 0 | 0 | C2H2 zinc finger factors |
| **CEBPA** | Edge-like | 1.14916607569117 | 0.720243553305064 | 0 | 0 | Basic leucine zipper factors (bZIP) |
| **CEBPD** | Edge-like | 0.885365140181014 | 0.675971020014019 | 0 | 0 | Basic leucine zipper factors (bZIP) |
| **EBF1** | Edge-like | 0.772804475160697 | 0.702321652457242 | 0 | 0 | Rel homology region (RHR) factors |
| **ESRRA** | Edge-like | 1.01165110850617 | 0.743481811986402 | 0 | 0 | Nuclear receptors with C4 zinc fingers |
| **GATA4** | Edge-like | 0.570115960578189 | 0.624363191609465 | 3.98200536773938e-176 | 8.90674698861847e-176 | Other C4 zinc finger-type factors |
| **GATA6** | Edge-like | 0.604162303722128 | 0.634653625080591 | 4.02697881193362e-206 | 1.01153872537856e-205 | Other C4 zinc finger-type factors |
| **HLF** | Edge-like | 0.79631654750051 | 0.674843948364187 | 0 | 0 | Basic leucine zipper factors (bZIP) |
| **HNF4A** | Edge-like | 0.755261392725437 | 0.687678099781921 | 0 | 0 | Nuclear receptors with C4 zinc fingers |
| **HNF4G** | Edge-like | 0.732741045893844 | 0.682043822234308 | 0 | 0 | Nuclear receptors with C4 zinc fingers |
| **MAFK** | Edge-like | 1.31397702328113 | 0.636472512496212 | 1.13731241006714e-211 | 2.92649900639228e-211 | Basic leucine zipper factors (bZIP) |

| | | | | | | |
|---|---|---|---|---|---|---|
| **MEF2A** | Edge-like | 0.501162975630266 | 0.627384226077534 | 1.09848640817204e-184 | 2.55640403078272e-184 | MADS box factors |
| **NFE2L1** | Edge-like | 0.987788342162399 | 0.602557976536048 | 2.00839826673592e-120 | 3.77245134375027e-120 | Basic leucine zipper factors (bZIP) |
| **NFIL3** | Edge-like | 0.667880296933832 | 0.650385724396849 | 1.41635062257534e-256 | 4.15069418560272e-256 | Basic leucine zipper factors (bZIP) |
| **RUNX3** | Edge-like | 1.40168993558409 | 0.712771262592825 | 0 | 0 | Runt domain factors |
| **TEAD1** | Edge-like | 2.93555946331386 | 0.809891068886278 | 0 | 0 | TEA domain factors |
| **TEAD4** | Edge-like | 3.19922131367136 | 0.804258258958052 | 0 | 0 | TEA domain factors |
| **TFAP2A** | Edge-like | 0.842986104726836 | 0.703075395090143 | 0 | 0 | Basic helix-span-helix factors (bHSH) |
| **TFAP2C(var.2)** | Edge-like | 0.640497912003452 | 0.685190138030004 | 0 | 0 | Basic helix-span-helix factors (bHSH) |
| **TAL1::TCF3** | Core-like | 1.86325485382186 | 0.792868124938652 | 0 | 0 | Basic helix-loop-helix factors (bHLH) |
| **ELK4** | Core-like | 0.681471940658312 | 0.657898828270026 | 1.24399887507092e-282 | 3.93725643959945e-282 | Tryptophan cluster factors |
| **LBX1** | Core-like | 1.12608655123602 | 0.70785337009373 | 0 | 0 | Homeo domain factors(Homeobox) |
| **POU6F1** | Core-like | 0.718422179359969 | 0.663862546343918 | 3.22996568018039e-304 | 1.07608856608115e-303 | Homeo domain factors(Homeobox) |
| **SHOX** | Core-like | 0.954970330752711 | 0.69678848721436 | 0 | 0 | Homeo domain factors(Homeobox) |
| **ALX3** | Core-like | 1.0730103301761 | 0.71327923234647 | 0 | 0 | Homeo domain factors(Homeobox) |
| **BHLHE41** | Core-like | 0.577129614441183 | 0.64613848726039 | 2.0898516867004e-242 | 5.87944941191712e-242 | Basic helix-loop-helix factors (bHLH) |
| **EN1** | Core-like | 0.575519678829813 | 0.629983266604015 | 3.25912937411785e-192 | 7.72669997684118e-192 | Homeo domain factors(Homeobox) |
| **FLI1** | Core-like | 0.596848091141509 | 0.635981427036508 | 3.64263348488408e-210 | 9.29752820940171e-210 | Tryptophan cluster factors |
| **HEY2** | Core-like | 0.509403744019978 | 0.636022053409525 | 2.73571882471609e-210 | 7.01097172487969e-210 | Basic helix-loop-helix factors (bHLH) |
| **HINFP** | Core-like | 0.537232042085553 | 0.644434567810707 | 7.77273545148412e-237 | 2.16746323382795e-236 | C2H2 zinc finger factors |
| **ISX** | Core-like | 0.954970330752711 | 0.69678848721436 | 0 | 0 | Homeo domain factors(Homeobox) |
| **JDP2(var.2)** | Core-like | 0.643922143861641 | 0.622679165073466 | 1.91599192373401e-171 | 4.2111905823737e-171 | Basic leucine zipper factors (bZIP) |
| **LHX6** | Core-like | 1.11552943411475 | 0.7172952260356 | 0 | 0 | Homeo domain factors(Homeobox) |
| **MEOX1** | Core-like | 1.09888775623782 | 0.729448595516932 | 0 | 0 | Homeo domain factors(Homeobox) |
| **MIXL1** | Core-like | 1.09279516175051 | 0.717365084718396 | 0 | 0 | Homeo domain factors(Homeobox) |
| **MLXIPL** | Core-like | 0.547105383082262 | 0.640196990278209 | 2.89285914735143e-223 | 7.59825659864504e-223 | Basic helix-loop-helix factors (bHLH) |
| **MSC** | Core-like | 1.8129642426768 | 0.829094901913193 | 0 | 0 | Basic helix-loop-helix factors (bHLH) |
| **MYF6** | Core-like | 1.30006204739664 | 0.76624689683555 | 0 | 0 | Basic helix-loop-helix factors (bHLH) |
| **NEUROG2** | Core-like | 1.05885022949847 | 0.729982001782944 | 0 | 0 | Basic helix-loop-helix factors (bHLH) |
| **NHLH1** | Core-like | 2.59453725666786 | 0.892266283580649 | 0 | 0 | Basic helix-loop-helix factors (bHLH) |
| **NKX6-1** | Core-like | 1.22778315511328 | 0.741214586861207 | 0 | 0 | Homeo domain factors(Homeobox) |
| **NKX6-2** | Core-like | 1.22778315511328 | 0.741214586861207 | 0 | 0 | Homeo domain factors(Homeobox) |
| **OLIG2** | Core-like | 0.743173928050698 | 0.679347545252476 | 0 | 0 | Basic helix-loop-helix factors (bHLH) |
| **PAX4** | Core-like | 1.09241750166836 | 0.707817306682806 | 0 | 0 | Paired box factors |
| **SPDEF** | Core-like | 0.5901264996419 | 0.649152063553576 | 2.0345270736766e-252 | 5.90759466806096e-252 | Tryptophan cluster factors |
| **TFAP4** | Core-like | 1.11512948743797 | 0.729526406028404 | 0 | 0 | Basic helix-loop-helix factors (bHLH) |
| **ZBTB18** | Core-like | 1.00336061447988 | 0.656944975733781 | 2.97280887512454e-279 | 9.26989171405829e-279 | C2H2 zinc finger factors |
| **LBX2** | Core-like | 0.818463421714491 | 0.670597731890301 | 0 | 0 | Homeo domain factors(Homeobox) |
| **MEOX2** | Core-like | 1.10632785438887 | 0.734182581964908 | 0 | 0 | Homeo domain factors(Homeobox) |
| **NOTO** | Core-like | 1.06282884128276 | 0.706121679282656 | 0 | 0 | Homeo domain factors(Homeobox) |
| **PDX1** | Core-like | 0.798686214730164 | 0.67943685656314 | 0 | 0 | Homeo domain factors(Homeobox) |
| **PHOX2A** | Core-like | 0.636471278239845 | 0.654883344071638 | 5.07236942981643e-272 | 1.54365858128548e-271 | Homeo domain factors(Homeobox) |
| **PROP1** | Core-like | 0.668444949063637 | 0.663317525871762 | 3.27391141826981e-302 | 1.07936767071083e-301 | Homeo domain factors(Homeobox) |
| **PRRX1** | Core-like | 0.954970330752711 | 0.69678848721436 | 0 | 0 | Homeo domain factors(Homeobox) |

| | | | | | | |
|---|---|---|---|---|---|---|
| **RAX2** | Core-like | 0.954970330752711 | 0.69678848721436 | 0 | 0 | Homeo domain factors(Homeobox) |
| **RAX** | Core-like | 0.985760971092778 | 0.704862595269012 | 0 | 0 | Homeo domain factors(Homeobox) |
| **UNCX** | Core-like | 1.16715056022078 | 0.72729407688962 | 0 | 0 | Homeo domain factors(Homeobox) |
| **VAX1** | Core-like | 1.00729889634577 | 0.72606253742346 | 0 | 0 | Homeo domain factors(Homeobox) |
| **VSX1** | Core-like | 0.905052669683634 | 0.683951967592616 | 0 | 0 | Homeo domain factors(Homeobox) |
| **VSX2** | Core-like | 0.905052669683634 | 0.683951967592616 | 0 | 0 | Homeo domain factors(Homeobox) |
| **EGR3** | Core-like | 0.504186215859092 | 0.655764255924376 | 4.23372532718683e-275 | 1.30729177176062e-274 | C2H2 zinc finger factors |
| **CUX2** | Core-like | 0.64068140804847 | 0.665578019960771 | 0 | 0 | Homeo domain factors(Homeobox) |
| **ERF** | Core-like | 0.657379471004141 | 0.646282020436386 | 7.0452824375131e-243 | 1.99984923002054e-242 | Tryptophan cluster factors |
| **ETS1** | Core-like | 0.631920719478707 | 0.641977492775877 | 6.43330764699312e-229 | 1.75529471575287e-228 | Tryptophan cluster factors |
| **ETV2** | Core-like | 0.626492199591296 | 0.639365510514806 | 1.19341709988054e-220 | 3.12162406704291e-220 | Tryptophan cluster factors |
| **PAX3** | Core-like | 0.770687868717408 | 0.6884854772324 | 0 | 0 | Paired box factors |
| **POU2F1** | Core-like | 1.27544126187243 | 0.723113429647323 | 0 | 0 | Homeo domain factors(Homeobox) |
| **POU3F1** | Core-like | 1.23067624955415 | 0.717260690283038 | 0 | 0 | Homeo domain factors(Homeobox) |
| **POU3F2** | Core-like | 1.33332432725126 | 0.717797818965493 | 0 | 0 | Homeo domain factors(Homeobox) |
| **POU3F3** | Core-like | 1.30356979042826 | 0.73355432078955 | 0 | 0 | Homeo domain factors(Homeobox) |
| **POU3F4** | Core-like | 1.05348353539705 | 0.688444050339719 | 0 | 0 | Homeo domain factors(Homeobox) |
| **POU4F3** | Core-like | 0.50195447905771 | 0.615930837662577 | 2.55117469398615e-153 | 5.24316097822478e-153 | Homeo domain factors(Homeobox) |
| **POU5F1B** | Core-like | 1.09528443802719 | 0.694337949756797 | 0 | 0 | Homeo domain factors(Homeobox) |
| **POU6F2** | Core-like | 0.993401975289138 | 0.704342024001634 | 0 | 0 | Homeo domain factors(Homeobox) |
| **RFX2** | Core-like | 2.06715280992955 | 0.791642062361202 | 0 | 0 | Fork head/winged helix factors |
| **TGIF1** | Core-like | 0.509652207607499 | 0.626186715368148 | 2.87172523647826e-181 | 6.56246236350448e-181 | Homeo domain factors(Homeobox) |
| **RFX5** | Core-like | 1.88106497836796 | 0.775290033945102 | 0 | 0 | Fork head/winged helix factors |
| **BHLHE40** | Core-like | 0.691057254683596 | 0.673542423466284 | 0 | 0 | Basic helix-loop-helix factors (bHLH) |
| **BHLHE23** | Core-like | 1.03635882213501 | 0.737665102678055 | 0 | 0 | Basic helix-loop-helix factors (bHLH) |
| **FIGLA** | Core-like | 1.1726573211007 | 0.701730095108541 | 0 | 0 | Basic helix-loop-helix factors (bHLH) |
| **HEY1** | Core-like | 0.572901970488485 | 0.653787059073935 | 3.25094707138669e-268 | 9.84616983821903e-268 | Basic helix-loop-helix factors (bHLH) |
| **MNT** | Core-like | 0.533653710753998 | 0.638519027678675 | 5.29054247680802e-218 | 1.37815365753888e-217 | Basic helix-loop-helix factors (bHLH) |
| **OLIG1** | Core-like | 0.999877622879463 | 0.729461517238531 | 0 | 0 | Basic helix-loop-helix factors (bHLH) |
| **OLIG3** | Core-like | 0.79054014206131 | 0.68893789092127 | 0 | 0 | Basic helix-loop-helix factors (bHLH) |
| **ATF7** | Core-like | 0.835676515701805 | 0.664812362926851 | 9.9445952052904e-308 | 3.34836636433448e-307 | Basic leucine zipper factors (bZIP) |
| **MTF1** | Core-like | 0.535209269019501 | 0.643792244173227 | 9.40882908177785e-235 | 2.58947339511538e-234 | C2H2 zinc finger factors |
| **BARX1** | Core-like | 0.697876400505633 | 0.662671386423878 | 7.66380681468493e-300 | 2.48778959676695e-299 | Homeo domain factors(Homeobox) |
| **BSX** | Core-like | 0.500612695206394 | 0.618286907128632 | 1.56705060734336e-159 | 3.31753523226872e-159 | Homeo domain factors(Homeobox) |
| **EMX2** | Core-like | 0.987238065046911 | 0.703707085158068 | 0 | 0 | Homeo domain factors(Homeobox) |
| **EVX1** | Core-like | 0.952455823807564 | 0.708384601614651 | 0 | 0 | Homeo domain factors(Homeobox) |
| **EVX2** | Core-like | 1.00725563201078 | 0.714571097640411 | 0 | 0 | Homeo domain factors(Homeobox) |
| **GBX1** | Core-like | 0.955918115270255 | 0.699304413786587 | 0 | 0 | Homeo domain factors(Homeobox) |
| **GBX2** | Core-like | 0.987155444342814 | 0.69942291071005 | 0 | 0 | Homeo domain factors(Homeobox) |
| **GSX1** | Core-like | 1.12438208346125 | 0.71887378410569 | 0 | 0 | Homeo domain factors(Homeobox) |
| **HESX1** | Core-like | 1.00864074099205 | 0.702329737705862 | 0 | 0 | Homeo domain factors(Homeobox) |
| **HOXB3** | Core-like | 1.23957213165073 | 0.740415041161454 | 0 | 0 | Homeo domain factors(Homeobox) |
| **NEUROD1** | Core-like | 1.71344942945453 | 0.780103918926677 | 0 | 0 | Basic helix-loop-helix factors (bHLH) |

| | | | | | | |
|---|---|---|---|---|---|---|
| **POU5F1** | Core-like | 0.884106517793516 | 0.679923545835681 | 0 | 0 | Homeo domain factors(Homeobox) |
| **SOX10** | Core-like | 0.984748396447851 | 0.682885595346408 | 0 | 0 | High-mobility group (HMG) domain factors |
| **SOX13** | Core-like | 0.875063044280334 | 0.673999793706082 | 0 | 0 | High-mobility group (HMG) domain factors |
| **ZBTB7A** | Core-like | 0.534175008011328 | 0.642173746842834 | 1.51702732464695e-229 | 4.15704890260399e-229 | C2H2 zinc finger factors |
| **ZEB1** | Core-like | 0.71447369841536 | 0.618192452479118 | 2.7957979503212e-159 | 5.89913367517774e-159 | Homeo domain factors(Homeobox) |
| **FOS::JUN(var.2)** | Core-like | 0.549465178204179 | 0.603677976928836 | 5.03204203560094e-123 | 9.56541323884503e-123 | Basic leucine zipper factors (bZIP) |
| **FOSB::JUN** | Core-like | 0.670331539894945 | 0.626085843219318 | 5.55305927816232e-181 | 1.26441961261754e-180 | Basic leucine zipper factors (bZIP) |
| **FOSL1::JUN(var.2)** | Core-like | 0.598703437606039 | 0.618981031059456 | 2.19429370361113e-161 | 4.7405730866411e-161 | Basic leucine zipper factors (bZIP) |
| **FOSL2::JUN(var.2)** | Core-like | 0.595271569640916 | 0.606534164384819 | 8.73091695953733e-130 | 1.70051398011912e-129 | Basic leucine zipper factors (bZIP) |
| **JUN::JUNB(var.2)** | Core-like | 0.64884076388976 | 0.630286570163221 | 4.21452971948102e-193 | 1.00293132046296e-192 | Basic leucine zipper factors (bZIP) |
| **FOSB::JUNB(var.2)** | Core-like | 0.613054762807583 | 0.620293643154724 | 6.39954600744221e-165 | 1.39206619337145e-164 | Basic leucine zipper factors (bZIP) |
| **FOSL2::JUNB(var.2)** | Core-like | 0.729901240479147 | 0.641352593783114 | 6.3172343397427e-227 | 1.70889287908424e-226 | Basic leucine zipper factors (bZIP) |
| **FOSL2::JUND(var.2)** | Core-like | 0.703934458489897 | 0.625455507363827 | 3.38037512129712e-179 | 7.64206232778956e-179 | Basic leucine zipper factors (bZIP) |
| **ARGFX** | Core-like | 1.21663939220782 | 0.733043715992007 | 0 | 0 | Homeo domain factors(Homeobox) |
| **ARNT2** | Core-like | 0.553937127943682 | 0.641032352561989 | 6.57677997316798e-226 | 1.76402615382006e-225 | Basic helix-loop-helix factors (bHLH) |
| **ATOH7** | Core-like | 1.23983909316202 | 0.73375583160231 | 0 | 0 | Basic helix-loop-helix factors (bHLH) |
| **DRGX** | Core-like | 1.00729889634577 | 0.72606253742346 | 0 | 0 | Homeo domain factors(Homeobox) |
| **ETS2** | Core-like | 0.552857127410905 | 0.632317141662348 | 4.20874684902077e-199 | 1.03261114551556e-198 | Tryptophan cluster factors |
| **FERD3L** | Core-like | 1.31673710019013 | 0.781294771980846 | 0 | 0 | Basic helix-loop-helix factors (bHLH) |
| **HOXA1** | Core-like | 0.626827579292586 | 0.652207426973294 | 8.90706131401068e-263 | 2.64702808064261e-262 | Homeo domain factors(Homeobox) |
| **HOXA6** | Core-like | 1.12541828568963 | 0.742193822541841 | 0 | 0 | Homeo domain factors(Homeobox) |
| **HOXB4** | Core-like | 0.527683807777238 | 0.630617344888941 | 4.50381794827e-194 | 1.0881361684179e-193 | Homeo domain factors(Homeobox) |
| **HOXB6** | Core-like | 0.959045477592061 | 0.710023792378394 | 0 | 0 | Homeo domain factors(Homeobox) |
| **HOXB7** | Core-like | 1.1563233219815 | 0.746765997281462 | 0 | 0 | Homeo domain factors(Homeobox) |
| **HOXC4** | Core-like | 0.511797301715545 | 0.63042789523914 | 1.62223385824167e-193 | 3.90446400101512e-193 | Homeo domain factors(Homeobox) |
| **HOXC8** | Core-like | 0.84520743319308 | 0.69962520870048 | 0 | 0 | Homeo domain factors(Homeobox) |
| **LHX5** | Core-like | 1.22778315511328 | 0.741214586861207 | 0 | 0 | Homeo domain factors(Homeobox) |
| **MAZ** | Core-like | 0.628660034644659 | 0.721637678315255 | 0 | 0 | C2H2 zinc finger factors |
| **NFIC(var.2)** | Core-like | 0.792238274104338 | 0.648139706371788 | 4.94825629026732e-249 | 1.43024942088549e-248 | SMAD/NF-1 DNA-binding domain factors |
| **NFIX(var.2)** | Core-like | 0.785488304015855 | 0.641451531342753 | 3.0599330680748e-227 | 8.31303704760236e-227 | SMAD/NF-1 DNA-binding domain factors |
| **NHLH2** | Core-like | 1.74493382438688 | 0.84790197738097 | 0 | 0 | Basic helix-loop-helix factors (bHLH) |
| **NKX6-3** | Core-like | 0.518419194258199 | 0.630319238036564 | 3.38021430106014e-193 | 8.10483201731465e-193 | Homeo domain factors(Homeobox) |
| **POU6F1(var.2)** | Core-like | 0.634246381565415 | 0.646777682200177 | 1.63545118558624e-244 | 4.66324594809049e-244 | Homeo domain factors(Homeobox) |
| **RFX7** | Core-like | 0.787490776547953 | 0.692530736586296 | 0 | 0 | Fork head/winged helix factors |
| **SNAI1** | Core-like | 1.4510569367871 | 0.756958600578485 | 0 | 0 | C2H2 zinc finger factors |
| **SNAI3** | Core-like | 1.3707764516487 | 0.751047856893448 | 0 | 0 | C2H2 zinc finger factors |
| **SOHLH2** | Core-like | 0.786403752036385 | 0.689947379574258 | 0 | 0 | Basic helix-loop-helix factors (bHLH) |
| **SOX14** | Core-like | 0.524302081046041 | 0.620495047231527 | 1.82096590919551e-165 | 3.97472903627848e-165 | High-mobility group (HMG) domain factors |
| **TCF21(var.2)** | Core-like | 1.12232828161906 | 0.750084671632103 | 0 | 0 | Basic helix-loop-helix factors (bHLH) |
| **TLX2** | Core-like | 1.18970290634508 | 0.717976201430921 | 0 | 0 | Homeo domain factors(Homeobox) |
| **VEZF1** | Core-like | 0.727872767019815 | 0.699476091899818 | 0 | 0 | C2H2 zinc finger factors |
| **ASCL1** | Core-like | 3.00083014750166 | 0.900503344084179 | 0 | 0 | Basic helix-loop-helix factors (bHLH) |
| **ATF3** | Core-like | 0.567311108471243 | 0.610355298245409 | 4.07375033864187e-139 | 8.05838738862594e-139 | Basic leucine zipper factors (bZIP) |

| | | | | | | |
|---|---|---|---|---|---|---|
| **EMX1** | Core-like | 0.798686214730164 | 0.67943685656314 | 0 | 0 | Homeo domain factors(Homeobox) |
| **GSX2** | Core-like | 0.798686214730164 | 0.67943685656314 | 0 | 0 | Homeo domain factors(Homeobox) |
| **HES1** | Core-like | 0.525776355049057 | 0.646858621409341 | 8.83581142380231e-245 | 2.53080028564111e-244 | Basic helix-loop-helix factors (bHLH) |
| **HES2** | Core-like | 0.670133950451133 | 0.675814251579555 | 0 | 0 | Basic helix-loop-helix factors (bHLH) |
| **HOXA2** | Core-like | 0.824462851571846 | 0.68510144712208 | 0 | 0 | Homeo domain factors(Homeobox) |
| **HOXA5** | Core-like | 0.626827579292586 | 0.652207426973294 | 8.90706131401068e-263 | 2.64702808064261e-262 | Homeo domain factors(Homeobox) |
| **HOXB2** | Core-like | 0.798686214730164 | 0.67943685656314 | 0 | 0 | Homeo domain factors(Homeobox) |
| **HOXB5** | Core-like | 0.626827579292586 | 0.652207426973294 | 8.90706131401068e-263 | 2.64702808064261e-262 | Homeo domain factors(Homeobox) |
| **HOXD3** | Core-like | 0.798686214730164 | 0.67943685656314 | 0 | 0 | Homeo domain factors(Homeobox) |
| **JUNB(var.2)** | Core-like | 0.62762398881978 | 0.631785850102452 | 1.59635901512649e-197 | 3.88652021759642e-197 | Basic leucine zipper factors (bZIP) |
| **LHX9** | Core-like | 1.09241750166836 | 0.707817306682806 | 0 | 0 | Homeo domain factors(Homeobox) |
| **LMX1A** | Core-like | 1.02028166901839 | 0.725432521775836 | 0 | 0 | Homeo domain factors(Homeobox) |
| **LMX1B** | Core-like | 1.12903002491495 | 0.733245486973657 | 0 | 0 | Homeo domain factors(Homeobox) |
| **NEUROG1** | Core-like | 1.05230499582617 | 0.739712818640015 | 0 | 0 | Basic helix-loop-helix factors (bHLH) |
| **PRRX2** | Core-like | 1.24231786531049 | 0.718349904023696 | 0 | 0 | Homeo domain factors(Homeobox) |
| **SOX4** | Core-like | 1.22711953559055 | 0.70853621336827 | 0 | 0 | High-mobility group (HMG) domain factors |
| **TCFL5** | Core-like | 0.585751791956679 | 0.65180453876772 | 2.12650151539159e-261 | 6.26081608950175e-261 | Basic helix-loop-helix factors (bHLH) |
| **ZNF740** | Core-like | 0.697080906312539 | 0.731387514185375 | 0 | 0 | C2H2 zinc finger factors |
| **ASCL1(var.2)** | Core-like | 1.92289289020845 | 0.841086773223039 | 0 | 0 | Basic helix-loop-helix factors (bHLH) |
| **BHLHE22(var.2)** | Core-like | 2.70046151636989 | 0.885014876257071 | 0 | 0 | Basic helix-loop-helix factors (bHLH) |
| **HAND2** | Core-like | 1.43692862196955 | 0.738928469471909 | 0 | 0 | Basic helix-loop-helix factors (bHLH) |
| **MYF5** | Core-like | 2.32967388468704 | 0.841946624737581 | 0 | 0 | Basic helix-loop-helix factors (bHLH) |
| **NEUROG2(var.2)** | Core-like | 1.64623476947071 | 0.770230676279147 | 0 | 0 | Basic helix-loop-helix factors (bHLH) |
| **TCF12(var.2)** | Core-like | 1.40655007743147 | 0.749913880761502 | 0 | 0 | Basic helix-loop-helix factors (bHLH) |
| **ZNF148** | Core-like | 0.748188740749305 | 0.773845035910178 | 0 | 0 | C2H2 zinc finger factors |
| **CTCFL** | Core-like | 0.630212427659598 | 0.705046474636141 | 0 | 0 | C2H2 zinc finger factors |
| **EGR1** | Core-like | 0.548672625524904 | 0.683272526526653 | 0 | 0 | C2H2 zinc finger factors |
| **EHF** | Core-like | 0.506872956294055 | 0.609157954310954 | 3.70782663453804e-136 | 7.26642185654049e-136 | Tryptophan cluster factors |
| **ETV1** | Core-like | 0.563545209733655 | 0.613239790742558 | 2.23406654712599e-146 | 4.48940991851032e-146 | Tryptophan cluster factors |
| **GABPA** | Core-like | 0.568936545959167 | 0.618717586708588 | 1.11210547202014e-160 | 2.38631445352118e-160 | Tryptophan cluster factors |
| **MYOD1** | Core-like | 2.03449825409248 | 0.857567725331243 | 0 | 0 | Basic helix-loop-helix factors (bHLH) |
| **MYOG** | Core-like | 3.00405859033508 | 0.892607791940883 | 0 | 0 | Basic helix-loop-helix factors (bHLH) |
| **ONECUT1** | Core-like | 0.513835540358004 | 0.637579077500617 | 4.40146989787174e-215 | 1.14185673989869e-214 | Homeo domain factors(Homeobox) |
| **POU2F3** | Core-like | 0.966920983173304 | 0.68621170785894 | 0 | 0 | Homeo domain factors(Homeobox) |
| **SCRT2** | Core-like | 0.603013073997812 | 0.641314195523166 | 8.36853388051562e-227 | 2.25416253036868e-226 | C2H2 zinc finger factors |
| **SNAI2** | Core-like | 1.09141916675551 | 0.701183326835115 | 0 | 0 | C2H2 zinc finger factors |
| **SOX2** | Core-like | 0.698352354059052 | 0.646266330250946 | 7.93488925055792e-243 | 2.24231468553713e-242 | High-mobility group (HMG) domain factors |
| **TCF3** | Core-like | 1.41227309294333 | 0.751077629554001 | 0 | 0 | Basic helix-loop-helix factors (bHLH) |
| **TCF4** | Core-like | 1.43262606455519 | 0.743289580530764 | 0 | 0 | Basic helix-loop-helix factors (bHLH) |
| **TWIST1** | Core-like | 0.799731749068282 | 0.634041247549593 | 2.8609842031081e-204 | 7.1299330731001e-204 | Basic helix-loop-helix factors (bHLH) |
| **YY2** | Core-like | 0.523476032793573 | 0.634912986781165 | 6.58044289759656e-207 | 1.66616814167145e-206 | C2H2 zinc finger factors |

**Note:** AUC = Area under curve, FDR = False discovery rate, TF = transcription factor
