## Supplemental Table 8 for "Identification of an epigenetically and phenotypically distinct peritumoral glioblastoma cell population linked to inferior patient outcome"

**Table S8. Differentially expressed genes in the Core subclusters**

| Gene | Subcluster | Average Expression | Log2 fold-change | AUC | p-val | FDR | RNA.pct_in | RNA.pct_out |
|---|---|---|---|---|---|---|---|---|
| POSTN | Acore1 | 1.58792653745804 | 0.525887124363736 | 0.673652844976374 | 4.80475220857406e-116 | 6.09353482380984e-116 | 83.4224598930481 | 96.1111111111111 |
| NEFL | Acore1 | 1.87156370900465 | 0.792538586565955 | 0.789133751143555 | 0 | 0 | 89.3048128342246 | 96.0493827160494 |
| VOPP1 | Acore1 | 1.16086468625444 | 0.427520548681358 | 0.621346636297617 | 1.16016279630187e-57 | 1.35533036951153e-57 | 80.672268907563 | 95.0617283950617 |
| PLD5 | Acore1 | 1.61875602217613 | 0.609710195855717 | 0.690560306614228 | 2.40230114226551e-139 | 3.11784703733357e-139 | 84.3391902215432 | 94.9074074074074 |
| MARCHF1 | Acore1 | 1.43340301591283 | 0.520808865363685 | 0.636407963858944 | 1.74254751858582e-72 | 2.08314108617551e-72 | 74.4843391902215 | 94.6296296296296 |
| AC092957.1 | Acore1 | 2.29946761460287 | 0.591251235821095 | 0.726914865272708 | 1.38190414500247e-196 | 1.91134736514865e-196 | 92.6279602750191 | 95.3703703703704 |
| KIF26B | Acore1 | 1.20117143107025 | 0.580748652852924 | 0.642310594271379 | 7.54879405462668e-79 | 9.08945701941803e-79 | 71.1993888464477 | 93.3641975308642 |
| KCNIP4 | Acore1 | 3.19607383482466 | 0.939112136821178 | 0.694954505371172 | 1.21351026895534e-145 | 1.58421706129939e-145 | 91.4438502673797 | 96.6975308641975 |
| LRRC4C | Acore1 | 2.36292638543207 | 0.988981105404098 | 0.705738465419838 | 5.58292098617699e-162 | 7.38481611928173e-162 | 90.2979373567609 | 96.5740740740741 |
| TGFBI | Acore1 | 1.22921135343472 | 0.603618419077017 | 0.760391378773732 | 1.67178149337993e-258 | 2.49892599907313e-258 | 81.9327731092437 | 94.4444444444444 |
| TMEFF2 | Acore1 | 1.97068835626507 | 0.837788270280142 | 0.670251299172868 | 1.10815849864805e-111 | 1.40184503307786e-111 | 81.2070282658518 | 94.7530864197531 |
| CACNA1A | Acore1 | 2.46074336023281 | 0.929503580751794 | 0.778253473106414 | 1.76146497266776e-294 | 2.76959901362856e-294 | 95.912910618793 | 97.9938271604938 |
| ADAMTSL1 | Acore1 | 1.23114222069326 | 0.540512287939944 | 0.678965719284347 | 1.85198211325394e-123 | 2.36222208323206e-123 | 75.2482811306341 | 92.3765432098765 |
| SDK1 | Acore1 | 2.82285948513724 | 0.718933183853554 | 0.718849560026031 | 5.72213863482604e-183 | 7.76409584101227e-183 | 98.4721161191749 | 98.6111111111111 |
| ANK3 | Acore1 | 1.69881223039806 | 0.848442849601506 | 0.728212328702525 | 5.33364771922496e-199 | 7.42330928215026e-199 | 83.7280366692131 | 94.320987654321 |
| CSMD2 | Acore1 | 1.45007485869688 | 0.821288834902478 | 0.667111533165455 | 7.24462934798626e-108 | 9.11847620891915e-108 | 75.7448433919022 | 92.7469135802469 |
| PLXNA4 | Acore1 | 1.52023692597945 | 0.424966228146415 | 0.629824033990701 | 9.93474810285672e-66 | 1.17155048382744e-65 | 82.5439266615737 | 94.0123456790123 |
| PPFIA2 | Acore1 | 1.62815144895063 | 0.690536807340945 | 0.771125706174726 | 7.95723184222369e-280 | 1.22986581796348e-279 | 88.6554621848739 | 94.9074074074074 |
| MIR31HG | Acore1 | 0.768706970225492 | 0.340953145359297 | 0.602312987484556 | 7.39556511641221e-42 | 8.36129464828967e-42 | 62.1084797555386 | 91.358024691358 |
| CXCL14 | Acore1 | 0.98126677123907 | 0.331299107943553 | 0.608851175150195 | 5.62578439774781e-47 | 6.44789042721812e-47 | 67.8762414056532 | 92.1913580246914 |
| COL6A2 | Acore1 | 2.03259258247236 | 0.662534463450857 | 0.78636287006385 | 0 | 0 | 94.7288006111535 | 98.3024691358025 |
| JAKMIP2-AS1 | Acore1 | 1.89988200732292 | 0.655763304751397 | 0.754840774693716 | 2.1656966086028e-247 | 3.18953845155052e-247 | 92.5897631779985 | 96.0493827160494 |
| TNC | Acore1 | 1.81606629479646 | 0.789967389420037 | 0.736699098831452 | 1.04472198536989e-213 | 1.48609101759587e-213 | 91.5966386554622 | 97.3765432098765 |
| COL6A1 | Acore1 | 2.16305468361899 | 0.692892840969863 | 0.780366810023673 | 6.06058933146621e-299 | 9.58954008143387e-299 | 93.8502673796791 | 98.4567901234568 |
| EYA4 | Acore1 | 1.56689267222746 | 0.43334894184245 | 0.699472137339784 | 2.18184727534509e-152 | 2.87463409136375e-152 | 90.7563025210084 | 95.9259259259259 |
| RUNX2 | Acore1 | 1.72515299737685 | 0.681988294858668 | 0.72704902668138 | 6.93413038792776e-197 | 9.60405870904122e-197 | 87.624140565317 | 95.1234567901235 |
| SEMA3A | Acore1 | 2.0912962609812 | 0.562873931759461 | 0.715466641201935 | 1.96395247961993e-177 | 2.63972107475797e-177 | 93.3155080213904 | 97.5308641975309 |
| PDE1C | Acore1 | 2.28040667846171 | 0.815080322428628 | 0.710759261617105 | 6.86091922951237e-170 | 9.15399496932938e-170 | 90.2597402597403 | 95.8024691358025 |
| CALCRL | Acore1 | 1.52160266650873 | 0.298055166850773 | 0.636005243848381 | 6.72209011668184e-72 | 8.02637625872458e-72 | 85.3323147440795 | 97.037037037037 |
| BNC2 | Acore1 | 1.75180468669169 | 0.706998856152247 | 0.766982205340048 | 2.1073184005274e-271 | 3.2026115509535e-271 | 89.3430099312452 | 94.3827160493827 |
| PTPRG | Acore1 | 2.03695986449486 | 0.336449132307258 | 0.610989799960388 | 1.79913800868814e-48 | 2.07035444037761e-48 | 91.2910618792972 | 97.1913580246914 |
| DCBLD2 | Acore1 | 2.46727936395824 | 0.693143144085179 | 0.73767801733488 | 1.96834831271627e-215 | 2.8039149753793e-215 | 97.5553857906799 | 98.8271604938272 |
| DOCK5 | Acore1 | 1.5192099859456 | 0.496318545990084 | 0.728091783851588 | 1.12966929306695e-198 | 1.57116730607364e-198 | 88.0061115355233 | 95.0617283950617 |
| HDAC9 | Acore1 | 1.58922938491815 | 0.627694825464155 | 0.69285608182667 | 1.37025500476292e-142 | 1.78302538030308e-142 | 88.4644766997708 | 96.6975308641975 |
| CALD1 | Acore1 | 2.99930306787665 | 0.320536108754324 | 0.613687175206783 | 9.24700059835112e-51 | 1.06778297902438e-50 | 99.2742551566081 | 99.7839506172839 |
| MLLT3 | Acore1 | 1.23022537559279 | 0.457276258639664 | 0.651029140612474 | 2.71575495483324e-88 | 3.30383814456598e-88 | 82.0473644003056 | 94.4135802469136 |

| | | | | | | | | |
|---|---|---|---|---|---|---|---|---|
| ZNF423 | Acore1 | 0.864810485395885 | 0.353852314335336 | 0.62932393496119 | 1.65246237545759e-65 | 1.94751016553634e-65 | 70.9320091673033 | 91.358024691358 |
| ANKS1B | Acore1 | 1.07560287238117 | 0.315442512757528 | 0.619212314555452 | 7.32715578727357e-56 | 8.52490492992852e-56 | 71.7341482047364 | 93.7037037037037 |
| DCLK1 | Acore1 | 1.28035061347841 | 0.361716282857471 | 0.630376477190203 | 2.54022992866664e-66 | 3.00086229021458e-66 | 79.2207792207792 | 93.6111111111111 |
| PCDH17 | Acore1 | 1.16899994828679 | 0.380862692538697 | 0.659166360146752 | 7.63941033488212e-98 | 9.45471576099273e-98 | 81.4362108479756 | 95.2777777777778 |
| SLC5A3 | Acore1 | 0.89232408074509 | 0.3246789930504 | 0.623652196568863 | 5.34077983432171e-60 | 6.25384055541184e-60 | 71.2757830404889 | 92.6851851851852 |
| ARL4C | Acore1 | 1.20503495314575 | 0.346558480856932 | 0.667850364051344 | 1.45200024222806e-108 | 1.82986798012357e-108 | 82.3911382734912 | 95.9876543209877 |
| HIVEP3 | Acore1 | 1.39428384861617 | 0.515369928380879 | 0.608487005913476 | 1.79394248672741e-46 | 2.05374068314529e-46 | 78.6860198624905 | 94.320987654321 |
| LOX | Acore1 | 0.775637562841209 | 0.263200775474104 | 0.602415966386555 | 7.22906485378044e-42 | 8.17767517395978e-42 | 64.0565317035905 | 92.1604938271605 |
| NRIP1 | Acore1 | 1.09293524686293 | 0.336597424794249 | 0.615619311697743 | 1.42593528398756e-52 | 1.65038806017078e-52 | 77.2727272727273 | 93.4259259259259 |
| PRICKLE1 | Acore1 | 1.79558277029933 | 0.332073072330044 | 0.662664164992596 | 5.3557930158497e-102 | 6.66557936011164e-102 | 90.9854851031322 | 96.2654320987654 |
| CAV1 | Acore1 | 1.37357031134624 | 0.382130922424836 | 0.657272420752813 | 1.46668285526037e-95 | 1.80625967396597e-95 | 81.3980137509549 | 96.1111111111111 |
| PALLD | Acore1 | 1.78317117922967 | 0.417013831473657 | 0.670947570947571 | 1.96175803004538e-112 | 2.48481067770156e-112 | 92.2459893048128 | 97.5617283950617 |
| NHS | Acore1 | 0.836602947951142 | 0.343228312694531 | 0.605405419743655 | 3.59591033017872e-44 | 4.08626173883945e-44 | 67.07410236822 | 92.3765432098765 |
| RNF220 | Acore1 | 1.43384095252159 | 0.308310575655707 | 0.611864148016109 | 3.13110464370513e-49 | 3.60726341440683e-49 | 86.9365928189458 | 95.3086419753086 |
| ELOVL6 | Acore1 | 1.39108536183823 | 0.486816736847261 | 0.693759077705156 | 5.83107353363596e-144 | 7.59748994610549e-144 | 85.5614973262032 | 95.5864197530864 |
| PTPN14 | Acore1 | 1.70841932397043 | 0.688382456766219 | 0.73634265153873 | 3.99547976092032e-213 | 5.6794310745136e-213 | 88.5026737967914 | 95.9876543209877 |
| UST | Acore1 | 1.72611326712212 | 0.366772917147115 | 0.651351281253242 | 1.4869602028791e-88 | 1.81115737256894e-88 | 92.5133689839572 | 96.7592592592593 |
| SLC4A4 | Acore2 | 0.623421425280701 | 0.415699204514956 | 0.721783562839306 | 5.71086860462946e-174 | 4.07919186044962e-173 | 81.1233352634626 | 44.1055434519487 |
| RGS6 | Acore2 | 1.06662141032723 | 0.483474220232614 | 0.667076311033682 | 6.94343407006445e-96 | 1.5193510000141e-95 | 83.0920671685003 | 53.0864197530864 |
| EPHA3 | Acore2 | 0.465804040826331 | 0.258773805015319 | 0.710089109739416 | 2.35859801819956e-160 | 1.2818467490215e-159 | 78.5755645628257 | 40.159767610748 |
| PDE4B | Acore2 | 1.39309989377798 | 0.298696336920663 | 0.60348142345145 | 1.65818167443394e-36 | 2.19772256386209e-36 | 89.5193977996526 | 67.6349552166546 |
| EFEMP1 | Acore2 | 0.543179240115386 | 0.290308324463489 | 0.706097652769315 | 6.15499190048046e-152 | 2.85614473340161e-151 | 80.9496236247829 | 42.2173807794723 |
| TRPM3 | Acore2 | 0.930256290279135 | 0.317108772389647 | 0.658328704246859 | 2.94529738319929e-84 | 5.69690016092707e-84 | 87.3769542559351 | 60.396998305495 |
| GRIA1 | Acore2 | 0.510742042379267 | 0.257058765195396 | 0.679862892135487 | 2.10001051785684e-112 | 5.51183862954551e-112 | 81.1812391430226 | 49.7700314693779 |
| NFIA | Acore2 | 1.06119465296191 | 0.400771233055943 | 0.674631919292841 | 3.11641868851148e-102 | 7.22229128276125e-102 | 87.7822814128547 | 59.5739530380053 |
| PTPRZ1 | Acore2 | 2.61995782389634 | 0.303571372275939 | 0.607241671393872 | 1.96830466160274e-38 | 2.63847809866319e-38 | 97.9154603358425 | 95.9331880900508 |
| NEAT1 | Acore2 | 1.48075841928217 | 0.506889082884495 | 0.65828609282254 | 4.5682527404743e-83 | 8.74306744588383e-83 | 90.4458598726115 | 66.8119099491648 |
| SPARCL1 | Acore2 | 1.1047664555513 | 0.386868528803146 | 0.653763745162938 | 2.20888437383965e-78 | 4.04187442605608e-78 | 90.040532715692 | 67.9980634229 |
| IGFBP7 | Acore2 | 1.40591611914588 | 0.491362402613876 | 0.669796503816736 | 7.39687321940837e-95 | 1.60452781332069e-94 | 91.7776491024899 | 69.7893972403776 |
| TENM1 | Acore2 | 0.923666403656457 | 0.325749858920393 | 0.655398538063706 | 2.38286453720712e-82 | 4.5301607171238e-82 | 85.5819339895773 | 55.5555555555556 |
| ANK2 | Acore2 | 1.32530380820402 | 0.365707263140963 | 0.637957640039152 | 9.4874922143898e-63 | 1.50118547696041e-62 | 93.341053850608 | 78.3103364802711 |
| AC002069.2 | Acore3 | 1.21402450805556 | 0.309617253515535 | 0.629486909386012 | 1.02873710384409e-51 | 1.31468000491258e-51 | 97.8189028420357 | 68.4234752589183 |
| LHFPL3 | Acore3 | 0.822615851380167 | 0.378030559602791 | 0.716712770108237 | 1.3900911429623e-145 | 2.45165986413103e-145 | 95.9682749504296 | 54.4994246260069 |
| GLCCI1 | Acore3 | 1.1194783209904 | 0.330832205672148 | 0.652631546923213 | 9.98166396977919e-71 | 1.35804951969785e-70 | 98.8103106411104 | 73.6248561565017 |
| RGS6 | Acore3 | 0.951890474452425 | 0.304980018740229 | 0.702435356941033 | 5.09176293735399e-129 | 8.36772873846177e-129 | 95.902181097158 | 50.1035673187572 |
| IL1RAPL1 | Acore3 | 0.85214757843928 | 0.31658368717604 | 0.684471671292222 | 1.65070522891915e-104 | 2.49539717145752e-104 | 97.2901520158625 | 60.0690448791715 |
| DGKB | Acore3 | 0.565924272643329 | 0.279723366478289 | 0.724290898138648 | 3.69430975177701e-158 | 6.93116276130771e-158 | 95.902181097158 | 49.6662830840046 |
| SOX6 | Acore3 | 1.57640658027253 | 0.573413229603901 | 0.691155136496357 | 6.81836149249215e-110 | 1.05221627970558e-109 | 98.4798413747522 | 73.3256616800921 |
| GRIA2 | Acore3 | 0.547944919702295 | 0.270418893693921 | 0.750860855325955 | 3.36071843049583e-200 | 8.27763160220649e-200 | 96.0343688037013 | 45.7537399309551 |
| CHL1 | Acore3 | 0.61084457807495 | 0.25490433304359 | 0.727182979577836 | 1.55184161479918e-162 | 2.99294429083738e-162 | 95.5717118307997 | 49.2059838895282 |

| | | | | | | | | |
|---|---|---|---|---|---|---|---|---|
| CSGALNACT1 | Acore3 | 0.658054553158502 | 0.289575506404605 | 0.710427008884261 | 9.83083482243569e-138 | 1.67333358679756e-137 | 96.3648380700595 | 53.4407364787112 |
| ETV1 | Acore3 | 1.46986484713816 | 0.511717133011502 | 0.697272658821096 | 1.01320991862776e-116 | 1.59435077675494e-116 | 98.6120290812954 | 75.2359033371692 |
| PDE4B | Acore3 | 1.79550162959333 | 0.826509536530626 | 0.749860700929497 | 4.6990267396764e-188 | 1.05833935578297e-187 | 98.8103106411104 | 65.4775604142693 |
| DPP6 | Acore3 | 1.08410776838143 | 0.417380310382227 | 0.694594146472801 | 5.80025530918548e-116 | 9.11273418568025e-116 | 96.4970257766028 | 60.8055235903337 |
| SLC35F1 | Acore3 | 1.18688412677323 | 0.263992609545259 | 0.624051393485078 | 2.37317162192298e-47 | 2.99832169541753e-47 | 98.2815598149372 | 73.4177215189873 |
| SOX2-OT | Acore3 | 3.27775603013052 | 0.562778916857422 | 0.65369163452609 | 3.84740909689993e-71 | 5.23813355602441e-71 | 99.9339061467284 | 95.9493670886076 |
| EFEMP1 | Acore3 | 0.525676495210793 | 0.252412568172051 | 0.742968990650268 | 5.2667182599551e-194 | 1.24361706256319e-193 | 94.1837409120952 | 39.5166858457998 |
| MAP3K1 | Acore3 | 0.55167003303965 | 0.318126942516905 | 0.758055654218864 | 1.15515694468444e-209 | 3.11363057866425e-209 | 96.2987442167878 | 48.2163406214039 |
| LINC00511 | Acore3 | 1.63150657925328 | 0.416641559062116 | 0.661277216178619 | 2.66373052503853e-78 | 3.70219669915014e-78 | 99.4051553205552 | 83.1300345224396 |
| DNM3 | Acore3 | 1.24917404456476 | 0.296109045157207 | 0.638488375011504 | 2.77976612016079e-58 | 3.63368120282456e-58 | 98.8103106411104 | 80.7364787111622 |
| LMO3 | Acore3 | 0.439630905865887 | 0.27254723077766 | 0.791582654964987 | 1.34379465294508e-286 | 7.97504245071264e-286 | 94.1176470588235 | 34.5684695051784 |
| TRPM3 | Acore3 | 0.963181812427625 | 0.345881253737052 | 0.65271787203652 | 1.67004706240153e-72 | 2.28773570191991e-72 | 97.2240581625909 | 58.2968929804373 |
| GRIN2B | Acore3 | 1.05858511461821 | 0.262906605007569 | 0.64045400164436 | 1.02272463833624e-60 | 1.3448055730917e-60 | 97.0918704560476 | 66.3521288837745 |
| PMP2 | Acore3 | 0.838117284960715 | 0.3355670610178 | 0.707880151840931 | 3.82142188842348e-134 | 6.39033760605934e-134 | 97.5545274289491 | 53.8780207134637 |
| ZEB2 | Acore3 | 0.834930132293823 | 0.289984692852044 | 0.682941929438537 | 4.12930343901276e-102 | 6.17696849515745e-102 | 98.0171844018506 | 63.4752589182969 |
| NOVA1 | Acore3 | 2.3675650757799 | 0.386236297093111 | 0.637976813150623 | 1.07564526174078e-57 | 1.40149219770786e-57 | 99.8017184401851 | 94.9827387802071 |
| REV3L | Acore3 | 1.88675661950174 | 0.575712290892364 | 0.705120714452497 | 2.62033011773221e-125 | 4.26069937842635e-125 | 99.4712491738268 | 88.0092059838895 |
| PPP2R2B | Acore3 | 0.971287027755319 | 0.259129310501423 | 0.651212848827614 | 2.16532087595557e-70 | 2.94201205972224e-70 | 98.0171844018506 | 63.2451093210587 |
| NFIA | Acore3 | 1.07542186257062 | 0.400213799227143 | 0.667122224951837 | 1.41230559949749e-86 | 1.99337417007408e-86 | 97.3562458691342 | 57.6294591484465 |
| PTPRZ1 | Acore3 | 2.80053662273472 | 0.532079158220293 | 0.670409272305915 | 4.98211427392844e-87 | 7.05182487463331e-87 | 99.6695307336418 | 95.4200230149597 |
| LSAMP | Acore3 | 2.80092417641631 | 0.317716050645665 | 0.616237943956367 | 1.84458115229567e-41 | 2.30428626145618e-41 | 99.9339061467284 | 97.6985040276179 |
| NEAT1 | Acore3 | 1.41135997067219 | 0.3883596523449 | 0.644277025274624 | 4.954878427595e-64 | 6.58893407924867e-64 | 97.8849966953073 | 65.385500575374 |
| IGFBP7 | Acore3 | 1.34023284686915 | 0.378606553786152 | 0.652018448475316 | 1.47753119176671e-70 | 2.00887993442109e-70 | 97.4223397224058 | 68.9067894131185 |
| TENM1 | Acore3 | 1.02738251542409 | 0.449537779041409 | 0.71503813896746 | 8.1656722922624e-144 | 1.43006520004595e-143 | 95.5717118307997 | 53.5558112773303 |
| MAP2 | Acore3 | 2.39999135870002 | 0.713657997932173 | 0.778769422199777 | 1.28245050051123e-229 | 4.17057073336985e-229 | 99.7356245869134 | 90.9090909090909 |
| PLCB1 | Acore3 | 1.13652158903232 | 0.387026454076866 | 0.67552694446367 | 3.34179844092275e-94 | 4.8572651757598e-94 | 97.1579643093192 | 63.4752589182969 |
