## Supplemental Table 9 for "Identification of an epigenetically and phenotypically distinct peritumoral glioblastoma cell population linked to inferior patient outcome"

**Table S9. Differentially expressed genes in the Edge subclusters**

| Gene | Subcluster | Average Expression | Log2 fold-change | AUC | p-val | FDR | RNA.pct_in | RNA.pct_out |
|---|---|---|---|---|---|---|---|---|
| SYT1 | Aedge1 | 1.01293231629495 | 0.550992418600922 | 0.649196884099824 | 8.16052528591999e-110 | 1.03889564429281e-109 | 65.0053345526597 | 63.8569078947368 |
| MYBPC1 | Aedge1 | 1.28040186639247 | 0.570549374586586 | 0.656508047704137 | 2.44301260947485e-118 | 3.14213840446926e-118 | 70.4923030025911 | 71.3815789473684 |
| SPP1 | Aedge1 | 1.89587487067688 | 0.334609165576133 | 0.603174773281512 | 3.21370013472697e-51 | 3.78304901086165e-51 | 94.9550373418686 | 97.2450657894737 |
| COL1A2 | Aedge1 | 2.12565564929006 | 0.434551596435387 | 0.664729977127604 | 1.12766945593707e-127 | 1.46831960408473e-127 | 94.5739978661789 | 94.3256578947368 |
| LAMA2 | Aedge1 | 1.68876510807471 | 0.573286208239539 | 0.666881095378192 | 1.61667500139496e-131 | 2.11468280103984e-131 | 85.6881572930956 | 78.3305921052632 |
| SEMA3C | Aedge1 | 1.15065080754773 | 0.360979570005568 | 0.617808381264088 | 1.57009271836794e-67 | 1.88713067111531e-67 | 70.7361682670325 | 78.7006578947368 |
| PRSS12 | Aedge1 | 0.970678864840699 | 0.331510283497793 | 0.610812233272367 | 3.08293028176549e-60 | 3.67234101461048e-60 | 68.0993750952599 | 78.5361842105263 |
| RNF150 | Aedge1 | 1.654012542255 | 0.49634152467848 | 0.678789299709608 | 1.9656130000506e-150 | 2.6278248663778e-150 | 85.8862978204542 | 85.8963815789474 |
| EFEMP1 | Aedge1 | 2.13173718027522 | 0.356691279770846 | 0.648243752707386 | 9.23290470045654e-104 | 1.16650722684227e-103 | 95.71711629324 | 96.8338815789474 |
| LMO3 | Aedge1 | 1.68549302111671 | 0.291879605740529 | 0.638114683105512 | 2.19294115679331e-90 | 2.72584357587733e-90 | 89.8948331047096 | 93.0098684210526 |
| PAWR | Aedge1 | 0.842390498158393 | 0.317760796248213 | 0.608160530677689 | 2.3592868887775e-58 | 2.80533518285078e-58 | 64.0146319158665 | 70.8881578947368 |
| IGFBP7 | Aedge1 | 3.11488237077033 | 0.27131731987182 | 0.661893081827225 | 2.31263629532386e-123 | 2.99370394216681e-123 | 99.9237921048621 | 99.9177631578947 |
| ADGRL3 | Aedge1 | 2.28836694814752 | 0.410517429505395 | 0.639584787299754 | 3.17547044403124e-92 | 3.95943945639805e-92 | 95.3208352385307 | 91.1595394736842 |
| CAV1 | Aedge1 | 1.88164906722548 | 0.279143569647058 | 0.626699291918353 | 2.58611506542461e-76 | 3.15572308166517e-76 | 94.31489102271 | 95.8881578947368 |
| FBN1 | Aedge1 | 1.27125108848726 | 0.260315687844769 | 0.635245574928405 | 6.31588432761073e-87 | 7.83122669263575e-87 | 84.7584209724127 | 88.4457236842105 |
| COL4A5 | Aedge1 | 1.81050408145568 | 0.480069412462419 | 0.667264610357054 | 1.29970352769364e-131 | 1.70118262787126e-131 | 94.0710257582686 | 87.0065789473684 |
| SDK1 | Aedge2 | 1.17729230870485 | 0.272200743160391 | 0.613578753375766 | 3.78224855200237e-44 | 4.2931311600481e-44 | 99.4486560992419 | 62.9408644921771 |
| SNTG1 | Aedge2 | 0.991853652800847 | 0.49403242091039 | 0.783476167735891 | 0 | 0 | 98.6905582356995 | 32.3919384778573 |
| PDE4B | Aedge2 | 1.41247383775151 | 0.253031178521774 | 0.604110251600913 | 4.50996507061327e-37 | 5.03625356852403e-37 | 99.4486560992419 | 66.6666666666667 |
| SLC35F1 | Aedge2 | 0.973763483718562 | 0.272852286755432 | 0.667173180065285 | 1.91177793252728e-97 | 2.29092622232149e-97 | 99.0351481736733 | 50.994431185362 |
| GFAP | Aedge2 | 1.79823936979243 | 0.294767096096451 | 0.600288465000317 | 5.68499652462982e-34 | 6.31315549653506e-34 | 99.6554100620262 | 79.236276849642 |
| PTPRZ1 | Aedge2 | 1.79704586182029 | 0.391933517323584 | 0.634457376390353 | 1.19385825331786e-59 | 1.38018295181256e-59 | 99.8621640248105 | 79.5544948289578 |
| PTN | Aedge2 | 1.44370048396218 | 0.275196449412268 | 0.61583594996894 | 1.01346992952744e-44 | 1.151670374463e-44 | 99.6554100620262 | 79.9787854680456 |
| RGS6 | Aedge3 | 1.31532602507441 | 0.629165572678451 | 0.621214775695784 | 5.98931438448653e-39 | 2.07962305016894e-38 | 61.5698267074414 | 53.3325012481278 |
| AC092957.1 | Aedge3 | 1.63821668358154 | 0.937267838209012 | 0.752678258351514 | 7.911764838008e-161 | 3.955882419004e-158 | 83.2823649337411 | 53.0579131303045 |
| KCNMB2 | Aedge3 | 1.75327353867466 | 0.774489661107891 | 0.717669609245663 | 2.42840990069841e-112 | 4.85681980139682e-110 | 86.5443425076453 | 70.8686969545682 |
| TMEFF2 | Aedge3 | 1.65329366548255 | 0.512982467221156 | 0.629960640079636 | 3.95791848584184e-41 | 1.78687064823559e-40 | 78.7971457696228 | 73.0654018971543 |
| AGBL4 | Aedge3 | 0.473034248552717 | 0.358504603154074 | 0.603815479634778 | 2.07164909644036e-41 | 9.81824216322447e-41 | 40.2650356778797 | 26.1607588617074 |
| TENM2 | Aedge3 | 1.49122935328897 | 0.467738129948393 | 0.621697741359419 | 1.1846323692832e-36 | 3.39923204959311e-36 | 75.6371049949032 | 68.5971043434848 |
| GRIA4 | Aedge3 | 1.02114114857551 | 0.825752082161537 | 0.756132429795673 | 1.38105708276698e-211 | 1.38105708276698e-208 | 68.3995922528033 | 29.9051422865701 |
| PDGFRA | Aedge3 | 0.814073883542006 | 0.342166395510196 | 0.626230506431993 | 1.63951277794941e-42 | 1.00276010883756e-41 | 64.2201834862385 | 51.3979031452821 |
| THSD4 | Aedge3 | 1.05123097403439 | 0.357605082616786 | 0.629388422463145 | 4.75711352212123e-42 | 2.58538778376154e-41 | 72.0693170234455 | 62.3315027458812 |
| IGFBP5 | Aedge3 | 2.14947781730334 | 0.29814032270178 | 0.600263468202386 | 9.71538043264304e-25 | 1.77287964099325e-24 | 95.7186544342508 | 92.873190214678 |
| SNTG1 | Aedge3 | 1.33767133430227 | 0.853211548707997 | 0.696635602152327 | 4.0592283429763e-110 | 7.38041516904781e-108 | 66.5647298674822 | 40.2146779830255 |
| GFAP | Aedge3 | 1.97549453428949 | 0.476433990649213 | 0.619814162548227 | 8.79851264628185e-35 | 2.26765789852625e-34 | 87.8695208970438 | 81.8771842236645 |
| TMEM108 | Aedge3 | 0.720684722805581 | 0.372480921621664 | 0.607618897850981 | 2.9545839537022e-34 | 7.42357777312111e-34 | 53.7206931702345 | 41.762356465302 |
| KCNMB2-AS1 | Aedge3 | 0.681206617089807 | 0.339999233052017 | 0.603953397121443 | 5.44661754935891e-31 | 1.21170579518552e-30 | 55.1478083588175 | 45.282076884673 |
| CD74 | Aedge3 | 1.52396470734537 | 0.478559241003389 | 0.688488546487099 | 1.13421640753983e-83 | 1.13421640753983e-81 | 89.1946992864424 | 78.8317523714428 |
| HLA-DRA | Aedge3 | 1.70319761159877 | 0.471523208763235 | 0.681122302275435 | 4.68034469325625e-77 | 4.06986495065761e-75 | 91.6411824668705 | 84.0614078881678 |
| DOCK5 | Aedge3 | 1.13883263743562 | 0.323836478817552 | 0.615144752290524 | 1.6384712833003e-33 | 3.97205159587951e-33 | 75.1274209989806 | 63.2176734897653 |
| PMP2 | Aedge3 | 1.13457112343255 | 0.582448753292438 | 0.668410737614272 | 1.39831985527665e-74 | 1.07563065790511e-72 | 69.7247706422018 | 50.0748876684973 |
| SLC8A1 | Aedge3 | 0.941440516159159 | 0.302328406688044 | 0.601380485337234 | 4.50405648808097e-27 | 8.78840290357263e-27 | 67.1763506625892 | 57.3889166250624 |

| | | | | | | | | |
|---|---|---|---|---|---|---|---|---|
| TRIM9 | Aedge3 | 0.67406049376349 | 0.310480087562593 | 0.607296114950917 | 3.08344895054198e-33 | 7.3943619917074e-33 | 56.5749235474006 | 43.971542685971 |
| HLA-DRB1 | Aedge3 | 1.41024760517217 | 0.4364279807889 | 0.67515132754487 | 1.04903971208682e-72 | 7.23475663508149e-71 | 84.8114169215087 | 77.0219670494259 |
| ANKS1B | Aedge3 | 1.21727184917434 | 0.338544123900582 | 0.604025282158312 | 3.5381892212141e-27 | 6.93762592394922e-27 | 75.6371049949032 | 69.4583125312032 |
| SLC1A3 | Aedge3 | 0.834213734192533 | 0.407630702481416 | 0.609450566759443 | 3.07869934040998e-34 | 7.72571980027599e-34 | 56.5749235474006 | 44.882675986021 |
| GRIA1 | Aedge3 | 1.54340425891766 | 0.489001485303181 | 0.636563566983877 | 1.65611538031461e-45 | 2.49039906814227e-44 | 77.8797145769623 | 70.1572641038442 |
| RTN1 | Aedge3 | 0.823779168180036 | 0.313243910172494 | 0.615211802581551 | 1.88974143419672e-35 | 5.07313136697105e-35 | 63.710499490316 | 52.608587119321 |
| PTPRM | Aedge3 | 0.906343595422744 | 0.280586525514546 | 0.617245385744014 | 6.73079380208167e-35 | 1.74599061013792e-34 | 70.6422018348624 | 62.0069895157264 |
| CDH6 | Aedge3 | 1.24406289793392 | 0.51588075351811 | 0.662523734785182 | 1.07075611978477e-64 | 5.35378059892387e-63 | 77.5739041794088 | 64.4283574638043 |
| AL390957.1 | Aedge3 | 1.56203567607283 | 0.500874285190654 | 0.700527381710309 | 2.55902286553599e-94 | 3.41203048738132e-92 | 92.1508664627931 | 79.8676984523215 |
| FHIT | Aedge3 | 1.0206417452012 | 0.315053701460708 | 0.608752645751047 | 1.62223050308892e-30 | 3.55362651279062e-30 | 71.8654434250764 | 59.6854717923115 |
| HLA-DRB5 | Aedge3 | 0.907048771311579 | 0.331268293228434 | 0.648128787959752 | 8.29932440220604e-55 | 2.8618360007607e-53 | 72.986748216106 | 61.0209685471792 |
| ZFPM2 | Aedge3 | 1.08099349504268 | 0.366058050095165 | 0.653996451805472 | 4.32138852561866e-58 | 1.72855541024747e-56 | 78.0835881753313 | 65.3894158761857 |
| HLA-DPA1 | Aedge3 | 0.974719753918962 | 0.387827811952725 | 0.670302395540227 | 1.24353716670421e-71 | 8.02282043034974e-70 | 75.7390417940877 | 61.4952571143285 |
| NR2F2-AS1 | Aedge3 | 0.842948267805655 | 0.253945406386433 | 0.602671044401797 | 2.99199998183993e-27 | 5.8781924987032e-27 | 66.6666666666667 | 61.5202196704943 |
| HLA-DPB1 | Aedge3 | 0.685028881373258 | 0.308415694017631 | 0.61842983486035 | 2.60114271121115e-38 | 8.57048669262322e-38 | 58.8175331294597 | 49.3884173739391 |
