## Supplemental Table 10 for "Identification of an epigenetically and phenotypically distinct peritumoral glioblastoma cell population linked to inferior patient outcome"

**Table S10. The Edge-unique gene signature. DEGs of Edge profiles of the Edge-like cluster and Core profiles of the Core-like cluster in Figure 5**

| Gene | Cluster | LogFC | AUC | p-val | p-adj | RNA.pct_in | RNA.pct_out |
|---|---|---|---|---|---|---|---|
| S100A10 | edge | 0,90139862 | 0,815048958 | 0 | 0 | 96,41909814 | 72,13871308 |
| LTBP1 | edge | 1,31092909 | 0,848209374 | 0 | 0 | 88,03713528 | 41,44251055 |
| EFEMP1 | edge | 1,90128101 | 0,952471112 | 0 | 0 | 93,85941645 | 14,71518987 |
| PROS1 | edge | 0,93693226 | 0,836347969 | 0 | 0 | 85,39787798 | 38,10654008 |
| IGFBP7 | edge | 2,46103441 | 0,974343157 | 0 | 0 | 98,92572944 | 42,40506329 |
| SPARCL1 | edge | 1,14561255 | 0,836874984 | 0 | 0 | 94,11140584 | 46,63765823 |
| SPP1 | edge | 1,25431965 | 0,853074233 | 0 | 0 | 94,13793103 | 50,03955696 |
| ANK2 | edge | 1,6589459 | 0,906958064 | 0 | 0 | 98,63395225 | 73,95833333 |
| PRSS12 | edge | 0,86258423 | 0,800947583 | 0 | 0 | 64,15119363 | 5,814873418 |
| RNF150 | edge | 1,34172799 | 0,86527732 | 0 | 0 | 81,90981432 | 21,11023207 |
| PALLD | edge | 1,24101986 | 0,825149335 | 0 | 0 | 96,36604775 | 69,54113924 |
| PDE4D | edge | 1,30252268 | 0,806594987 | 0 | 0 | 97,37400531 | 75,23734177 |
| PAM | edge | 1,0739405 | 0,840033735 | 0 | 0 | 97,24137931 | 67,12816456 |
| CD74 | edge | 1,01462054 | 0,844144844 | 0 | 0 | 74,46949602 | 10,79905063 |
| SPARC | edge | 1,32561266 | 0,907748487 | 0 | 0 | 99,53580902 | 84,13765823 |
| HLA-B | edge | 0,92801648 | 0,832232515 | 0 | 0 | 95,26525199 | 64,34599156 |
| HLA-DRA | edge | 1,02477483 | 0,828804595 | 0 | 0 | 80,46419098 | 23,08808017 |
| HLA-DRB1 | edge | 0,92127505 | 0,825275937 | 0 | 0 | 72,04244032 | 11,03639241 |
| COL21A1 | edge | 2,08571699 | 0,92712686 | 0 | 0 | 95,17241379 | 27,57120253 |
| AL096854.1 | edge | 0,97254405 | 0,830209557 | 0 | 0 | 75,0132626 | 13,31751055 |
| LAMA2 | edge | 1,25303708 | 0,828894157 | 0 | 0 | 79,9204244 | 26,10759494 |
| AHR | edge | 1,51230373 | 0,901508656 | 0 | 0 | 93,63395225 | 41,74578059 |
| COL1A2 | edge | 1,75380426 | 0,913985502 | 0 | 0 | 92,25464191 | 22,61339662 |
| PEG10 | edge | 0,82195529 | 0,8012948 | 0 | 0 | 77,89124668 | 29,07436709 |
| CAV1 | edge | 1,13001898 | 0,817381455 | 0 | 0 | 92,79840849 | 47,58702532 |
| CALD1 | edge | 1,3045023 | 0,860254766 | 0 | 0 | 99,76127321 | 95,95200422 |
| CLU | edge | 1,28605729 | 0,886890828 | 0 | 0 | 99,86737401 | 91,99630802 |
| PRUNE2 | edge | 1,03519703 | 0,809365653 | 0 | 0 | 95,46419098 | 67,23364979 |
| NTRK2 | edge | 1,19153702 | 0,86481361 | 0 | 0 | 81,89655172 | 16,17879747 |
| NRP1 | edge | 1,18175896 | 0,870271771 | 0 | 0 | 87,5198939 | 27,99314346 |
| BICC1 | edge | 1,18297932 | 0,858638151 | 0 | 0 | 80,13262599 | 17,90611814 |
| NEAT1 | edge | 1,6932124 | 0,863133209 | 0 | 0 | 99,77453581 | 73,5100211 |
| NNMT | edge | 1,09509393 | 0,87627621 | 0 | 0 | 78,80636605 | 7,357594937 |
| GAPDH | edge | 0,71663451 | 0,802060818 | 0 | 0 | 99,65517241 | 95,20042194 |
| C1R | edge | 0,9381024 | 0,86117054 | 0 | 0 | 77,5066313 | 12,36814346 |
| EPS8 | edge | 1,00815102 | 0,841609237 | 0 | 0 | 91,35278515 | 56,98839662 |
| LMO3 | edge | 1,5282356 | 0,917422061 | 0 | 0 | 87,26790451 | 10,06065401 |
| PRICKLE1 | edge | 1,39238832 | 0,833770786 | 0 | 0 | 94,69496021 | 39,04272152 |
| MYBPC1 | edge | 1,09874069 | 0,803063015 | 0 | 0 | 63,63395225 | 4,931434599 |
| CHPT1 | edge | 1,14798332 | 0,854433799 | 0 | 0 | 91,24668435 | 52,99314346 |
| ITGBL1 | edge | 1,33960581 | 0,882132215 | 0 | 0 | 81,92307692 | 11,41877637 |
| GABRG3 | edge | 1,45099899 | 0,889695079 | 0 | 0 | 81,83023873 | 7,898206751 |
| THBS1 | edge | 0,93929801 | 0,804401709 | 0 | 0 | 69,29708223 | 14,14820675 |
| B2M | edge | 1,04052497 | 0,860431802 | 0 | 0 | 99,61538462 | 89,01635021 |
| FBN1 | edge | 1,00943191 | 0,857616683 | 0 | 0 | 82,04244032 | 23,65506329 |
| ANXA2 | edge | 1,36331414 | 0,924120688 | 0 | 0 | 99,01856764 | 77,00421941 |
| PKM | edge | 1,03793575 | 0,87166925 | 0 | 0 | 98,28912467 | 80,12921941 |
| PLCG2 | edge | 0,91380309 | 0,824729327 | 0 | 0 | 98,67374005 | 88,13291139 |
| MAP3K14 | edge | 0,92238805 | 0,814711832 | 0 | 0 | 80,29177719 | 33,86075949 |
| LGALS3BP | edge | 0,76366342 | 0,815267991 | 0 | 0 | 85,0795756 | 47,13871308 |
| COL4A5 | edge | 1,3150492 | 0,87139702 | 0 | 0 | 90,0928382 | 31,4214135 |
| DANT2 | edge | 0,96325256 | 0,848115509 | 0 | 0 | 82,86472149 | 21,45305907 |
