## Supplemental Table 11 for "Identification of an epigenetically and phenotypically distinct peritumoral glioblastoma cell population linked to inferior patient outcome"

**Table S11. The Edge-unique gene signature mapped to previous alias when required for survival analyses**

| Original gene symbol | TCGA | GLASS | Spitzer |
|---|---|---|---|
| S100A10 | S100A10 | S100A10 | S100A10 |
| LTBP1 | LTBP1 | LTBP1 | LTBP1 |
| EFEMP1 | DHRD | EFEMP1 | EFEMP1 |
| PROS1 | PROS1 | PROS1 | PROS1 |
| IGFBP7 | IGFBP7 | IGFBP7 | IGFBP7 |
| SPARCL1 | SPARCL1 | SPARCL1 | SPARCL1 |
| SPP1 | SPP1 | SPP1 | SPP1 |
| ANK2 | LQT4 | ANK2 | ANK2 |
| PRSS12 | PRSS12 | PRSS12 | PRSS12 |
| RNF150 | RNF150 | RNF150 | RNF150 |
| PALLD | PALLD | PALLD | PALLD |
| PDE4D | PDE4D | PDE4D | PDE4D |
| PAM | PAM | PAM | PAM |
| CD74 | CD74 | CD74 | CD74 |
| SPARC | SPARC | SPARC | SPARC |
| HLA-B | AS | HLA-B | HLA-B |
| HLA-DRA | HLA-DRA | HLA-DRA | HLA-DRA |
| HLA-DRB1 | HLA-DRB1 | HLA-DRB1 | HLA-DRB1 |
| COL21A1 | COL21A1 | COL21A1 | COL21A1 |
| AL096854.1 | - | - | - |
| LAMA2 | LAMA2 | LAMA2 | LAMA2 |
| AHR | AHR | AHR | AHR |
| COL1A2 | COL1A2 | COL1A2 | COL1A2 |
| PEG10 | PEG10 | PEG10 | PEG10 |
| CAV1 | CAV1 | CAV1 | CAV1 |
| CALD1 | CALD1 | CALD1 | CALD1 |
| CLU | CLU | CLU | CLU |
| PRUNE2 | KIAA0367 | PRUNE2 | PRUNE2 |
| NTRK2 | NTRK2 | NTRK2 | NTRK2 |
| NRP1 | NRP1 | NRP1 | NRP1 |
| BICC1 | BICC1 | BICC1 | BICC1 |
| NEAT1 | NEAT1 | - | NEAT1 |
| NNMT | NNMT | NNMT | NNMT |
| GAPDH | GAPDH | GAPDH | GAPDH |
| C1R | C1R | C1R | C1R |
| EPS8 | EPS8 | EPS8 | EPS8 |
| LMO3 | LMO3 | LMO3 | LMO3 |
| PRICKLE1 | PRICKLE1 | PRICKLE1 | PRICKLE1 |
| MYBPC1 | MYBPC1 | MYBPC1 | MYBPC1 |
| CHPT1 | CHPT1 | CHPT1 | CHPT1 |
| ITGBL1 | ITGBL1 | ITGBL1 | ITGBL1 |
| GABRG3 | GABRG3 | GABRG3 | GABRG3 |
| THBS1 | THBS1 | THBS1 | THBS1 |
| B2M | B2M | B2M | B2M |
| FBN1 | FBN1 | FBN1 | FBN1 |
| ANXA2 | ANXA2 | ANXA2 | ANXA2 |
| PKM | PKM | PKM | PKM |
| PLCG2 | PLCG2 | PLCG2 | PLCG2 |
| MAP3K14 | MAP3K14 | - | MAP3K14 |
| LGALS3BP | LGALS3BP | LGALS3BP | LGALS3BP |
| COL4A5 | COL4A5 | COL4A5 | COL4A5 |
| DANT2 | - | - | DANT2 |
