## Supplemental Table 12 for "Identification of an epigenetically and phenotypically distinct peritumoral glioblastoma cell population linked to inferior patient outcome"

**Table S12. The OSM-activation gene signature. DEGs of Edge cells cultured in absence or presence of OSM for 7 days**

| Gene | baseMean | log2FoldChange | pvalue | padj |
|---|---|---|---|---|
| ACP3 | 365,4579691 | 3,879203806 | 2,47E-06 | 0,002747679 |
| PLXDC1 | 314,9121874 | 3,86258944 | 8,52E-07 | 0,001376547 |
| SOCS3 | 8140,88812 | 3,618834262 | 4,43E-09 | 1,57E-05 |
| RCAN2 | 1954,930308 | 3,028791696 | 2,58E-08 | 7,63E-05 |
| CEMIP | 11529,49814 | 2,955700475 | 9,97E-06 | 0,008473679 |
| NAMPT | 20524,99333 | 2,823734218 | 4,81E-13 | 4,27E-09 |
| PCOLCE2 | 1343,482212 | 2,817616643 | 1,00E-05 | 0,008473679 |
| SOD2 | 17428,26169 | 2,769255911 | 1,84E-06 | 0,002182391 |
| CEBPD | 911,7421348 | 2,724722726 | 6,86E-08 | 0,000174074 |
| EFNA1 | 361,4562859 | 2,621602156 | 5,35E-15 | 9,50E-11 |
| GCLM | 4990,950669 | 2,076078043 | 6,29E-05 | 0,041376311 |
| CSF1 | 11291,16185 | 1,991784175 | 2,05E-05 | 0,016566157 |
| OSMR | 8194,796231 | 1,887218442 | 5,96E-07 | 0,001059555 |
| TEAD4 | 1380,983098 | 1,831011364 | 1,78E-10 | 7,91E-07 |
| SRPX | 8554,314696 | 1,703394771 | 3,93E-05 | 0,029098036 |
| ITPKC | 1330,36483 | 1,513133169 | 3,90E-07 | 0,0007691 |
| ARID5A | 803,4797706 | 1,41472138 | 2,79E-06 | 0,002754049 |
| RALGDS | 2684,963422 | 1,390391134 | 4,26E-05 | 0,029300945 |
| SPRED3 | 2481,345035 | 1,354808736 | 1,70E-06 | 0,002155763 |
| SEMA4B | 2145,51005 | 1,330403108 | 4,19E-06 | 0,003916078 |
| VWA5A | 2053,10496 | 1,265649603 | 2,63E-06 | 0,002748881 |
| HECTD2 | 942,2616305 | 0,735624564 | 2,52E-05 | 0,019497344 |
