## Supplemental Table 13 for "Identification of an epigenetically and phenotypically distinct peritumoral glioblastoma cell population linked to inferior patient outcome"

**Table S13. The OSM-activation Edge gene signature mapped to previous alias when required for survival analyses**

| Original gene symbol | TCGA | GLASS | Spitzer |
|---|---|---|---|
| ACP3 | ACPP | ACPP | ACPP |
| PLXDC1 | PLXDC1 | PLXDC1 | PLXDC1 |
| SOCS3 | SOCS3 | SOCS3 | SOCS3 |
| RCAN2 | RCAN2 | RCAN2 | RCAN2 |
| CEMIP | CEMIP | KIAA1199 | CEMIP |
| NAMPT | NAMPT | NAMPT | NAMPT |
| PCOLCE2 | PCOLCE2 | PCOLCE2 | PCOLCE2 |
| SOD2 | SOD2 | SOD2 | SOD2 |
| CEBPD | CEBPD | CEBPD | CEBPD |
| EFNA1 | EFNA1 | EFNA1 | EFNA1 |
| GCLM | GCLM | GCLM | GCLM |
| CSF1 | CSF1 | CSF1 | CSF1 |
| OSMR | OSMR | OSMR | OSMR |
| TEAD4 | TEAD4 | TEAD4 | TEAD4 |
| SRPX | SRPX | SRPX | SRPX |
| ITPKC | ITPKC | ITPKC | ITPKC |
| ARID5A | ARID5A | ARID5A | ARID5A |
| RALGDS | RALGDS | RALGDS | RALGDS |
| SPRED3 | SPRED3 | SPRED3 | SPRED3 |
| SEMA4B | SEMA4B | SEMA4B | SEMA4B |
| VWA5A | VWA5A | VWA5A | VWA5A |
| HECTD2 | HECTD2 | HECTD2 | HECTD2 |
